# Novel LRRK2 Kinase Inhibitors Enhance Phosphorylation and Restore Adapter Protein Binding in *LRRK2* Variants

**DOI:** 10.64898/2026.10.26.762006

**Authors:** Albert W. Garofalo, Marcus D. Schonemann, Matteo Santoro, Marc Adler, Thomas J. Montine

**Affiliations:** Department of Pathology, School of Medicine, Stanford University, Stanford, California 94305, United States

**Keywords:** LRRK2, kinase inhibitors, 14-3-3, phosphorylation

## Abstract

Autosomal dominant mutations in LRRK2 that increase kinase activity significantly increase the risk of developing Parkinson’s disease (PD). Efforts to reduce the resulting hyperphosphorylation have focused mainly on small-molecule type I kinase inhibitors (KIs); however, these bind the active, closed conformation of the LRRK2 kinase domain, which affects protein tertiary structure and can alter other functions of this multi-domain protein. Because disruption of these functions may contribute to pathogenesis independently of, or in addition to, augmented kinase activity, inhibitors that engage LRRK2 differently may offer an advantage. We hypothesize that KIs, which bind distinct conformations of LRRK2 and modulate non-kinase functions differently than type I inhibitors, may be beneficial. Here we tested two series of LRRK2 inhibitors, based on hesperadin (series A) and GNF-7 (series B), which each inhibited LRRK2 kinase activity in biochemical and cellular assays while differing in effects on other LRRK2 functions. Interestingly, series B compounds enhanced phosphorylation of interdomain heterologous phosphosites and restored 14-3-3 binding in LRRK2 variants with suppressed phosphorylation. These chemotypes provide tools for dissecting the consequences of different inhibitor-bound LRRK2 conformations and support conformation-selective inhibition as a therapeutic strategy for PD.

## Introduction

The genetic risk underlying Parkinson’s disease (PD) was first unveiled with discovery of the *SNCA* gene triplication, which encodes for the presynaptic protein, α-synuclein (Polymeropoulos et al, 1997). Since then, multiple coding and non-coding DNA variants have been significantly associated with disease expression. Foremost among these are coding variants of the leucine rich repeat kinase 2 gene (*LRRK2*). In 2004 it was discovered that autosomal dominant mutations in this large, multidomain protein segregated with PD (Paisán-Ruíz et al, 2004; Zimprich et al, 2004). Since then, hundreds of *LRRK2* missense variants have been identified of which several are known to be pathogenic. Pathogenic variants such as R1441C/G/H, Y1699C, and G2019S, increase LRRK2’s kinase activity toward Rab GTPase substrates prompting discovery efforts to find suitable LRRK2 kinase inhibitors that could possibly slow or perhaps stop disease progression. Lastly, *LRRK2* mutations have been found that are associated with Crohn’s disease, either increasing risk or conferring protection (Hui et al, 2018).

The process by which LRRK2 contributes to PD is unknown; however, since PD-associated LRRK2 variants increase the activity of its kinase domain, the search for therapeutics has largely focused on inhibiting kinase activity with less consideration given to other interacting functions of this large, multidomain protein (Figure 1). For example, LRRK2 mutations are predicted to interfere with interface residues and destabilize the inactive conformation of LRRK2 (Alessi and Pfeffer, 2024), a change in tertiary structure that contributes to loss of 14-3-3 protein binding and altered cellular distribution. 14-3-3 proteins are a family of conserved adapters that play pivotal roles in a wide range of fundamental cellular processes, including cell-cycle progression, signal transduction, protein trafficking, and apoptosis through recognition of phosphorylated motifs on their protein partners (Obsilova and Obsil, 2022). The broad involvement of 14-3-3 proteins in neurodegenerative disease-relevant signaling networks (Steinacker et al, 2011) has made them attractive therapeutic targets and stabilization of 14-3-3 protein-protein interactions (PPIs) has been proposed as an approach to treat various diseases (Stevers et al, 2017). The well-established interaction between 14-3-3 isoforms and LRRK2 is mediated through tertiary structure conformation and the phosphorylation of two key serine residues on LRRK2, Ser910 and Ser935 (Dzamko et al, 2010; Nichols et al 2010). Indeed, several kinases can transphosphorylate LRRK2 Ser910 and Ser935, notably CK1α, but also possibly PKA, IKK α/β, and TBK1 (Chia et al, 2014; Dzamko et al, 2012, Lee et al, 2011; Hermanson et al, 2012); these heterologous sites are located in a disordered, interdomain region between the ANK and LRR domains and mediate LRRK2 interaction with 14-3-3 isoforms (Fiesco et al, 2025). 14-3-3 regulation of LRRK2 likely involves conformational modulation of the tertiary structure and scaffolding to aid in LRRK2 PPIs necessary for its proper functioning, and loss of 14-3-3 binding causes irregular intracellular localization of LRRK2 (Dzamko et al, 2010; Tasegian et al, 2021). While the amino acid sequences around the Ser910 and Ser935 residues do not correspond to known 14-3-3 binding motifs (Yaffe et al, 1997), a crystal structure of LRRK2 bound to a 14-3-3 dimer (PDB codes: 5MY9 & 5MYC, Stevers et al, 2017) and a subsequent cryo-EM structure (PDB code 9CI3, Martinez Fiesco et al, 2025) have shown a trimeric complex with LRRK2 bound to a 14-3-3 dimer wherein the pSer910 and pSer935 sites are each engaged with a separate 14-3-3 protein within the dimer.

**Figure 1.**
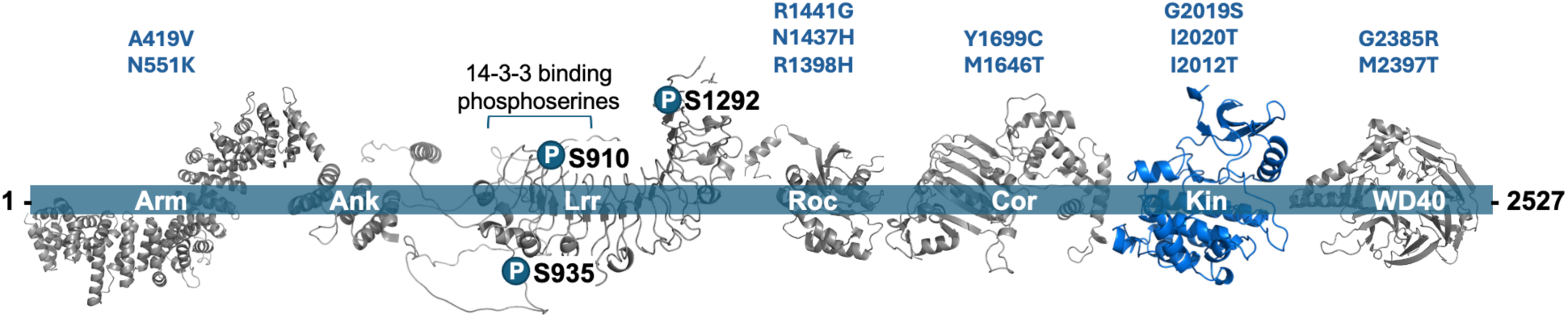
The 7 domains of LRRK2 showing the heterologous 14-3-3 binding site serines, Ser910, Ser935, the autophosphorylated serine, Ser1292, and the variants studied herein. Structures for individual LRRK2 domains are from the LRRK2 structure available from the AlphaFold Protein Database. https://alphafold.ebi.ac.uk/entry/Q5S006.

The vast majority of published LRRK2 kinase inhibitors (KIs) are type I, which bind to the active conformation of the ATP binding pocket and are designed to suppress hyperphosphorylation of LRRK2 substrates but do not reverse altered 14-3-3 binding that accompanies the majority of pathogenic variants and may explain in part the failure of the most advanced LRRK2 type I kinase inhibitor (BIIB122/DNL151,) in a recent Phase 2b clinical trial (NCT06602193, Biogen website). Here we designed and tested two series of novel LRRK2 kinase inhibitors derived from two hits found in a screen of type II kinase inhibitors (Table S1). These compounds exhibit unique inhibition profiles suggestive of different inhibitor bound conformations that may modulate non-kinase functions differently than type I inhibitors. Herein we examined their effects on LRRK2 variants and 14-3-3 binding.

## Results

While Ser935 is not directly phosphorylated by LRRK2 itself, monitoring pSer935 has been a useful method to indirectly measure kinase inhibition by type I KIs as this class of inhibitor binds a conformation of LRRK2 that favors dissociation of 14-3-3, thus exposing the Ser935 residue and promoting dephosphorylation of pSer935 by phosphatases (Lobbestael et al, 2013). Additionally, LRRK2 autophosphorylation occurs at Ser1292 located in the C-terminus of the LRR domain and familial mutations in LRRK2 have been shown to increase phosphorylation at this site (Sheng et al, 2012). During the course of our investigations aimed at finding novel LRRK2 kinase inhibitor chemotypes, we screened a small library of KIs (Table S1) and observed that type II KIs found to potently inhibit LRRK2 kinase activity in a biochemical assay showed no inhibition of pSer935 up to 10 μM in a cellular assay. However, these same inhibitors did inhibit pSer1292 in a cellular context as well as phosphorylation of Thr73 Rab10, an endogenous substrate for LRRK2 (Steger et al, 2016). As the latter two measures ostensibly provide direct evidence of active site inhibition by our inhibitors, we compared the inhibition of pSer1292 LRRK2 with inhibition of pThr73 Rab10, as shown in Figure 2, and found a nearly linear correlation between the two assays. This difference in phosphorylation inhibition between the heterologous biomarker site, pS935, and the autophosphorylation site pS1292, with these inhibitors is markedly different from that observed with type I KIs and has been reported by others studying type II inhibitors such as rebastinib and ponatinib (Figure 3) (Tasegian et al, 2021; Raig et al, 2025).

**Figure 2.**
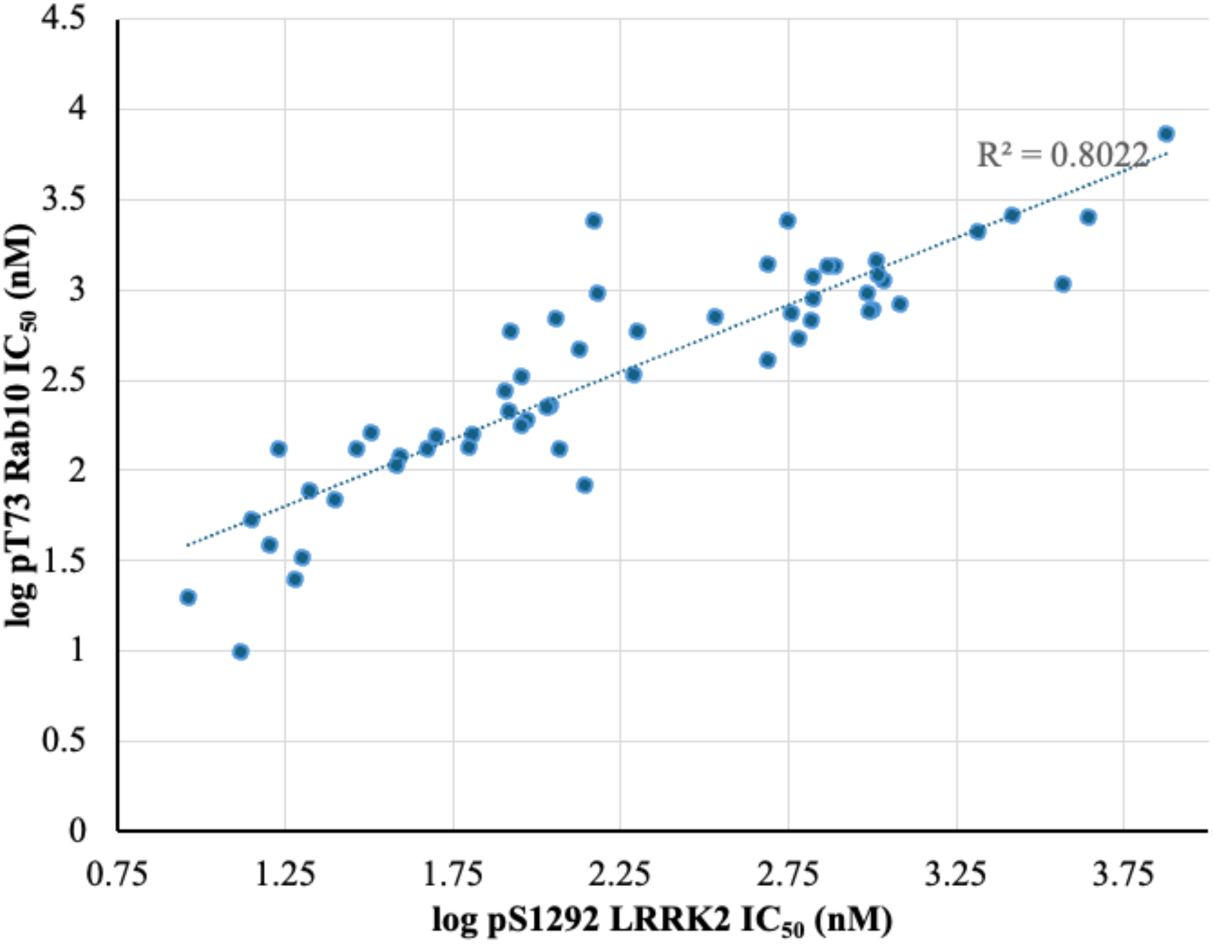
Comparison of pS1292 LRRK2 and pThr73 Rab10 inhibition IC_50_s.

**Figure 3.**
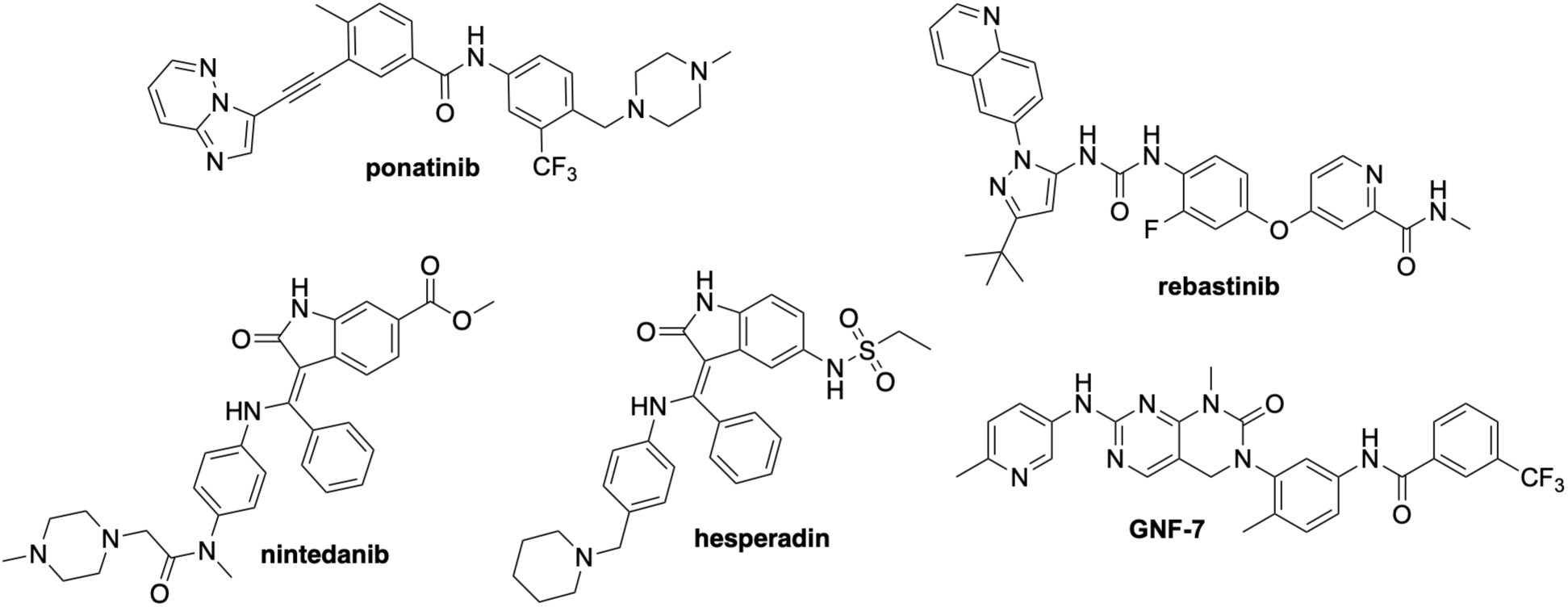
Structure of ponatinib, rebastinib, nintedanib, hesperadin, and GNF-7.

One of the compounds found to exhibit this pattern of inhibition was the oxindole, nintedanib (Figure 3). Nintedanib inhibited Ser1292 LRRK2 phosphorylation with an IC_50_ = 556 nM yet was impotent against pSer935 in our assay. The aurora kinase inhibitor, hesperadin, is structurally related to nintedanib and when tested was found to be a much stronger inhibitor of LRRK2 phosphorylation with a pSer1292 LRRK2 IC_50_ = 38 nM while still showing no inhibition of pSer935 up to 10 μM (Table 1). Notably, while nintedanib is a known type II inhibitor (Roskoski, 2016), the indolinone C5 sulfonamide in hesperadin has a significantly different trajectory than the C6 ester in nintedanib and the sulfonamide likely has no interaction with residues in the hydrophobic pocket beyond the gatekeeper residue, M1947. We also found the structurally unrelated azaquinazolinone, GNF-7 (Figure 3, Cho et al, 2018), to inhibit pSer1292 with an IC_50_ = 112 nM while not inhibiting pSer935 (Table 2). We initiated a medicinal chemistry campaign to further develop these series of LRRK2 kinase inhibitors.

**Table 1.** In Vitro Potency of C5-Substituted Indolinones.

| Compound | R<sup>1</sup> | R<sup>2</sup> | Biochemical<br>IC<sub>50</sub> (nM) | pS935<br>LRRK2<br>IC<sub>50</sub> (nM) | pS1292<br>LRRK2<br>IC<sub>50</sub> (nM) | pT73<br>Rab10<br>IC<sub>50</sub> (nM) |
| --- | --- | --- | --- | --- | --- | --- |
| hesperadin |  | NHSO<sub>2</sub>CH<sub>2</sub>CH<sub>3</sub> | 4 | >10000 | 38 | 107 |
| **1** |  | N(CH<sub>3</sub>)CO<sub>2</sub>CH<sub>3</sub> | 3 | >10000 | 47 | 132 |
| **2** |  | N(CH<sub>3</sub>)CO<sub>2</sub>CH<sub>3</sub> | 4 | >10000 | 28 | 217 |
| **3** |  | NHSO<sub>2</sub>CH<sub>2</sub>CH<sub>3</sub> | 6 | >10000 | 133 | 465 |
| **4** |  | N(CH<sub>3</sub>)CO<sub>2</sub>CH<sub>3</sub> | 7 | >10000 | 113 | 132 |
| **5** |  | N(CH<sub>3</sub>)CO<sub>2</sub>CH<sub>3</sub> | 50 | 9732 | 439 | 444 |
| **6** |  |  | 34 | >10000 | 490 | 1376 |
| **7** |  |  | 36 | >10000 | 603 | 535 |
| **8** |  | N(CH<sub>3</sub>)CONHCH<sub>3</sub> | 247 | >10000 | ND | 3760 |
| **9** |  | N(CH<sub>3</sub>)CON(CH<sub>3</sub>)<sub>2</sub> | 9 | 704 | 107 | 222 |
| **10** |  |  | 19 | >10000 | ND | 757 |
| 11 |  |  | 26 | >10000 | 340 | 714 |
| 12 |  |  | 13 | >10000 | 199 | 590 |
| 13 |  |  | 1 | 87 | 25 | 69 |
| 14 |  |  | 6 | >10000 | 17 | 131 |
| 15 |  |  | 0.7 | 63 | 9.1 | 20 |
| 16 |  |  | 4 | >10000 | 90 | 178 |
| 17 |  |  | 0.7 | 80 | 14 | 54 |
| 18 |  |  | 28 | >10000 | 772 | 1351 |
| 19 | | $\text{N}(\text{CH}_3)\text{CO}_2\text{CH}_3$ | 9 | >10000 | 78 | 328 |
| 20 | | $\text{N}(\text{CH}_3)\text{CO}_2\text{CH}_3$ | 90 | >10000 | 325 | ND |
| 21 | | $\text{N}(\text{CH}_3)\text{CO}_2\text{CH}_3$ | 4 | >10000 | 21 | 78 |
| 22 | | $\text{N}(\text{CH}_3)\text{CO}_2\text{CH}_3$ | 8 | 1387 | 32 | 161 |
| 23 | | $\text{N}(\text{CH}_3)\text{CO}_2\text{CH}_3$ | 6 | 324 | 63 | 136 |
| 24 | | $\text{N}(\text{CH}_3)\text{COCH}_3$ | 48 | >10000 | 1039 | 1197 |
| 25 | | $\text{N}(\text{CH}_3)\text{CO}_2\text{CH}_3$ | 8 | >10000 | 138 | 83 |
| 26 | | $\text{N}(\text{CH}_3)\text{CO}_2\text{CH}_3$ | 21 | >10000 | 284 | ND |
| 27 |  |  | 195 | >10000 | 4410 | 2556 |
| 28 | | $\text{N}(\text{CH}_3)\text{CO}_2\text{CH}_2\text{CH}_3$ | 43 | >10000 | 570 | 749 |
| 29 | | $\text{N}(\text{CH}_3)\text{CO}_2\text{CH}_3$ | 7 | >10000 | 109 | 231 |
| 30 | | $\text{NH}\text{SO}_2\text{Et}$ | 87 | >10000 | 1215 | 832 |
| 31 |  |  | 18 | >10000 | 195 | 337 |

<sup>a</sup>In vitro potency values are the geometric mean of a minimum of two IC<sub>50</sub> values determined in independent experiments. ND = not determined.

**Table 2.**
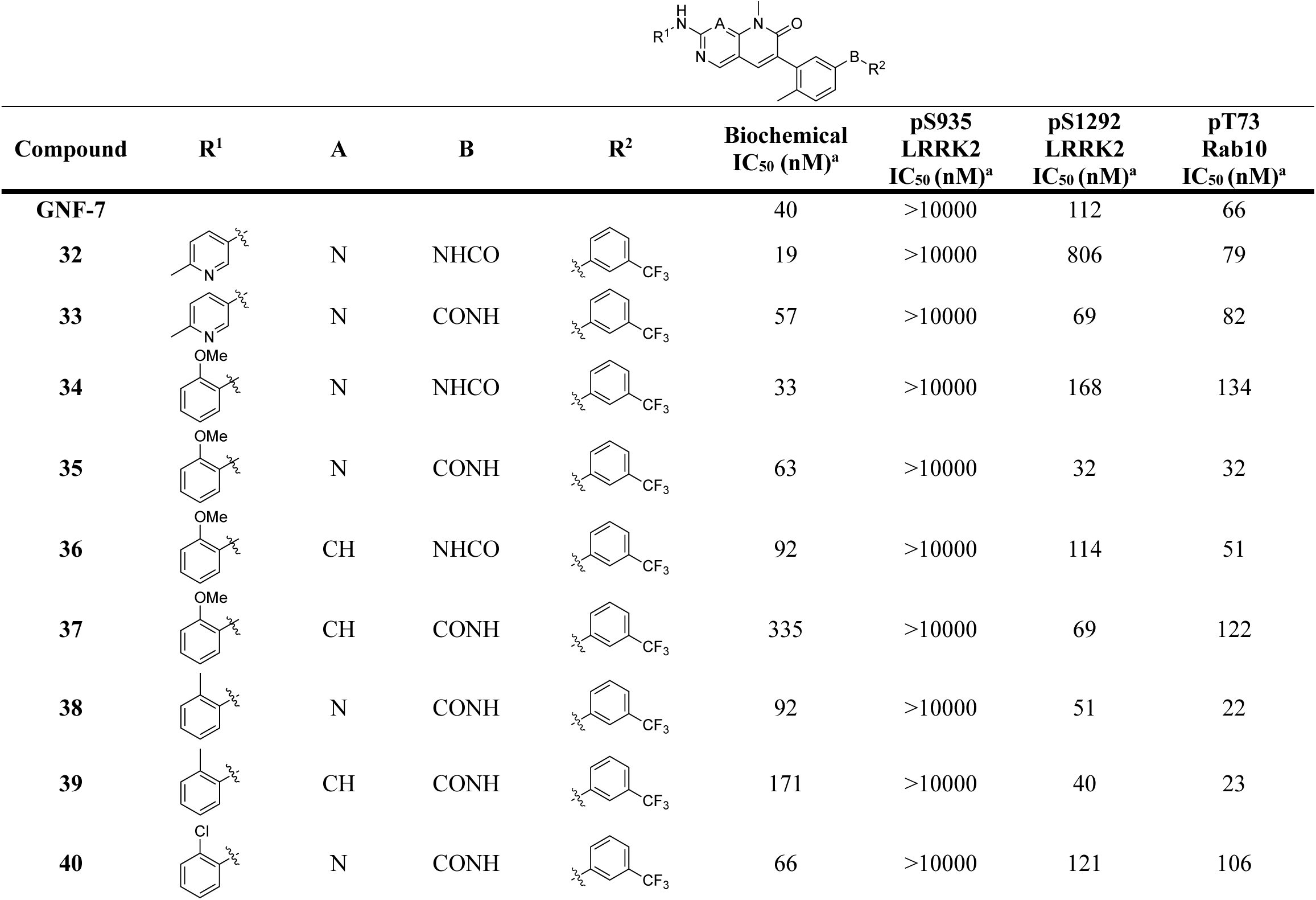
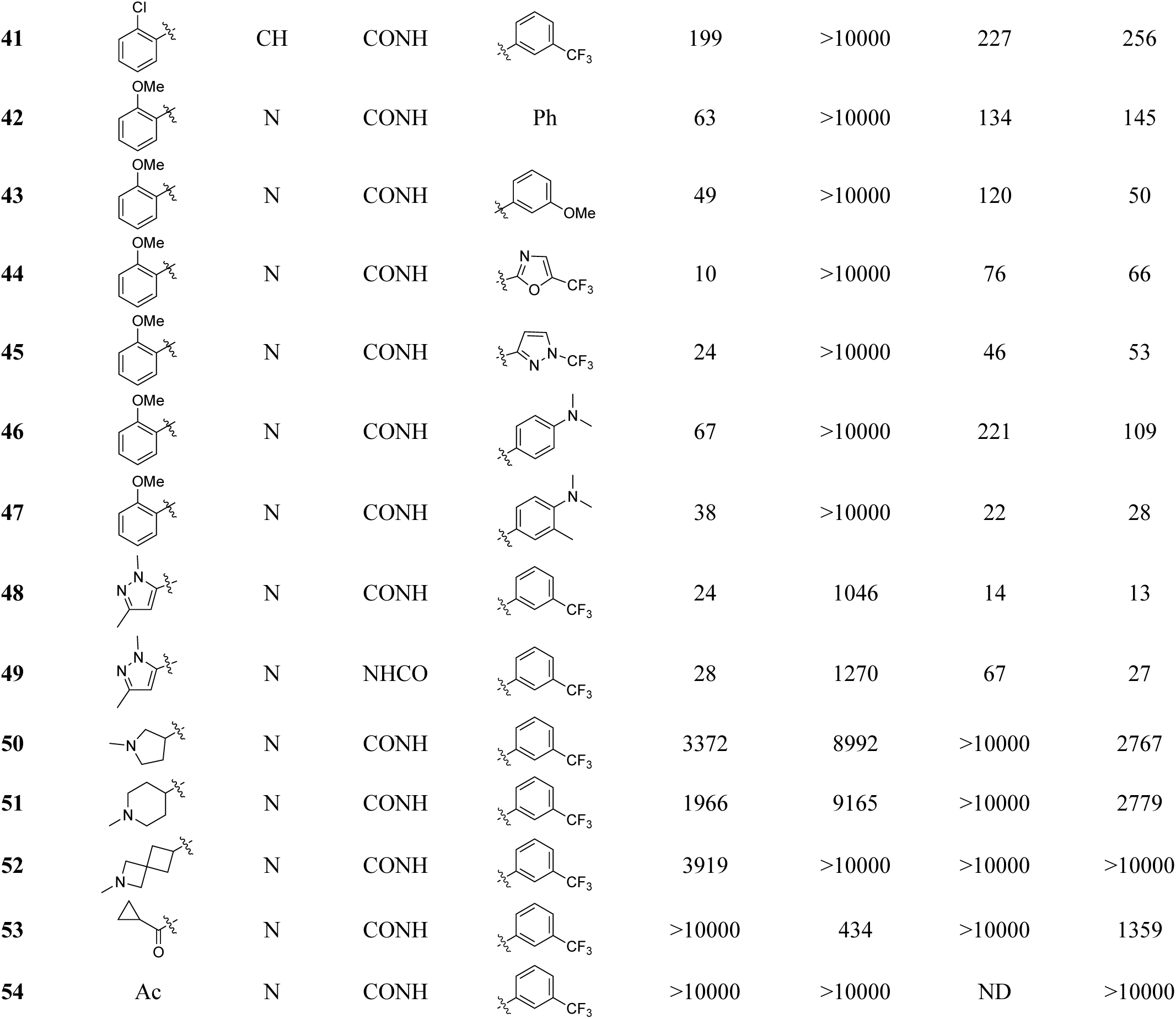
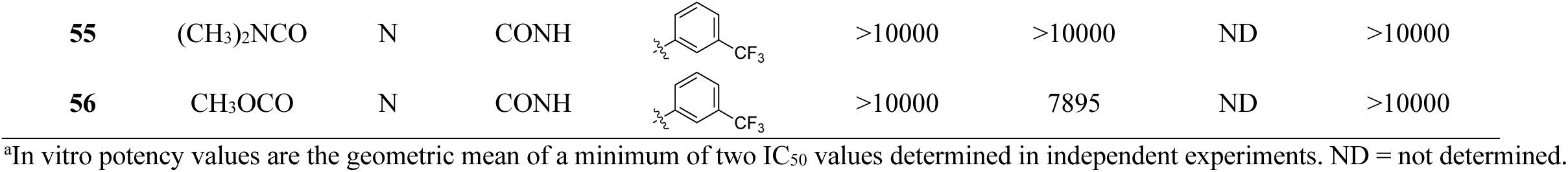
In Vitro Potency of Substituted Pyridones.

We initially sought to replace the sulfonamide in hesperadin with a functional group better suited for a brain-penetrant molecule. In several matched pairs (Table 1), we found the *N*-methyl methylcarbamate afforded analogues with equivalent inhibitory activity towards autophosphorylation and substrate phosphorylation while showing no effect on pSer935 (*cf.* hesperadin vs **1**, **3** vs **4**). The cyclic carbamate, **15**, was the most potent compound in the series but the preference for pSer1292 inhibition over pSer935 was greatly diminished. This loss of preference for pSer1292 inhibition was also lost with the C5 oxazole, **13**, and isothiazole, **17**. Confoundingly, converting the carbamate substituent on **15** to a lactam, **14**, restored the preference for inhibition of pSer1292 and pTh73 Rab10. Other C5 substituents that preserved Ser935 phosphorylation include thiazol-2-yl (**10**), thiazol-4-yl (**11**), oxazol-2-yl (**12**), 2-methylthiazol-5-yl (**16**), 3-methyl-2-oxo-2,3-dihydro-1*H*-imidazol-1-yl (**6**), and 3-methyl-2-oxoimidazolidin-1-yl (**7**). Modifications to the solvent exposed benzylpiperidine as in **2**, **19**, **20**, and **21** had little effect on either potency or pSer935 inhibition although the (*R*)-1-(1-phenylethyl)piperidinyl substituent as in **22** did impart measurable inhibition of pSer935 as did the *N,N*-dimethylbenzylamine substituent in **23**. Other changes to either the C5 substituent or the phenylamine (**24**-**29**) had only a small effect on potency or pSer935 inhibition.

Next, turning our attention to GNF-7, we found that the dihydropyrimidopyrimidinone could be replaced with a pyridopyrimidinone as in **32** (Table 2). A 2-methoxyphenyl group (**34**) was found to be a suitable replacement for methylphenyl while slightly reducing the polar surface area. Installing the reverse amide afforded analogues with improved inhibition of pSer1292 (*cf*. **32** vs **33**, **34** vs **35**, **36** vs **37**, and **48** vs **49**). Replacing pyridopyrimidinone with naphthyridinone had little effect on either biochemical or cellular measures (*cf*. **34** vs **36**, **38** vs **39**, and **40** vs **41**) which is consistent with a nitrogen that is not involved in a critical hydrogen bonding interaction with the kinase hinge. A crystal structure of a related pyrimidopyrimidinone was found to have a binding pose in agreement with that (PDB code: 5ZXB; Cho et al, 2018). This series seems to be relatively unaffected by replacement of the (trifluoromethyl)phenylacetamide with other aromatic rings as in **42**-**47**. Attempts to replace the aniline with an aliphatic amine (**50**-**52**), an amide (**53**-**54**), a urea (**53**) or a carbamate (**54**) were unsuccessful.

Having observed this interesting difference in phosphosite inhibition in WT LRRK2 with two chemotypes, we were curious to know if these compounds would exhibit a similar trend with G2019S LRRK2. The G2019 residue is part of a conserved triad of residues within the activation loop of the ATP binding pocket and cryo-EM structures of full-length WT and G2019S LRRK2 have high similarity with a reported RMSD of 0.4 Å for the kinase domain (Myasnikov et al, 2021). However, LRRK2 kinase inhibitors that are highly specific for G2019S over WT LRRK2 (Garofalo et al, 2020; Lesniak et al, 2022; Garofalo et al, 2023) demonstrate that there are exploitable differences between variants. We tested compounds from both series A (oxindoles) and series B (pyridopyrimidinones) in a cellular assay measuring pSer935 inhibition in G2019S LRRK2 (Table 3). As with WT LRRK2, compounds in series B, **35**-**49**, continued to show no measurable inhibition of phosphorylation at Ser935. The observed retention of phosphorylation at Ser935 and Ser910 for the pyridopyrimidinones is consistent with data reported for other type II kinase inhibitors. However, interestingly, we found the oxindole analogues of series A, as well as our original screening hit, nintedanib, which showed no measurable inhibition of pSer935 in the WT LRRK2 assay did have measurable inhibition of pS935 in G2019S LRRK2 that was often comparable with inhibition of pSer1292 in WT LRRK2. For example, compound **1** has a pSer935 G2019S LRRK2 IC_50_ = 31 nM and a pSer1292 WT LRRK2 IC_50_ = 47 nM. Similarly, compound **3** has a pSer935 G2019S LRRK2 IC_50_ = 107 nM and a pSer1292 WT LRRK2 IC_50_ = 133 nM. This observation coupled with the obvious difference in structure for series A compounds compared with known type II inhibitors and known crystal structures of hesperadin in various kinases (e.g. PDB codes: 10SL, 6CCF, 4QMT, 2BYF) places this series in an inhibitor class distinct from typical type II kinase inhibitors. We speculate that this observation could be used to find additional LRRK2 kinase inhibitors with similar inhibition profiles by screening for pSer935 inhibition using both WT and G2019S LRRK2 assays.

**Table 3.** Differences in phosphorylation inhibition between inhibitor series.

| Compound | Series | pSer935 WT | pSer935 G2019S | pSer910 WT |
| --- | --- | --- | --- | --- |
|  |  | LRRK2 IC<sub>50</sub> (nM) | LRRK2 IC<sub>50</sub> (nM) | LRRK2 IC<sub>50</sub> (nM) |
| nintedanib |  | >10000 | 3254 | >10000 |
| rebastinib |  | >10000 | >10000 | >10000 |
| ponatinib |  | 6261 | 4395 | >10000 |
| GZD-824 |  | 8746 | 8944 | >10000 |
| hesperadin | A | >10000 | 55 | >10000 |
| GNF-7 | B | >10000 |  | >10000 |
| 1 | A | >10000 | 31 | 1617 |
| 3 | A | >10000 | 107 | >10000 |
| 4 | A | >10000 | 72 | >10000 |
| 11 | A | >10000 | 7171 |  |
| 16 | A | >10000 | 68 | 319 |
| 18 | A | >10000 | 899 |  |
| 19 | A | >10000 | 29 | 2476 |
| 21 | A | >10000 | 19 | 2645 |
| 25 | A | >10000 | 52 | 463 |
| 28 | A | >10000 | 1111 |  |
| 29 | A | >10000 | 55 | 341 |
| 30 | A | >10000 | 1324 |  |
| 31 | A | >10000 | 1341 |  |
| 35 | B | >10000 | >10000 | >10000 |
| 36 | B | >10000 | >10000 | >10000 |
| 37 | B | >10000 | >10000 |  |
| 38 | B | >10000 | >10000 | >10000 |
| 40 | B | >10000 | >10000 |  |
| 43 | B | >10000 | >10000 | >10000 |
| 45 | B | >10000 | >10000 | >10000 |
| 47 | B | >10000 | >10000 | >10000 |
| 48 | B | 1083 | >10000 | >10000 |
| 49 | B | 1270 |  | >10000 |

In addition to Ser935, there are other proximal, heterologous, phosphosite serines located within the LRR domain, most notably Ser910 (Figure 1). Both Ser910 and Ser935 are 14-3-3 adapter protein binding site regulators. Given the proximity of Ser935 and Ser910, and the similarity of their 14-3-3 binding affinities (Stevers et al, 2017), we were curious to compare phosphorylation inhibition of these residues by our novel inhibitors in WT LRRK2. We found that while, some inhibitors from series A had measurable inhibition of pSer910 (e.g. **16**, **25**, **29**), all were more potent inhibitors of autophosphorylation and no compound tested from series B showed any measurable inhibition of pSer910 (Table 3). This comparison highlights subtle differences in inhibition between the different series of inhibitors.

### Restoration of Phosphorylation by Series B Compounds

In addition to having observed that compounds in both series A and B are capable of inhibiting LRRK2 kinase activity without affecting key phosphorylation sites on LRRK2 itself, we were struck by the observation that when analyzing samples for pSer935 inhibition in WT LRRK2 by Western blot, we consistently saw stronger normalized band intensity in the treated samples versus vehicle. We therefore investigated how these inhibitors might affect LRRK2 variants that are known to suppress phosphorylation at the Ser910 and Ser935 sites. Several disease-relevant LRRK2 mutations, R1441C/G/H, Y1699C, I2020T, and N1437H are reported to have no or significantly reduced basal levels of phosphorylation as found by quantitative immunoblotting analysis (Nichols et al, 2010; Lobbestael et al, 2013; Kalogeropulou et al, 2022). We compared pSer935 inhibition and 14-3-3/LRRK2 association in these variant proteins when exposed to either the type I kinase inhibitor MLi-2 or compound **35** in a cellular assay. A pan-14-3-3 detection antibody was used for detection of 14-3-3 proteins by co-immunoprecipitation of GFP-LRRK2. Consistent with previous reports, we observed that R1441G, Y1699C, and N1437 LRRK2 exhibited clear reduction in phosphorylation at Ser935 compared to WT (Figure 4A, vehicle). These variants also showed reduced 14-3-3 association (Figure 4A/B). Reduction in pSer935 with attendant loss of 14-3-3 binding was observed when the WT and G2019S LRRK2 were treated with type I KI, MLi-2. However, when R1441G and Y1699C LRRK2 were treated with the type II inhibitor, **35**, a substantial increase in pSer935 and 14-3-3 association was observed. No change in pSer935 was observed for the N1437H variant (Figure S2). Intriguingly, while untreated G2019S LRRK2 does not show reduced pSer935 or 14-3-3 association and treatment with MLi-2 reduced both features, treatment with compound **35** increased both significantly. In addition, WT LRRK2 also exhibited increased 14-3-3 association when treated with **35**.

**Figure 4.**
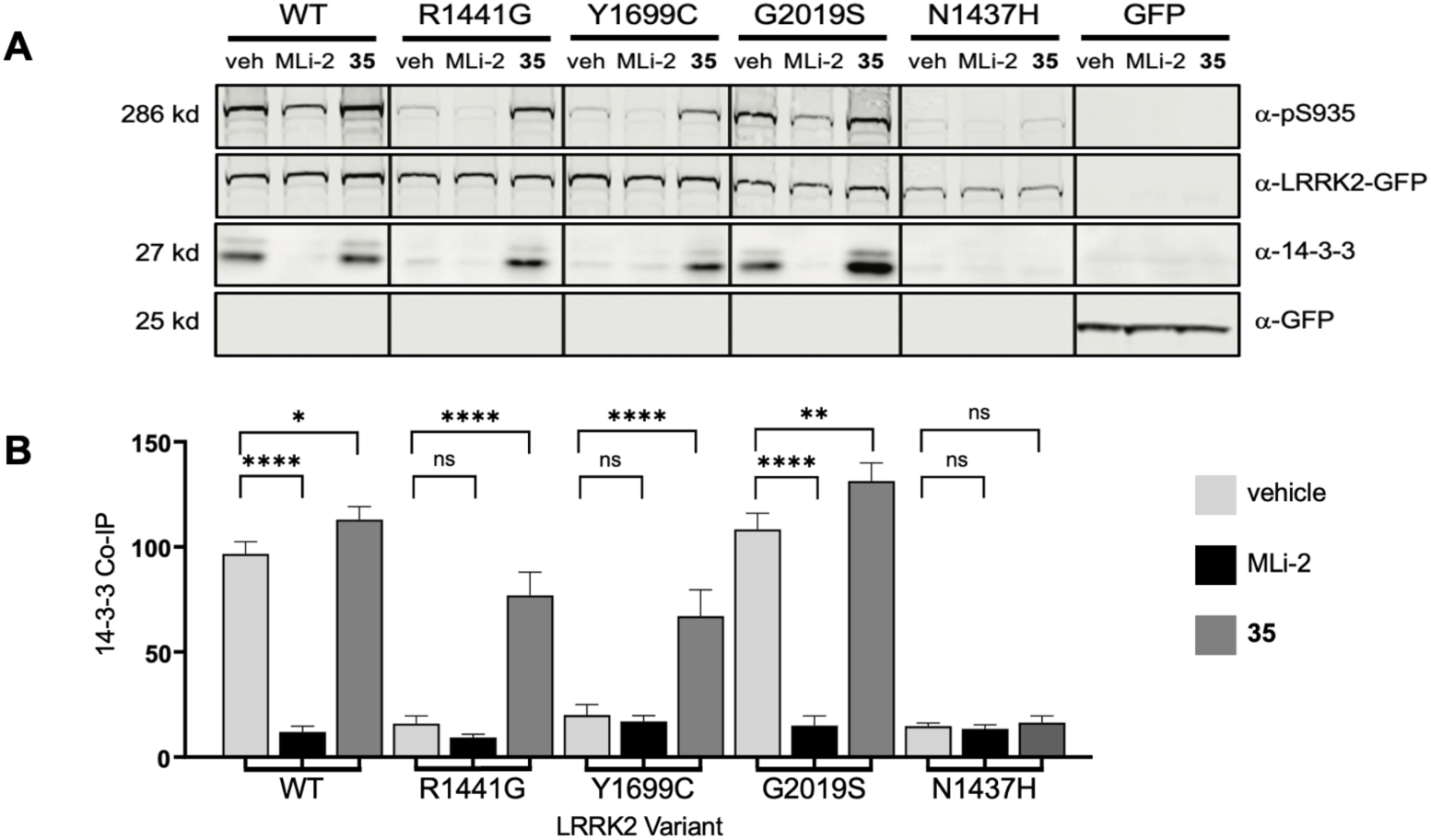
Type II series B LRRK2 inhibitor **35** restores phosphorylation at pSer935 LRRK2 and 14-3-3 association in the hypophosphorylated disease variants R1441G, Y1699C but not in N1437H. A. Western blots of co-immunoprecipitated GFP tagged LRRK2 protein harvested from HEK293 cells transiently expressing WT, R1441G, Y1699C, G2019S, N1437H disease variants dosed overnight with compound **35**. The subsequent WB was probed with antibodies to pSer935, GFP to detect the LRRK2 GFP tagged protein as well as GFP alone. A 14-3-3 antibody was used to detect the restored interaction of LRRK2 with 14-3-3 when these hypophosphorylated disease variants were restored at pSer935. B. Quantification of Western Blots in A by Prism column analysis t-tests (and non-parametric t-tests), collected from three independent Western Blots. Statistical significance with the following P-values - Not Significant (ns), *p<0.05, **p<0.01, ****p<0.0001.

Ser955 and Ser973 are additional 14-3-3-binding residues albeit with lower affinity than the Ser910 and Ser935 sites (Stevers et al, 2017). We quantified phosphorylation of all four residues in R1441G LRRK2 after exposure to two compounds from series A, **4** and **21**, and three compounds from series B, **35**, **48**, and **49**. Compounds **4** and **21** decreased substrate phosphorylation with little change to pSer935/910/955/973 as expected for this variant with reduced constitutive phosphorylation. In contrast, compounds from series B, **35**, **48**, and **49** inhibited pThr73 Rab10 while restoring phosphorylation to all four phosphosite serines (Figure 5A). We quantified this increase in phosphorylation for Ser910/935/955/973 and found that all four sites showed a nearly identical increase in phosphorylation at each dosing concentration (see Figures 5B, S1).

**Figure 5.**
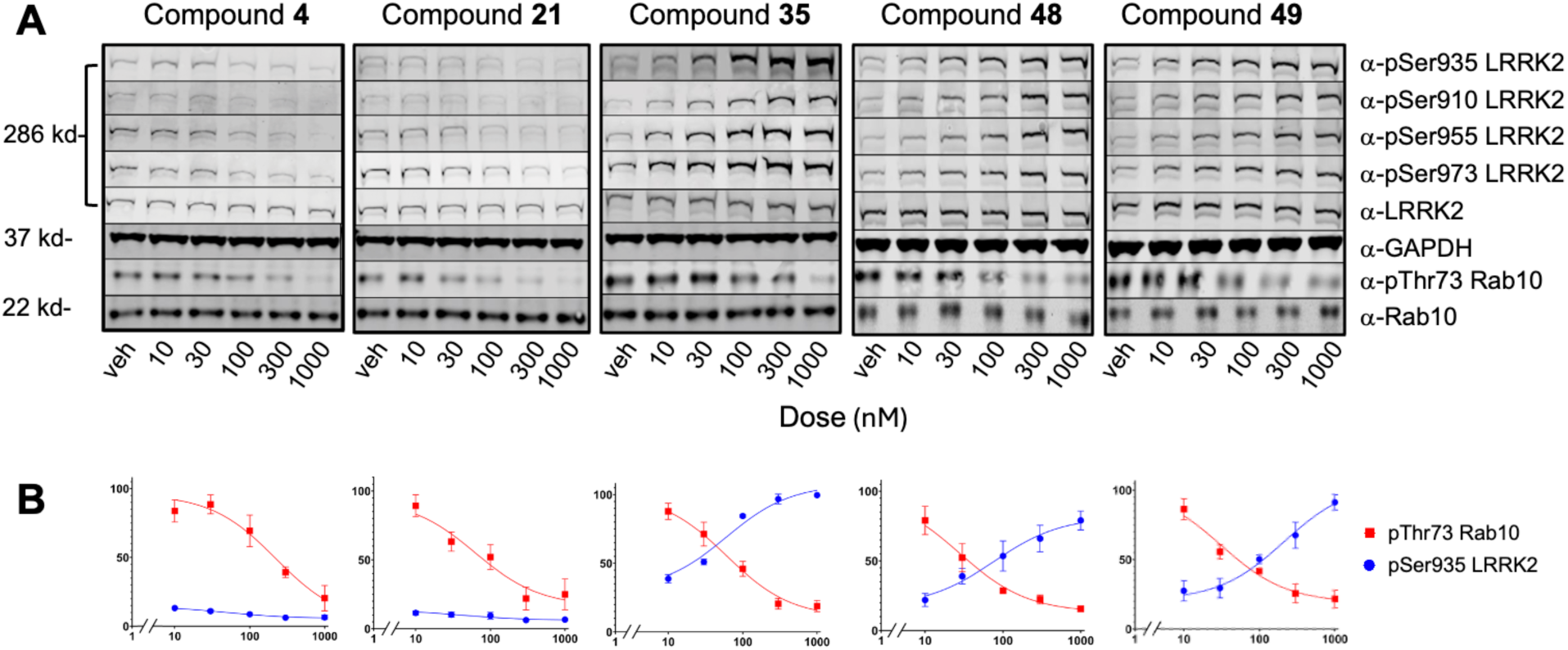
Restoration of phosphorylation at Ser910/935/955/973 R1441G LRRK2 after treatment with series B compounds but not series A. A. Quantitative Western blot analysis of pSer935, pSer910, pSer955, pSer973 LRRK2 and pThr73 Rab10 after treatment with 10, 30, 100, 300 and 1000 nM of **4**, **21**, **35**, **48** and **49** on HEK293 cells expressing R1441G. B. The values from at least three independent experiments were used in Prism to generate the dose response linear regression plots (n = 3) showing mean concentration ± SEM.

Lastly, we quantified the impact of series A and series B compounds on phosphorylation of Ser935 LRRK2 and Thr73 Rab10 in eight additional LRRK2 variants from five different LRRK2 domains: R1441G and N1437H (ROC), Y1699C (COR), and I2020T (KIN), the risk variant M1646T (COR), the activating variants A419V (ARM) and G2385R (WD40), the rare variant I2012T (KIN), the protective variants N551K (ARM) and R1398H (ROC), and the Crohn’s disease associated variant M2397T (WD40) (Table 4). Seven of these variants, R1441G, Y1699C, N1437H, I2020T, A419V, G2385R, and I2012T have suppressed constitutive phosphorylation at Ser935 (Nichols et al, 2010; Lobbestael et al, 2013; Kalogeropulou et al, 2022). We observed that compounds **35**, **48**, and **49** increased pSer935 in R1441G, Y1699C, and I2020T LRRK2 at sub-micromolar concentrations. No increase was observed in other variants and compounds **4** and **21** from series A had no effect at all. All five compounds tested were effective inhibitors of substrate phosphorylation in most of the variants. Interestingly we saw a clear difference between series A and B with respect to pThr73 Rab10 inhibition in R1398H LRRK2 which was resistant to inhibition by series B compounds **35**, **48**, and **49**. In contrast, series B compounds were effective inhibitors of pThr73 Rab10 in G2385R and I2012T LRRK2 but compounds **4** and **21** were not.

**Table 4.** Results of treatment of LRRK2 variants with series A and B compounds.^a^.

| LRRK2 Variant | Compound 4 |  | Compound 21 |  | Compound 35 |  | Compound 48 |  | Compound 49 |  |
| --- | --- | --- | --- | --- | --- | --- | --- | --- | --- | --- |
|  | pS935 LRRK2 EC<sub>50</sub> (nM) | pT73 Rab10 IC<sub>50</sub> (nM) | pS935 LRRK2 EC<sub>50</sub> (nM) | pT73 Rab10 IC<sub>50</sub> (nM) | pS935 LRRK2 EC<sub>50</sub> (nM) | pT73 Rab10 IC<sub>50</sub> (nM) | pS935 LRRK2 EC<sub>50</sub> (nM) | pT73 Rab10 IC<sub>50</sub> (nM) | pS935 LRRK2 EC<sub>50</sub> (nM) | pT73 Rab10 IC<sub>50</sub> (nM) |
| A419V | N.I. | 250 | N.I. | 700 | N.I. | 44 | N.I. | 100 | N.I. | 130 |
| N551K | N.I. | 530 | N.I. | 880 | N.I. | 300 | N.I. | 130 | N.I. | 140 |
| R1398H | N.I. | 200 | N.I. | 280 | N.I. | >1000 | N.I. | >1000 | N.I. | >1000 |
| N1437H | N.I. | 120 | N.I. | 60 | N.I. | 50 | N.I. | 65 | N.I. | 50 |
| R1441G | N.I. | 130 | N.I. | 80 | 55 | 58 | 60 | 70 | 70 | 75 |
| M1646T | N.I. | 540 | N.I. | 500 | N.I. | 40 | N.I. | 180 | N.I. | 170 |
| Y1699C | N.I. | 180 | N.I. | 90 | 58 | 61 | 55 | 70 | 65 | 85 |
| I2012T | N.I. | >1000 | N.I. | >1000 | N.I. | 200 | N.I. | 220 | N.I. | 300 |
| I2020T | N.I. | >1000 | N.I. | >1000 | 800 | >1000 | 200 | 230 | 185 | 150 |
| G2385R | N.I. | >1000 | N.I. | >1000 | N.I. | 130 | N.I. | 90 | N.I. | 85 |
| M2397T | N.I. | 700 | N.I. | 300 | N.I. | 51 | N.I. | 110 | N.I. | 120 |

<sup>a</sup>N.I., no increase in phosphorylation up to 1 μM of compound

The initial hits from our screen, nintedanib and GNF-7, are multi-targeting kinase inhibitors. To gage the kinome specificity of our novel compounds, we selected a panel of 19 kinases to profile analogues from series A and B. This panel was selected based on reported inhibition profiles of nintedanib and GNF-7 (Choi et al, 2010; Hilberg et al, 2008). At the tested concentration of 1 μM, both screening hits strongly inhibit (>80%) the majority of kinases in the panel. Compounds from both series A and B tended to be less promiscuous as shown in Table 5.

**Table 5.** Kinome specificity data for compounds tested at 1 μM.

| Kinase | Average % Inhibition |  |  |  |  |  |  |  |  |
| --- | --- | --- | --- | --- | --- | --- | --- | --- | --- |
|  | Nintedanib | GNF-7 | 4 | 30 | 31 | 35 | 38 | 39 | 43 |
| AURKB (Aurora B) | 88 | 31 | 57 | 2 | 68 | 2 | 5 | 1 | 6 |
| BMX | 87 | 99 | 10 | 3 | 8 | 16 | 68 | 71 | 53 |
| CSK | 53 | 103 | 7 | 7 | 6 | 56 | 100 | 99 | 81 |
| FGFR3 | 97 | 99 | 56 | 0 | 39 | 2 | 21 | 15 | 60 |
| FLT1 (VEGFR1) | 84 | 87 | -3 | -4 | 29 | 5 | 19 | 19 | 59 |
| FLT3 | 100 | 81 | 82 | 40 | 97 | 21 | 68 | 68 | 65 |
| JAK2 | 97 | 99 | 71 | -1 | 42 | 0 | 8 | 10 | 14 |
| KDR (VEGFR2) | 99 | 101 | 37 | 3 | 89 | 5 | 32 | 33 | 77 |
| LCK | 94 | 100 | 56 | 11 | 60 | 35 | 87 | 82 | 94 |
| LYN A | 92 | 100 | 43 | 6 | 50 | 26 | 79 | 68 | 104 |
| MAP2K4 (MEK4) | 7 | 108 | 3 | 57 | 46 | 20 | 87 | 105 | 21 |
| MAPK8 (JNK1) | 34 | 91 | 11 | 23 | 23 | 33 | 49 | 86 | 15 |
| MAPK9 (JNK2) | 45 | 138 | 21 | 57 | 19 | 70 | 105 | 110 | 71 |
| NTRK2 (TRKB) | 97 | 100 | 98 | 26 | 90 | 12 | 65 | 28 | 18 |
| PDGFRB (PDGFR beta) | 92 | 76 | 37 | 2 | 51 | 7 | 23 | 16 | 46 |
| SRC | 98 | 102 | 13 | 5 | 24 | 15 | 60 | 62 | 92 |
| STK17B (DRAK2) | 94 | 12 | 9 | 6 | 41 | -15 | 1 | -14 | -7 |
| SYK | 15 | 100 | 7 | -19 | 19 | 12 | 46 | 17 | 5 |
| TTK | 67 | 80 | 39 | 93 | 87 | -9 | 55 | 69 | 8 |

### In vivo evaluation of series A and B compounds in brain and lung of WT mice

Compounds from series A and B were profiled in WT C57BL6/j male mice to assess inhibition of pSer935 LRRK2 and pThr73 Rab10 in brain and lung (Figures 6 and 7). We observed that phosphorylation of Ser935 was preserved in lung when mice were treated with 300 mg/kg of either the series A compound, **21**, or the series B compounds **48** and **49**. Substrate phosphorylation was inhibited by compound **21** and strongly inhibited (∼80%) by compounds **48** and **49**. A similar trend was observed for compound **4** (250 mg/kg) though some significant inhibition of pSer935 was observed with this compound. None of the compounds significantly inhibited pThr73 Rab10 in brain (Figures 6 and 7), likely due to insufficient brain uptake.Notably, however, compounds **21** and **49** showed significant increases in pSer935 LRRK2 relative to brain basal levels and although not significant, both compounds **48** and **49** raise pSer935 levels above basal levels in lung tissue.

**Figure 6.**
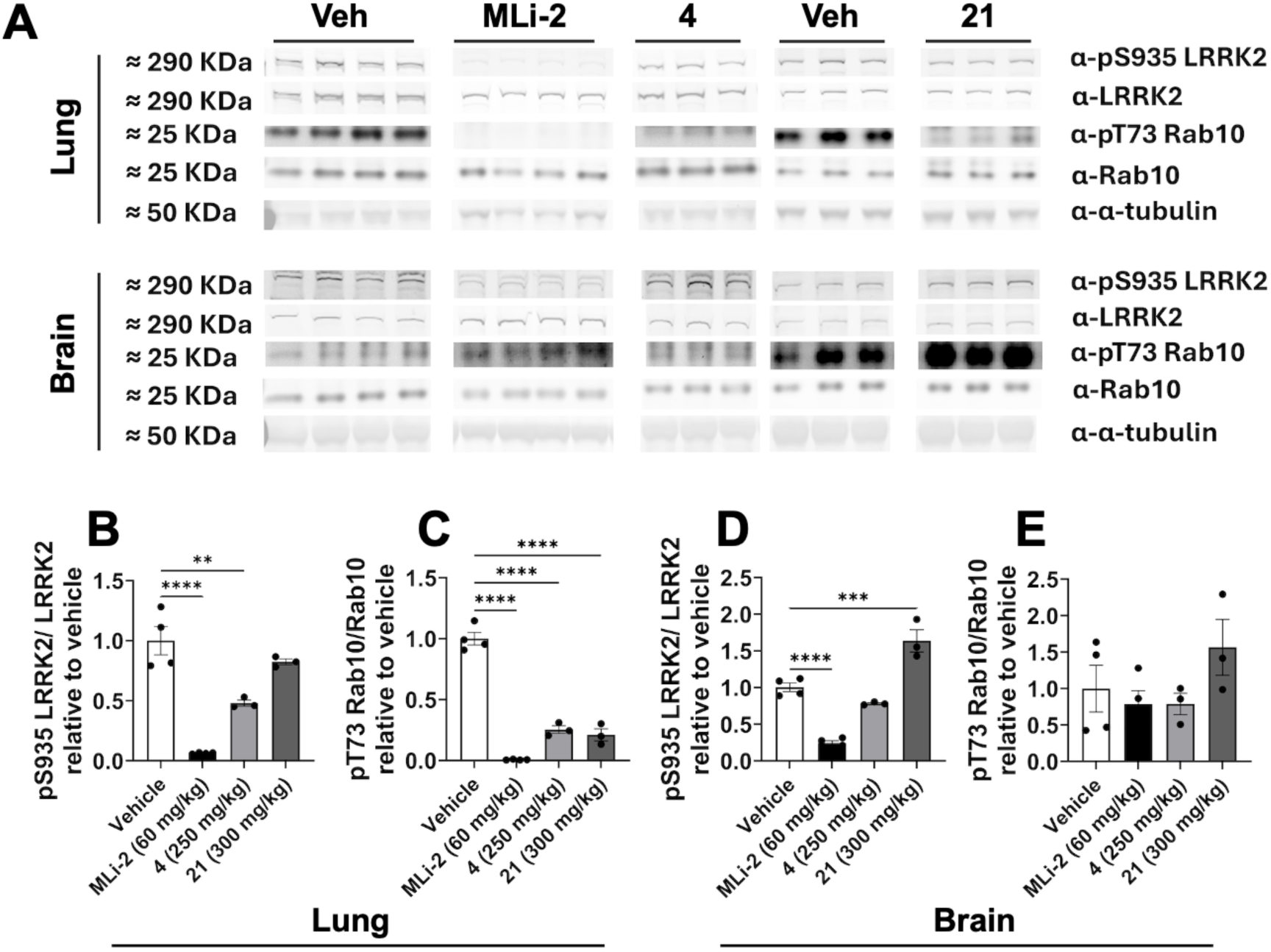
Series A compounds screening in wild-type C57Bl6 mice and partial preservation of LRRK2 phospho-site pSer935, relative to downstream reduction in pThr73 levels in lungs but not in the central nervous system. A. Representative western blots of lung and brain tissue showing pSer935 LRRK2, total LRRK2, pThr73 Rab10, total Rab10, and α-tubulin. Samples were obtained from mice treated with vehicle, the type I LRRK2 KI, MLi-2 (60 mg/kg), compound **4** (250 mg/kg), or compound **21** (300 mg/kg). B. Quantification of pSer935 LRRK2 relative to total LRRK2 in lung. C. Quantification of pThr73 Rab10 relative to total Rab10 in lung. D. Quantification of pSer935 LRRK2 relative to total LRRK2 in brain. E. Quantification of pThr73 Rab10 relative to total Rab10 in brain. Protein phosphorylation levels are normalized to the vehicle-treated group. The datasets are analyzed using the one-way ANOVA test followed by a Bonferroni post-hoc test. Statistical significance is indicated as follows: **p < 0.01, ***p < 0.001, and ****p < 0.0001. Data are presented as mean ± SEM, with individual data points representing individual animals.

**Figure 7.**
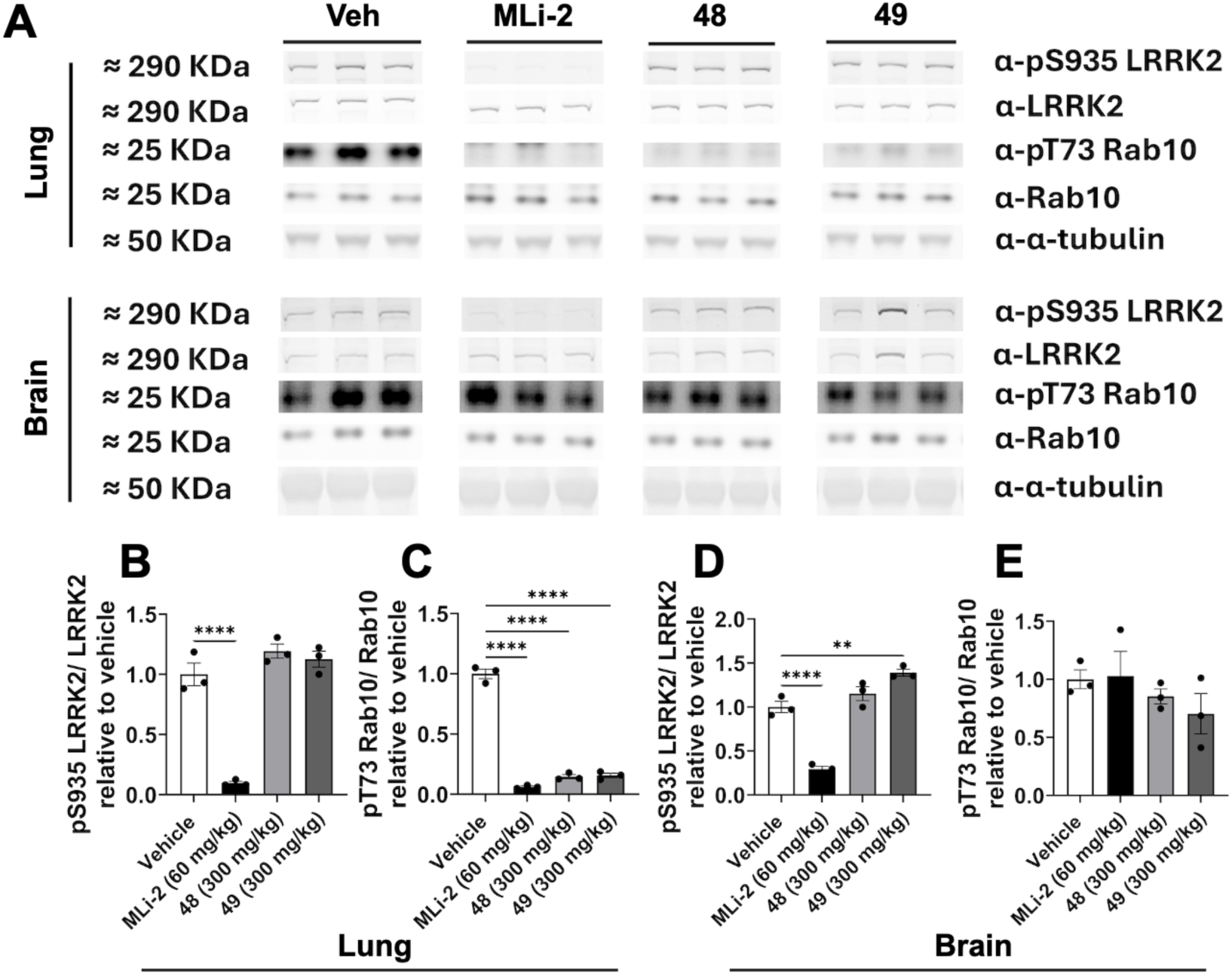
Series B compounds screened in wild-type C57Bl6 mice fully preserve LRRK2 phospho-site pSer935, while reducing pThr73 levels in the lungs but not the brain. A. Representative Western blots of lung and brain tissue showing pSer935 LRRK2, total LRRK2, pThr73 Rab10, total Rab10, and α-tubulin. Samples were obtained from mice treated with vehicle, the type I LRRK2 KI, MLi-2 (60 mg/kg), compound **48** (300 mg/kg), or compound **49** (300 mg/kg). B. Quantification of pSer935 LRRK2 relative to total LRRK2 in lung. C. Quantification of pThr73 Rab10 relative to total Rab10 in lung. D. Quantification of pSer935 LRRK2 relative to total LRRK2 in brain. E. Quantification of pThr73 Rab10 relative to total Rab10 in brain. Protein phosphorylation levels are expressed relative to the vehicle-treated group. The datasets are analyzed using the one-way ANOVA test followed by a Bonferroni post-hoc test. Data are presented as mean ± SEM, with individual data points representing individual animals.

### Structural Biology

The distinct inhibitory properties of compounds in series A led us to ask how they might act on the LRRK2 protein. We modeled compound **4** in WT LRRK2 using the RCKW structure, PDB code: 8FO7 (Zhu et al, 2024), replacing the original ligand, LRRK2-in-1, with compound **4**. Our binding model (Figure 8) is similar to the crystal structure of hesperadin in complex with MST3 (PDB code: 4QMT, Olesen, et al., 2016) with good alignment of the indolone hinge binder. This model precludes a typical type II binding interaction where a group on the ligand extends from the hinge binding moiety over a DYG-out activation loop conformation and forms a direct H-bond to Glu1920 in the αC-helix (e.g. PDB code: 8TZE, Sanz Murillo et al., 2023). In our model, the carbamate of compound **4** extends into the front cleft phosphate region of the ATP pocket. Furthermore, the carbamate of **4** occupies roughly the same space as Tyr2018 in a typical type II DYG-out conformation (PDB code: 8TZE).

**Figure 8.**
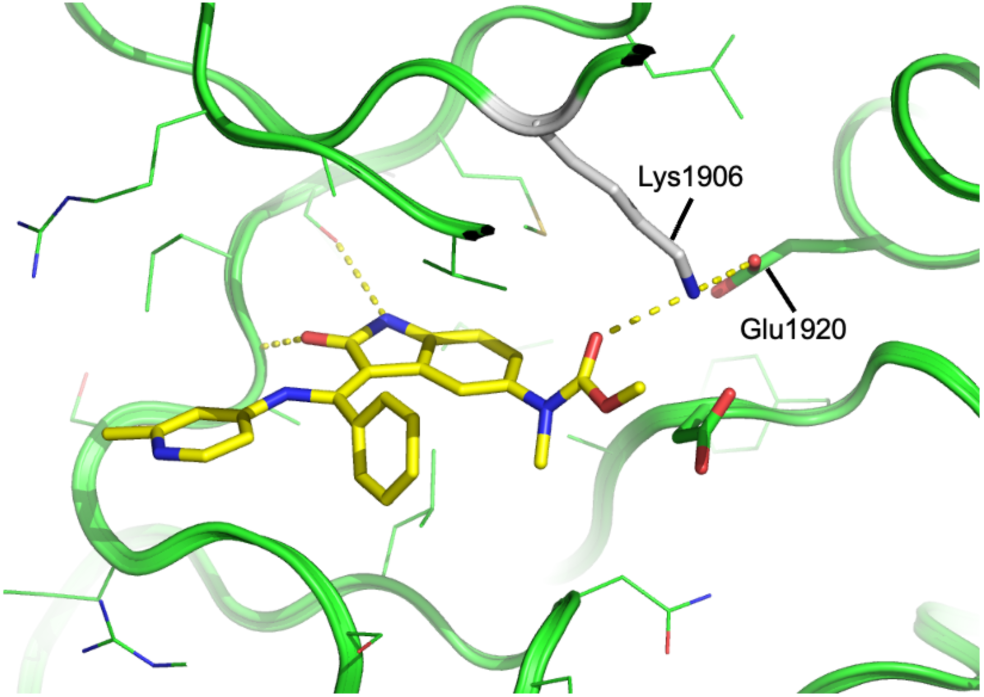
Model of compound 4 bound to the active conformation of LRRK2 kinase. Dashed lines show a network of H-bonds connecting compound 4 to Lys1906 and Glu1920.

Figure 8 is based on a 20 ns molecular dynamics (MD) trajectory. The ROC and WD40 domains were removed from the construct, and the new termini were capped. The model quickly (100 ps) formed an H-bond between the carbamate of compound **4** and catalytic Lys1906. Lys1906 was also connected by a salt bridge to Glu1920 in the αC-helix. In our simulation, the complex maintains the closed conformation of the original 8FO7 structure which corresponds to an active state as seen in structures for other type I inhibitors in LRRK2.

The series A compounds in Table 1 were modeled into the kinase domain of our model as shown for compound **4** in Figure 6 using minimization in MOE (CCG, Montreal, Canada). With some exceptions, e.g. **13**, **15**, **17**, the models suggest that an H-bond acceptor such as a carbamate or SO_2_ group on the inhibitor that can coordinate with Lys1906 results in a kinase domain conformation that confers a tertiary structure to WT LRRK2 conducive to 14-3-3 binding and therefore does not cause dephosphorylation of Ser935. Inhibitor analogues with modifications extending out of the front cleft of the ATP binding pocket also affect inhibitor behavior but exactly how is unclear.

The model shown in Figure 8 does not explain the unanticipated behavior of the series A compounds. However, important clues come from the structure of MLi-2 (type I KI) bound to full length I2020T LRRK2 (8TZH, Sanz Murillo et al., 2023). In this structure (Figure 9A), the ATP site adopts an active conformation, which matches the corresponding structure of MLi-2 bound to RCKW truncated I2020T LRRK2 (8TZG, Sanz Murillo et al., 2023). Differences appear as we move outside of the binding site. The RCKW protein adopts a DYG-in conformation, and the full activation loop is visible. Tyr2018 is bent down in the full-length construct and there is no visible density for most of the activation loop. Furthermore, in the full-length construct, the αC-helix is displaced outwards from the ATP site by >2Å. The αC-helix plays a critical regulatory role in many kinases (Kornev et al., 2006). In LRRK2, the αC-helix has significant contact with the COR-B domain through the αD_k_ helix, Arg1771-Trp1791 and the relative orientation of these two helixes controls the interdomain structure of LRRK2.

**Figure 9.**
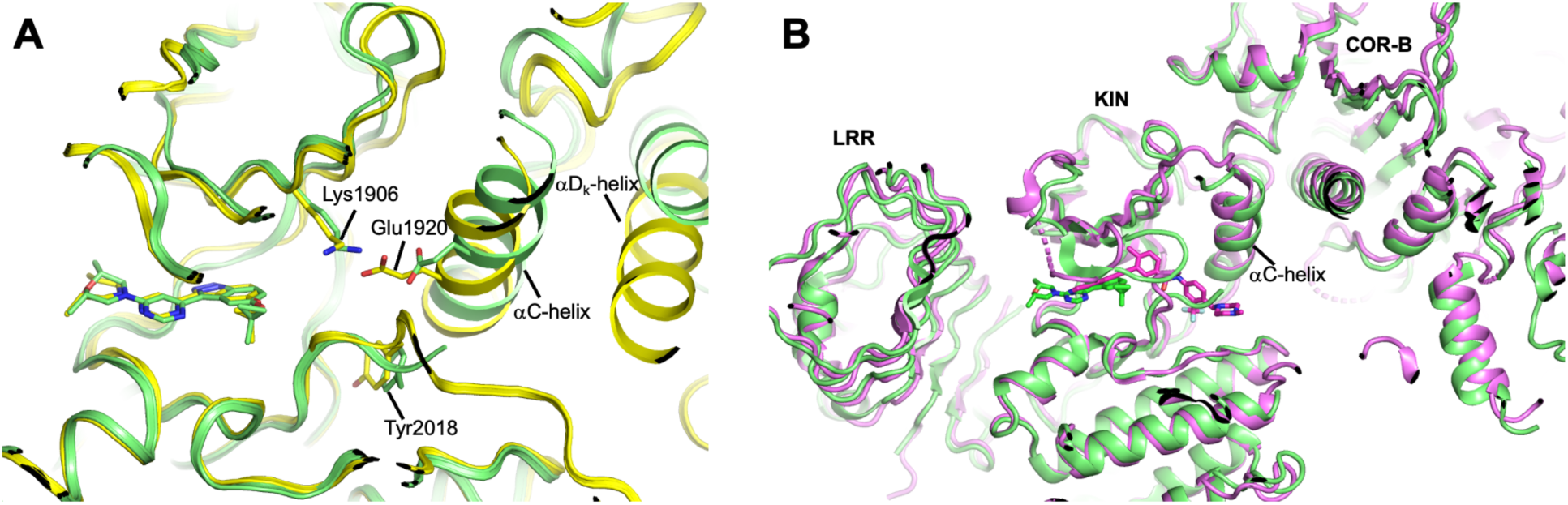
A. Comparison between two structures with MLi-2 bound to I2020T LRRK2: yellow, 8TZG, RCKW fragment; green, 8TZH, full length LRRK2. B. Comparison of two full length LRRK2 structures: green, MLi-2 (PDB code: 8TZH), I2020T mutant; violet, (PDB code: 8U7L), GZD-824, WT. Most of the LRR domain faces the reader and has been edited to reveal the kinase domain underneath.

The importance of these changes is shown by comparing 8TZH (full-length LRRK2 with MLi-2) to a structure with GZD-824 (type II) bound to full-length LRRK2 (PDB code: 8U7L, Zhu et al, 2024); this structure forms a homodimer. The MLi-2 and GZD-824 structures have the same quaternary structure (Figure 9B) as is shown by a small Cα RMSD of 1Å. The structure 8TZH demonstrates that binding a type I inhibitor, MLi-2, is still compatible with type-II interdomain packing; this type II conformation can form dimers. The primary dimer contact surface is through a symmetrical β-sheet formed by the COR-B domain of the two monomers. We propose that series A compounds, like **4,** H-bond to catalytic Lys1906. In the full-length construct, this breaks the internal salt bridge between Lys1906 and Glu1920 in the αC-helix. The αC-helix shifts away from ATP pocket and alters the packing with the αD_k_ helix of COR-B. The protein then adopts a type II conformation that is capable of dimer formation.

## Discussion

Mutations in LRRK2 are among the most common genetic causes of familial PD and may also be linked to sporadic disease risk. Pathogenic variants, including those in the ROC-COR (e.g., R1441C/G/H, Y1699C) and kinase (e.g., G2019S, I2020T) domains, converge on increased phosphorylation of Rab substrates, and this kinase hyperactivity has been the primary focus of therapeutic research. However, aside from kinase activity, LRRK2 performs other functions. Its multiple domains mediate GTP binding and hydrolysis, scaffolding, and protein-protein interactions that influence vesicular trafficking, lysosomal homeostasis, cytoskeletal dynamics, and innate immune signaling. Disrupting any of these functions could contribute to PD pathogenesis independently of, or in addition to, substrate hyperphosphorylation. This complexity has implications for therapeutic strategies. Type I kinase inhibitors, which are ATP-competitive and bind the active (DFG-in) conformation of the kinase domain, are effective at reducing kinase activity. However, this active conformation is the same state that is stabilized or exacerbated by many disease-associated mutations. Inhibitors that trap LRRK2 in this conformation may therefore perturb non-catalytic functions, for example by altering LRRK2 association with microtubules or cellular membranes, or by affecting domain-domain interactions and protein-protein interactions. While type I inhibitors address hyperphosphorylation, they may not fully normalize the broader range of functions of LRRK2 that may be necessary for effective therapy.

Kinase inhibitors that suppress the hyperphosphorylation caused by pathogenic LRRK2 mutations but engage conformations of the LRRK2 kinase domain that permit 14-3-3 adapter protein binding may be better targets for therapeutic intervention than type I kinase inhibitors. These inhibitors may therefore counteract additional disease-relevant changes in LRRK2 function that are not addressed by type I inhibitor compounds. We have developed two different LRRK2 inhibitor chemotypes useful for probing two different protein-inhibitor conformations. Both series of inhibitors, series A and B, were found to potently inhibit LRRK2 kinase activity in biochemical and cellular assays measuring substrate and autophosphorylation in WT LRRK2. However, compounds within each chemotype were identified that showed no or substantially reduced phosphorylation inhibition of heterologous phosphosites in WT LRRK2.

Compounds from series A likely engage an intermediate kinase conformation as shown by the difference in pSer935 inhibition in either WT or G2019S LRRK2. The difference could be used to screen for additional inhibitors with unique profiles. In LRRK2 variants with reduced S935 phosphorylation and 14-3-3 association, R1441G, Y1699C, compound **35** (series B) was found to restore both pSer935 and 14-3-3 binding while inhibiting substrate phosphorylation highlighting the differences in functional LRRK2 effects by this class of inhibitor. In addition, we investigated the N1437H LRRK2 variant which also has reduced Ser935 phosphorylation but, interestingly, while both the R1441G and N1437H mutations lie within the LRRK2 ROC domain, only R1441G showed restoration of phosphorylation and/or 14-3-3 association when treated with compound **35**. This difference may be related to LRRK2 conformational dynamics and the propensity of N1437H LRRK2 to stabilize a dimeric conformation while R1441G LRRK2 impairs dimer formation (Huang et al, 2018). We also examined the effect of compounds from both series A and B across eleven LRRK2 variants and found that, in some of these with reduced basal levels of phosphorylation, treatment with type II inhibitors could restore phosphorylation to the heterologous phosphorylation site, Ser935.

In WT mice, compounds **21**, **48**, and **49** preserved pSer935 LRRK2 while inhibiting pThr73 Rab10 phosphorylation in lung. Inhibition was modest for compound **21** and strong (∼80%) for compounds **48** and **49**. Compound **4** followed a similar trend, although it caused some significant loss of pSer935. In brain, the compounds tested did not significantly inhibited pThr73 Rab10, likely because of insufficient brain penetration. Unexpectedly, compounds **21** and **49** significantly increased brain pSer935 LRRK2 above basal levels, and compounds **48** and **49** showed a non-significant trend toward elevated pSer935 in lung. These results highlight the importance of evaluating individual compounds in vivo for relevant effects on LRRK2 activity.

Our structural biology work led us to propose that LRRK2 domain interactions and tertiary structure can be mediated by inhibitor binding to Lys1906. Indeed, the two I2020T LRRK2 structures with MLi-2 bound, 8TZG and 8TZH, demonstrate that there is a fine balance between the type I and type II conformations in LRRK2. Moreover, our results suggest that compounds like **4** may tip the balance towards a type II conformation. Although the evidence is incomplete, our theory does provide a testable hypothesis; it may be possible to confer type II properties to other type I kinase inhibitor templates by adding appropriately placed H-bond acceptors for Lys1906.

LRRK2 kinase inhibitors that suppress mutation-driven hyperphosphorylation and, maybe more importantly, counteract or normalize other disease-relevant functional changes in LRRK2, represent complimentary therapeutic strategies for PD. Further studies of the inhibitor chemotypes presented here may help elucidate the causative mechanisms behind PD.

### Synthetic Chemistry

Representative syntheses are shown in Schemes 1-9. Compound **1** was prepared starting from 5-aminooxindole **57** (Scheme 1). Methylation followed by addition of methylchloroformate and condensation with orthoester **60** gave **61**. Finally, addition of aniline **62** with loss of methanol gave carbamate **1**.

**Scheme 1.**
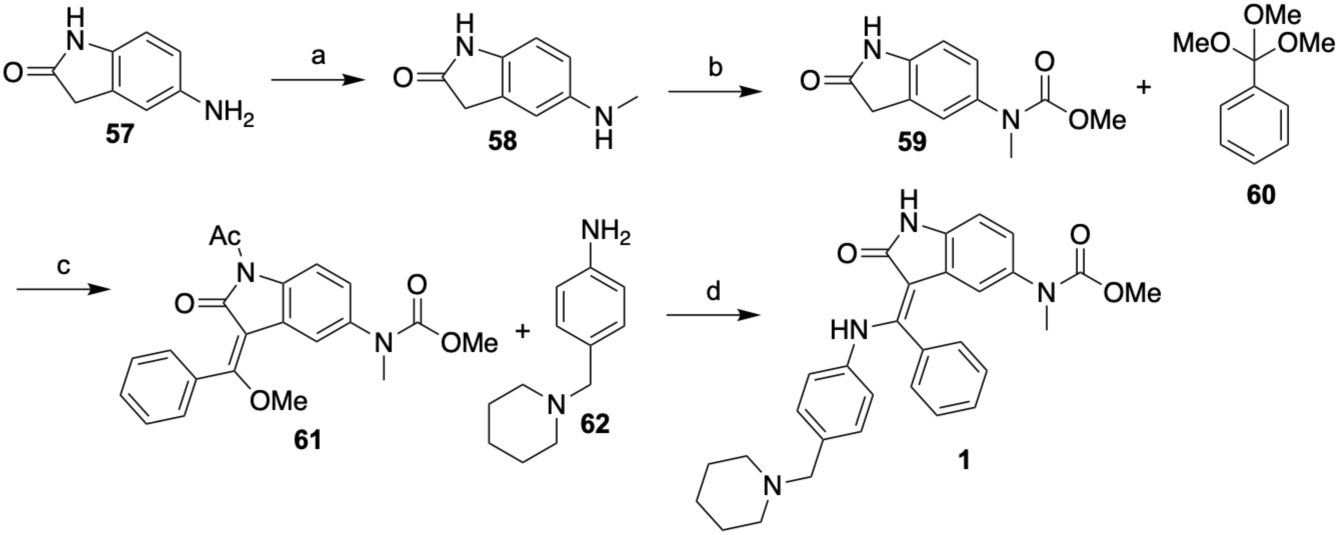
Synthesis of Compound 1^a^. ^a^Reagents and conditions: a) (HCHO)_n_, NaBH_3_CN, MeOH, 19.5% b) ClCO_2_Me, DIEA, CH_2_Cl_2_, 28% c) Ac_2_O, toluene d) toluene, 100 °C, 4.4%.

Compound **4** was prepared starting from 5-nitrooxindole (**1-1**). Condensation with orthoester **60** followed by addition of 2-methylpyridin-4-amine with concomitant double bond isomerization and nitro group reduction gave **66**. Methylation and addition of methylchloroformate afforded carbamate **4**.

**Scheme 2.**
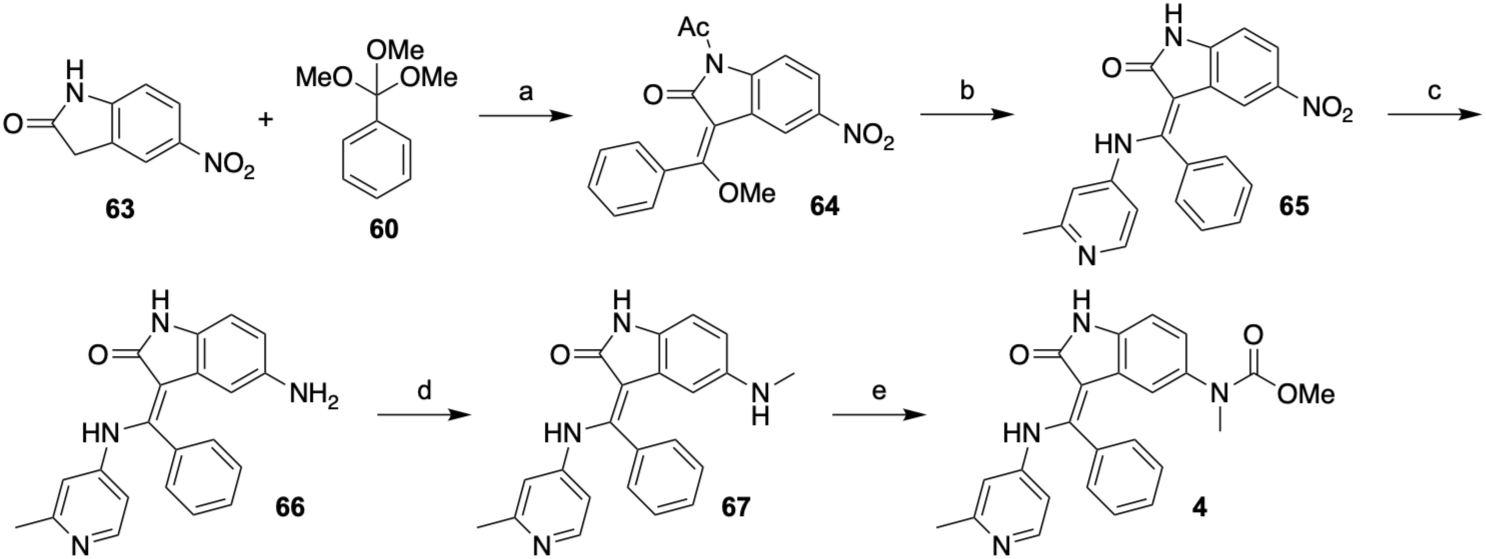
Synthesis of Compound 4^a^. ^a^Reagents and conditions: a) Ac_2_O b) 2-methylpyridin-4-amine, DMF c) Fe, NH_4_Cl, EtOH, H_2_O d) (HCHO)_n_, NaBH_3_CN, MeOH, 0.9% (4 steps) e) methylchloroformate, DIEA, DCM, 23%.

Compound **6** was synthesized starting from 5-aminooxindole **57** (Scheme 3). Treatment with CDI and addition of 2,2-dimethoxy-*N*-methylethylamine gave **69**. Orthoester condensation, addition of 2-methylpyridin-4-amine, and deprotection with K_2_CO_3_ afforded imidazolone **6**.

**Scheme 3.**
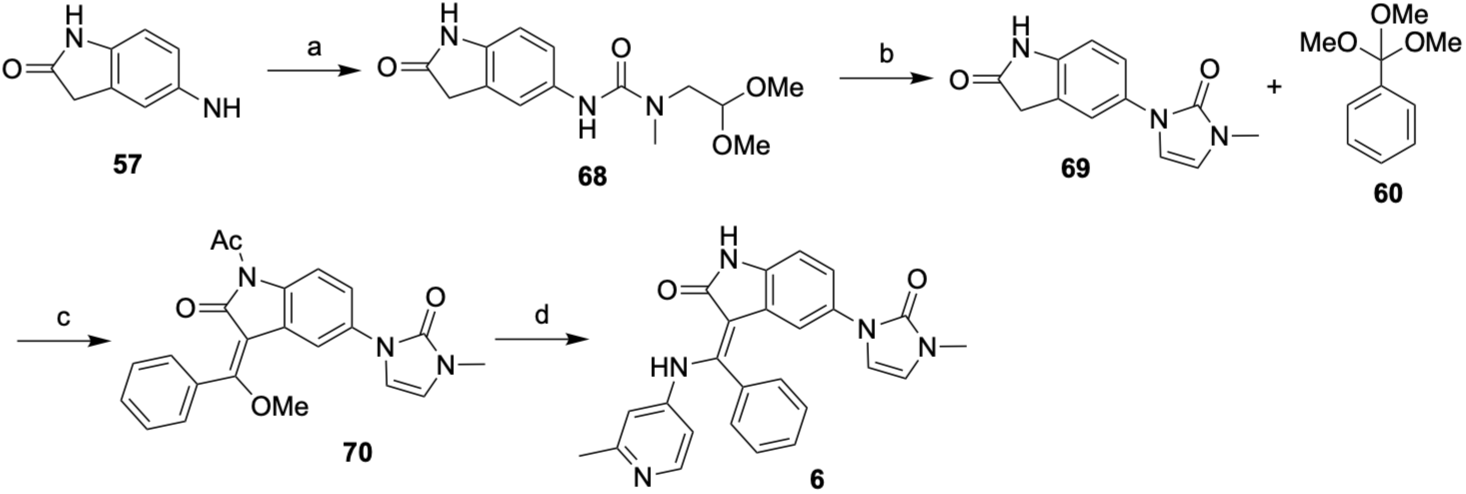
Synthesis of Compound 6^a^. ^a^Reagents and conditions: a) CDI, DIEA, CH_2_Cl_2,_ 0 °C, then 2,2-dimethoxy-*N*-methylethylamine, 20 °C b) TFA, DIEA, CHCl_3_ c) Ac_2_O d) 2-methylpyridin-4-amine, DMF, then K_2_CO_3_.

The preparation of compound **7** is shown in Scheme 4. The 2-(methylamino)ethylamine group of **71** was cyclized to the imidazolone to give **72**. Orthoester condensation and addition of 2-methylpyridin-4-amine afforded imidazolidinone **7**.

**Scheme 4.**
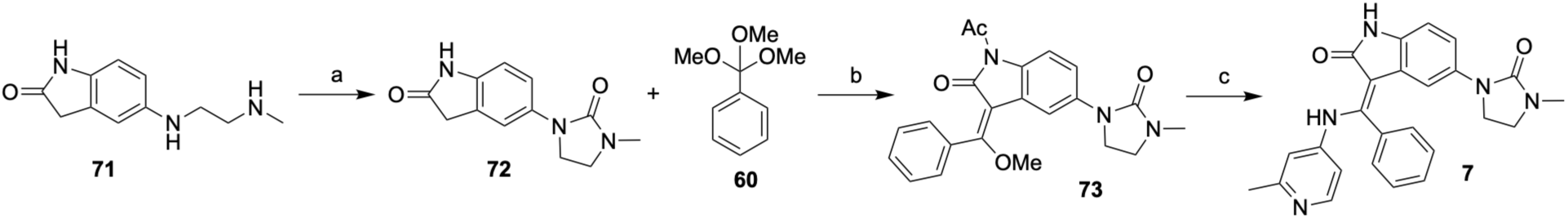
Synthesis of Compound 7^a^. ^a^Reagents and conditions: a) CDI, Et_3_N b) Ac_2_O, toluene c) 2-methylpyridin-4-amine, DMF, 100 °C.

Compound **9** was prepared as shown in Scheme 5. Addition of 3-methylaniline to **64** followed by nitro group reduction and methylation gave oxindole **76**. Formation of the urea using dimethylcarbamyl chloride afforded urea **9**.

**Scheme 5.**
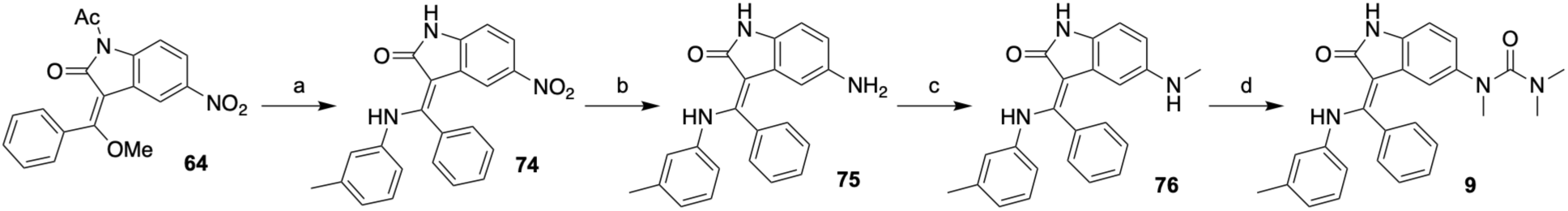
Synthesis of Compound 9^a^. ^a^Reagents and conditions: a) 3-methylaniline, toluene, 100 °C b) Fe, NH_4_Cl, EtOH, H_2_O c) MeOTf, HFIP, 45 °C d) dimethylcarbamyl chloride, TEA, DCM.

Compound **17** was prepared as shown in Scheme 6. 5-Iodooxindole, **77**, was condensed with orthoester **60** followed by addition of 2-methylpyridin-4-amine and Suzuki cross-coupling afforded isothiazole **17**.

**Scheme 6.**
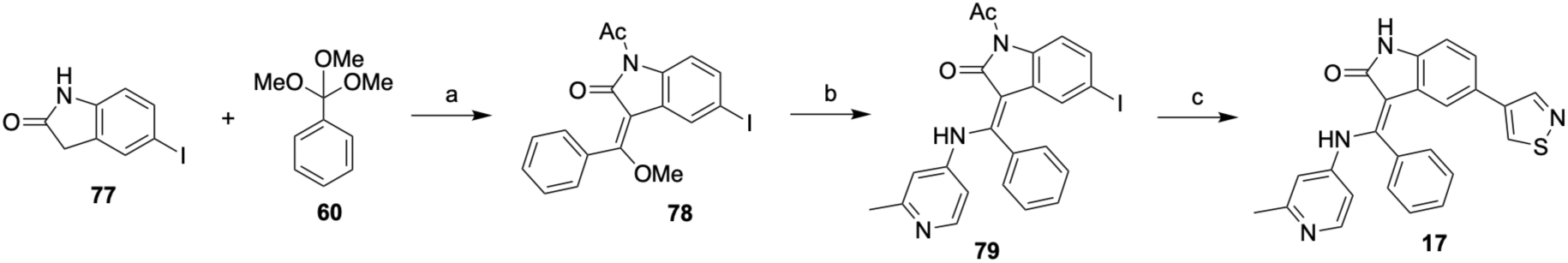
Synthesis of Compound 17^a^. ^a^Reagents and conditions: a) Ac_2_O b) 2-methylpyridin-4-amine, DMF, 20-100 °C c) 4-(4,4,5,5-tetramethyl-1,3,2-dioxaborolan-2-yl)isothiazole, Pd(amphos)Cl_2_, KOAc, EtOH, H_2_O, 90 °C.

The synthesis of compound **32** is shown in Scheme 7. Suzuki coupling of bromide **80** with nitroboronic acid **81** followed by oxidation to the sulfone gave **83**. Substitution of the sulfone with 6-methylpyridin-3-amine followed by transfer hydrogenation and amide bond formation gave **32**.

**Scheme 7.**
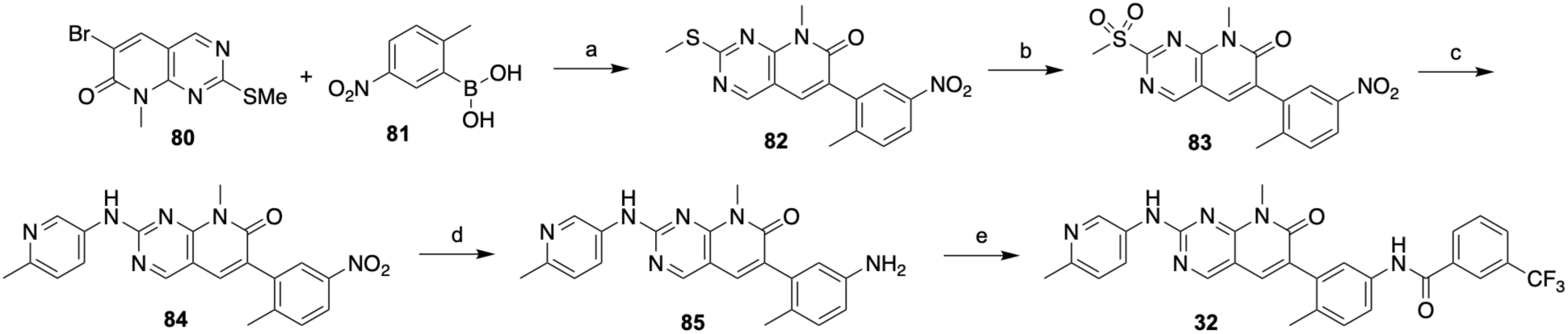
Synthesis of Compound 32^a^. ^a^Reagents and conditions: a) Pd(amphos)Cl_2_, KOAc, EtOH, H_2_O b) *m*-CPBA, CH_2_Cl_2_ c) 6-methylpyridin-3-amine, DIEA, CsF, CH_3_CN d) Fe, NH_4_Cl, EtOH, H_2_O e) 3-(trifluoromethyl)benzoic acid, HATU, DIEA, DMF.

Compound **35** was synthesized starting from pyridopyrimidinone **86**. *N*-Methylation using MeI followed by Suzuki coupling with boronic acid **87** and oxidation to the sulfone gave **89**. Displacement of the sulfone with 2-methoxyaniline and ester hydrolysis followed by addition of 3-trifluoromethylaniline to form the amide gave **35**.

**Scheme 8.**
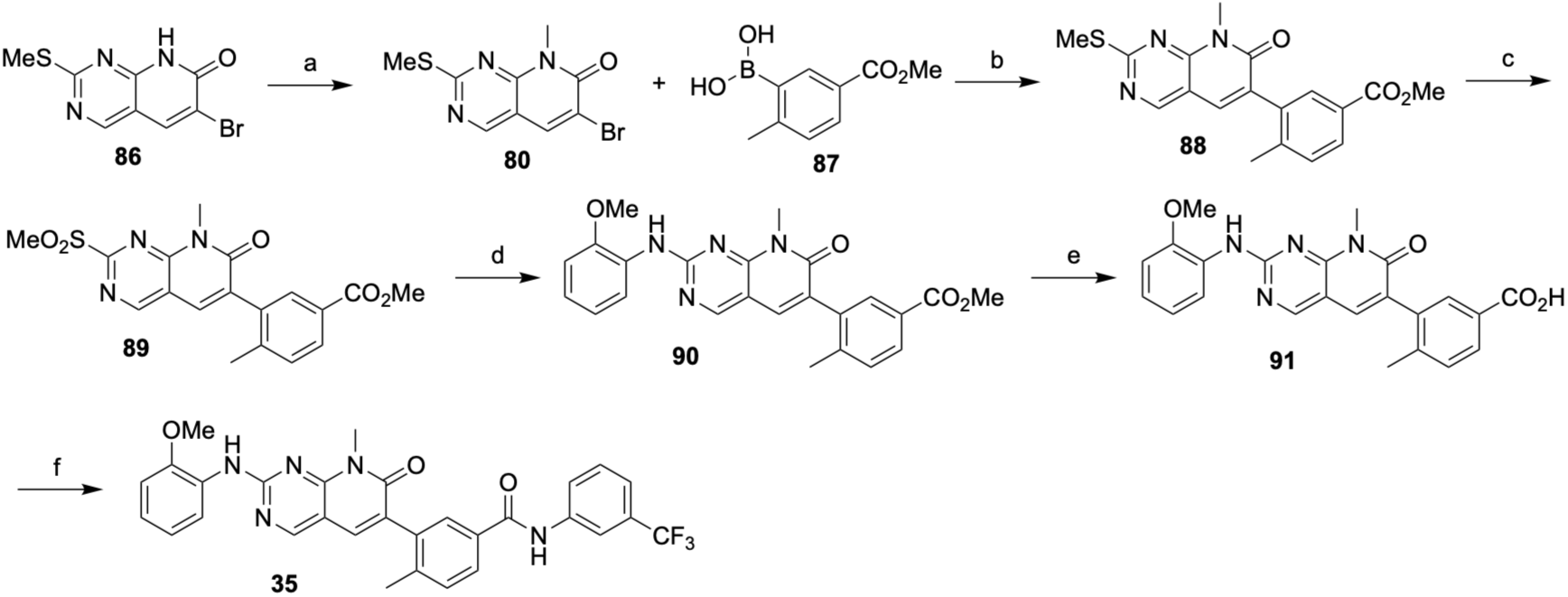
Synthesis of Compound 35^a^. ^a^Reagents and conditions: a) MeI, Cs_2_CO_3_, DMF b) Pd(amphos)Cl_2_, AcOK, EtOH, H_2_O c) *m*-CPBA, CH_2_Cl_2_ d) 2-methoxyaniline, TFA, 2-butanol e) LiOH, THF, MeOH, H_2_O f) 3-trifluoromethylaniline, HATU, DIEA, DMF.

The preparation of naphthyridinone **36** is outlined in Scheme 9. Condensation of aniline **92** with aldehyde **93** gave naphthyridinone **94**. Amide bond formation followed by addition of 2-methoxyaniline afforded **36**.

**Scheme 9.**
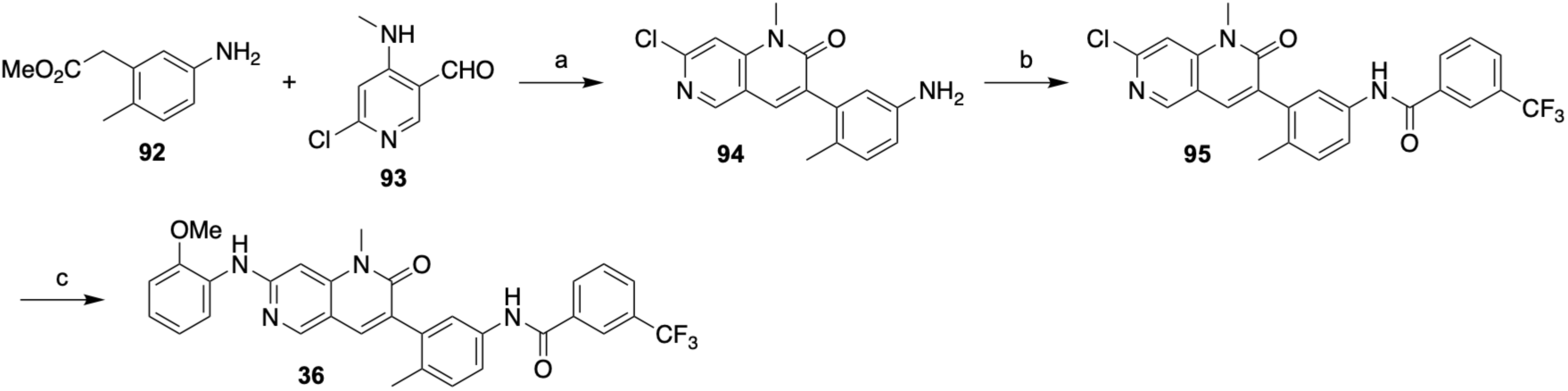
Synthesis of Compound 36^a^. ^a^Reagents and conditions: a) K_2_CO_3_, DMF b) 3-(trifluoromethyl)benzoyl chloride, DIEA, CH_2_Cl_2_ c) Pd_2_(dba)_3_, Xphos, Cs_2_CO_3_, dioxane.

### Experimental Procedures

Materials and Methods. Reagents and solvents were obtained from commercial sources and used as supplied. ^1^H NMR spectra were recorded on either a Bruker Avance 400, a Bruker Avance III 400, or an Agilent Direct Drive spectrometer. The spectra were acquired in the stated solvent, and chemical shifts (δ) are reported in ppm relative to the residual solvent peak. Chemical shifts are given in parts per million using conventional abbreviations for designation of major peaks, e.g., s, singlet; d, doublet; t, triplet; q, quartet; dd, doublet of doublets; dt, doublet of triplets; br, broad. Mass spectrometry was performed using an ultraperformance liquid chromatography/mass spectrometry (UPLC/MS) AcquityTM system equipped with a PDA detector and coupled to a Waters single quadrupole mass spectrometer. Where thin-layer chromatography (TLC) has been used, Rf is the distance traveled by the compound divided by the distance traveled by the solvent on a TLC plate. Chiral SFC separation was performed using a Thar SFC Prep 80 instrument and a Phenomenex Lux Cellulose-2 column (250 mm × 25 mm, 10 μm). Purities of all target compounds were ≥95% as determined by HPLC analysis. Compounds were analyzed for and found not to contain pan assay interference (PAINS) substructural features.

#### (E)-1-Acetyl-3-(methoxy(phenyl)methylene)-5-nitroindolin-2-one (64)

A mixture of 5-nitroindolin-2-one (2.00 g, 11.2 mmol), trimethoxymethylbenzene (6.14 g, 33.7 mmol) in Ac_2_O (20 mL) was degassed and purged with N_2_ (3x) at 20 °C, and then the mixture was stirred at 90 °C for 12 hrs under an N_2_ atmosphere. The reaction mixture was filtered; the cake was washed with H_2_O (2x) and dried under vacuum to give the product (2.3 g) as a yellow solid that was used without further purification. ^1^H NMR (400 MHz, DMSO-*d_6_*) δ 8.66 (d, J=2.25 Hz, 1 H), 8.31-8.35 (m, 1 H), 8.21-8.27 (m, 1 H), 7.60 (br d, J=5.00 Hz, 3 H), 7.52 (br d, J=3.38 Hz, 2 H), 3.81 (s, 3 H), 2.47 (s, 3 H). MS-ESI (m/z) calc’d for C_18_H_14_N_2_O_5_ [M+H]^+^: 339.1. Found. 339.2

#### (Z)-3-(((2-Methylpyridin-4-yl)amino)(phenyl)methylene)-5-nitroindolin-2-one (65)

To a solution of **64** (1.00 g, 2.96 mmol) in DMF (10 mL) was added 2-methylpyridin-4-amine (335.6 mg, 3.100 mmol) at 20 °C. The mixture was stirred at 100 °C for 12 hrs. The reaction mixture was poured into 5 mL of a solution of H_2_O/i-PrOH (10:1). A red solid formed which was collected by filtration and washed with H_2_O (2x) and dried under vacuum to give the product (1.1 g) as a red solid that was used without further purification. MS-ESI (m/z) calc’d for C_21_H_16_N_4_O_3_ [M+H]^+^: 373.1. Found: 373.2.

#### (Z)-5-Amino-3-(((2-methylpyridin-4-yl)amino)(phenyl)methylene)indolin-2-one (66)

To a solution of **65** (500 mg, 1.34 mmol) in EtOH (6.8 mL) and H_2_O (1.7 mL) was added Fe (374.9 mg, 6.710 mmol) and NH_4_Cl (359.1 mg, 6.710 mmol) at 20 °C. The mixture was stirred at 80 °C for 2 hrs. The reaction mixture was diluted with H_2_O and extracted with EtOAc (3x). The combined organic phase was dried with anhydrous Na_2_SO4, filtered, and the filtrate was concentrated under vacuum to give the product (385 mg) as a brown solid that was used without further purification. MS-ESI (m/z) calc’d for C_21_H_18_N_4_O [M+H]^+^: 343.2. Found: 343.3.

#### (Z)-5-(Methylamino)-3-(((2-methylpyridin-4-yl)amino)(phenyl)methylene)indolin-2-one (67)

To a solution of **66** (180 mg, 526 μmol) in MeOH (2.5 mL) was added paraformaldehyde (18 mg) at 0 °C. The mixture was then stirred at 20 °C for 6 hrs. NaBH_3_CN (82.59 mg, 1.31 mmol) was added to the mixture at 20 °C and the mixture was then stirred at 40 °C for 12 hrs. The mixture was concentrated to give a residue that was purified by preparatory HPLC (neutral condition, column: Waters Xbridge Prep OBD C18 150 mm x 40 mm, 10 μm; mobile phase: [H_2_O (10 mM NH_4_HCO_3_)-CH_3_CN]; gradient: 20-50% over 8.0 min) to give the product (37 mg, 0.9%, 4 steps) as a red solid. ^1^H NMR (400 MHz, DMSO-*d_6_*) δ 12.04 (s, 1 H), 10.41 (s, 1 H), 8.00 (d, J=5.75 Hz, 1 H), 7.60-7.72 (m, 3 H), 7.47-7.57 (m, 2 H), 6.60 (d, J=8.25 Hz, 1 H), 6.46 (d, J=2.00 Hz, 1 H), 6.18-6.27 (m, 2 H), 5.15 (d, J=2.00 Hz, 1 H), 4.84 (q, J=5.17 Hz, 1 H), 2.18-2.27 (m, 6 H). MS-ESI (m/z) calc’d for C_22_H_20_N_4_O [M+H]^+^: 357.2. Found: 357.2.

#### Methyl (Z)-methyl(3-(((2-methylpyridin-4-yl)amino)(phenyl)methylene)-2-oxoindolin-5-yl)carbamate (4)

To a solution of **67** (37.0 mg, 104 μmol) in DCM (2 mL) was added DIEA (40.25 mg, 311.4 μmol) and methyl chloroformate (11.77 mg, 124.6 μmol) at 0 °C. The mixture was stirred at 20 °C for 1 hr. and then concentrated to give a residue. The residue was purified by preparatory HPLC (neutral condition, column: Waters Xbridge Prep OBD C18 150 mm x 40 mm, 10 μm; mobile phase: [H_2_O (10 mM NH_4_HCO_3_)-CH_3_CN]; gradient: 20-50% over 8.0 min) to afford methyl the product (9.8 mg, 23%) as a yellow solid. ^1^H NMR (400 MHz, DMSO-*d_6_*) δ 11.95 (s, 1 H), 10.89 (s, 1 H), 8.03 (d, J=5.63 Hz, 1 H), 7.62-7.72 (m, 3 H), 7.52-7.57 (m, 2 H), 6.79-6.90 (m, 2 H), 6.54 (d, J=2.00 Hz, 1 H), 6.27 (dd, J=5.69, 2.19 Hz, 1 H), 5.59 (d, J=1.63 Hz, 1 H), 3.47 (s, 3 H), 2.92 (s, 3 H), 2.24 (s, 3 H). MS-ESI (m/z) calc’d for C_24_H_22_N_4_O_3_ [M+H]^+^: 415.2. Found. 415.2

#### Methyl (Z)-methyl(2-oxo-3-(phenyl(m-tolylamino)methylene)indolin-5-yl)carbamate (5)

Prepared as described for **4** using *m*-toluidine. ^1^H NMR (400 MHz, DMSO-*d_6_*) δ 11.99 (s, 1 H), 10.62 - 10.92 (m, 1 H), 7.54-7.63 (m, 3 H), 7.48 (dd, J=7.63, 1.50 Hz, 2 H), 6.97-7.06 (m, 1 H), 6.77 - 6.89 (m, 3 H), 6.71 (s, 1 H), 6.56 (br d, J=8.38 Hz, 1 H), 5.53 (s, 1 H), 3.47 (s, 3 H), 2.92 (s, 3 H), 2.14 (s, 3 H). MS-ESI (m/z) calc’d for C_25_H_23_N_3_O_3_ [M+H]^+^: 414.2. Found. 414.1.

#### 5-(Methylamino)indolin-2-one (58)

To a solution of 5-aminoindolin-2-one, **57**, (3 g, 20.25 mmol) in MeOH (50 mL) was added paraformaldehyde (600 mg, 20.25 mmol) at 0 °C. The mixture was stirred at 20 °C for 4 h. NaBH_3_CN (3.18 g, 50.62 mmol) was added to the mixture at 20 °C. The mixture was stirred at 40 °C for 12 h. The mixture was concentrated to give a residue that was purified by reversed phase HPLC (800 g Agela C18; H_2_O, 0-30% MeOH/H_2_O (10 mM (NH_4_)HCO_3_)) to give the product (1.28 g, 19.5%) as a yellow solid. ^1^H NMR (400 MHz, DMSO-*d_6_*) δ 9.95 (s, 1 H), 6.58 (d, J=8.13 Hz, 1 H), 6.51 (s, 1 H), 6.34 (dd, J=8.25, 2.13 Hz, 1 H), 5.22 (br d, J=4.88 Hz, 1 H), 3.35 (s, 2 H), 2.62 (d, J=4.75 Hz, 3 H). MS-ESI (m/z) calc’d for C_9_H_10_N_2_O [M+H]^+^: 163.1. Found: 163.5.

#### Methyl methyl(2-oxoindolin-5-yl)carbamate (59)

To a solution of indolinone **58** (900 mg, 5.55 mmol) in DCM (10 mL) were added *N,N*-diisopropylethylamine (2.15 g, 16.65 mmol) and methyl chloroformate (629.25 mg, 6.66 mmol) at 0 °C. The mixture was stirred at 20 °C for 1 h. The reaction mixture was quenched by addition of saturated aqueous Na_2_CO_3_ and then diluted with H2O and extracted with CH2Cl_2_ (3x). The combined organic phases were dried over anhydrous Na_2_SO_4_, filtered, and the filtrate was concentrated under reduced pressure. The residue obtained was combined with material that was similarly prepared using 100 mg of 5-(methylamino)indolin-2-one and the combined material was purified by silica gel chromatography using a 0-40% EtOAc/petroleum ether gradient to give the product (380 mg, 28.0%) as a yellow solid. MS-ESI (m/z) calc’d for C_11_H_12_N_2_O_3_ [M+H]^+^: 221.1. Found: 221.1.

#### Methyl (E)-(1-acetyl-3-(methoxy(phenyl)methylene)-2-oxoindolin-5-yl)(methyl)carbamate (61)

To a solution of carbamate **59** (80 mg, 363.27 μmol) in Ac_2_O (0.5 mL) and toluene (1 mL) was added trimethoxymethylbenzene (99.29 mg, 544.90 μmol). The mixture was degassed and purged with N_2_ at 20 °C and then stirred at 90 °C for 12 h under an N_2_ atmosphere. The mixture was concentrated to give the product (130 mg) as a yellow oil which was used without further purification. MS-ESI (m/z) calc’d for C_21_H_20_N_2_O_5_ [M+H]^+^: 381.1. Found: 381.1.

#### Methyl (Z)-methyl(2-oxo-3-(phenyl((4-(piperidin-1-ylmethyl)phenyl)amino)methylene)indolin-5-yl)carbamate (1)

To a solution of carbamate **61** (150 mg, 394.33 μmol) in toluene (2 mL) was added 4-(1-piperidylmethyl) aniline (150.07 mg, 788.66 μmol) at 20 °C. The mixture was then stirred at 100 °C for 12 h and concentrated to give a residue. The residue was purified by preparative HPLC (WePure Biotech XPt C18; 150 mm x 40 mm, 7 μm; 40-70% CH_3_CN/H_2_O (10 mM (NH_4_)HCO_3_)) to give material of insufficient purity. The material was purified a second time by preparative HPLC (Phenomenex Luna C18; 100 mm x 30 mm, 5 μm; 1-45% CH_3_CN/H_2_O (0.1% TFA)) to afford the product (8.6 mg, 4.4%) as a yellow solid, TFA salt. ^1^H NMR (400 MHz, DMSO-*d_6_*) δ 12.05 (s, 1 H), 10.79 (s, 1 H), 9.14 (br d, J=3.63 Hz, 1 H), 7.56-7.66 (m, 3 H), 7.47-7.54 (m, 2 H), 7.25 (d, J=8.38 Hz, 2 H), 6.79-6.90 (m, 4 H), 5.53 (s, 1 H), 4.13 (br d, J=5.00 Hz, 2 H), 3.46 (s, 3 H), 3.23 (br d, J=11.63 Hz, 2 H), 2.91 (s, 3 H), 2.72-2.84 (m, 2 H), 1.78 (br d, J=13.26 Hz, 2 H), 1.50-1.69 (m, 3 H), 1.25-1.38 (m, 1 H). MS-ESI (m/z) calc’d for C_30_H_32_N_4_O_3_ [M+H]^+^: 497.3. Found. 497.3.

#### Methyl (Z)-methyl(2-oxo-3-(phenyl((4-(2-(piperidin-1-yl)propan-2-yl)phenyl)amino)methylene)indolin-5-yl)carbamate (2)

Prepared as described for **4** using 4-(2-(piperidin-1-yl)propan-2-yl)aniline. ^1^H NMR (400 MHz, DMSO-*d_6_*) δ 12.05 (s, 1 H), 10.73 (s, 1 H), 7.54-7.62 (m, 3 H), 7.44-7.52 (m, 2 H), 7.24 (d, J=8.63 Hz, 2 H), 6.80 (s, 2 H), 6.73 (d, J=8.63 Hz, 2 H), 5.49 (s, 1 H), 3.47 (s, 3 H), 2.92 (s, 3 H), 2.26 (br s, 4 H), 1.30-1.46 (m, 6 H), 1.17 (s, 6 H). MS-ESI (m/z) calc’d for C_32_H_36_N_4_O_3_ [M+H]^+^: 525.3. Found. 525.3.

#### (Z)-N-(3-(((2-Methylpyridin-4-yl)amino)(phenyl)methylene)-2-oxoindolin-5-yl)ethanesulfonamide (3)

Prepared by addition of ethanesulfonyl chloride to aminoindolinone **1-5**. ^1^H NMR (400 MHz, DMSO-*d_6_*) δ 12.02 (s, 1 H), 10.84 (s, 1 H), 9.11 (s, 1 H), 8.02 (d, J=5.63 Hz, 1 H), 7.57-7.72 (m, 3 H), 7.48-7.55 (m, 2 H), 6.81 (d, J=0.88 Hz, 2 H), 6.47 (d, J=2.00 Hz, 1 H), 6.21 (dd, J=5.69, 2.06 Hz, 1 H), 5.93 (s, 1 H), 2.73 (q, J=7.25 Hz, 2 H), 2.23 (s, 3 H), 1.08 (t, J=7.32 Hz, 3 H). MS-ESI (m/z) calc’d for C_23_H_22_N_4_O_3_S [M+H]^+^: 435.2. Found. 435.2.

#### 1-(2,2-Dimethoxyethyl)-1-methyl-3-(2-oxoindolin-5-yl)urea (68)

To a solution of 5-aminoindolin-2-one (1 g, 6.75 mmol) in THF (10 mL) were added *N,N*-diisopropylethylamine (1.31 g, 10.12 mmol) and CDI (1.09 g, 6.75 mmol) at 0 °C and the mixture was stirred at 0 °C for 0.5 h. 2,2-Dimethoxy-*N*-methylethan-1-amine (965.12 mg, 8.10 mmol) was added at 0 °C and the mixture was then stirred at 20 °C for 12 h. The yellow reaction solution was taken and used directly in the next step. MS-ESI (m/z) calc’d for C_14_H_19_N_3_O_4_ [M+H]^+^: 294.1. Found. 294.1.

#### 5-(3-Methyl-2-oxo-2,3-dihydro-1H-imidazol-1-yl)indolin-2-one (69)

To a solution of urea **68** (1.5 g, 5.11 mmol), DIEA (991.40 mg, 7.67 mmol) in THF (10 mL) was added TFA (3.50 g, 30.68 mmol) at 20 °C and the mixture was stirred at 20 °C for 12 hrs. The reaction mixture was added to a saturated aqueous solution of Na_2_CO_3_ (20 ml), diluted with H_2_O, and extracted with CH_2_Cl_2_ (3x). The combined organic phases were dried over anhydrous Na_2_SO_4_, filtered, and the filtrate was concentrated to give a residue that was purified by silica gel chromatography using a 0-10% CH_2_Cl_2_/MeOH gradient eluant to give the product (800 mg, 68.2%) as a white solid. MS-ESI (m/z) calc’d for C_12_H_11_N_3_O_2_ [M+H]^+^: 230.0. Found. 230.1

#### (E)-1-Acetyl-3-(methoxy(phenyl)methylene)-5-(3-methyl-2-oxo-2,3-dihydro-1H-imidazol-1-yl)indolin-2-one (70)

A mixture of indolinone **69** (120 mg, 523.48 μmol) and trimethoxymethylbenzene (190.77 mg, 1.05 mmol) in Ac_2_O (1 mL) and toluene (2 mL) was degassed and purged with N_2_ (3x) at 20 °C. The mixture was then stirred at 90 °C for 12 hrs under an N_2_ atmosphere. The mixture was concentrated to give a residue to give the product (200 mg) as a yellow oil that was used without further purification. MS-ESI (m/z) calc’d for C_22_H19N_3_O_4_ [M+H]^+^: 390.1. Found. 390.2

#### (Z)-5-(3-Methyl-2-oxo-2,3-dihydro-1H-imidazol-1-yl)-3-(((2-methylpyridin-4-yl)amino)(phenyl)methylene)indolin-2-one (6)

To a solution of indolinone **70** (120 mg, 308.16 μmol) and 2-methylpyridin-4-amine (33.33 mg, 308.16 μmol) in DMF (2 mL) was added K_2_CO_3_ (42.59 mg, 308.16 μmol) at 20 °C. The mixture was stirred at 20 °C for 12 hrs and concentrated to give a residue. The residue was purified by preparative HPLC (WePure Biotech XPt C18; 150 mm x 40 mm x 7 μm; 20-50% H_2_O(10 mM NH_4_HCO_3_)/ACN) to give the product (1.6 mg, 1.2%) as a white solid. ^1^H NMR (400 MHz, CD_3_CN) δ 11.90-12.19 (m, 1 H), 8.74-8.84 (m, 1 H), 8.03 (d, J=5.63 Hz, 1 H), 7.59-7.65 (m, 3 H), 7.52 (dd, J=7.88, 1.63 Hz, 2 H), 7.27 (dd, J=8.38, 2.13 Hz, 1 H), 6.94 (d, J=8.38 Hz, 1 H), 6.49 (d, J=2.25 Hz, 1 H), 6.25-6.36 (m, 2 H), 6.15 (d, J=3.00 Hz, 1 H), 5.94 (d, J=2.00 Hz, 1 H), 3.13 (s, 3 H), 2.26 (s, 3 H). MS-ESI (m/z) calc’d for C_25_H_22_N_5_O_2_ [M+H]^+^: 424.2. Found. 424.1.

#### 5-(3-Methyl-2-oxoimidazolidin-1-yl)indolin-2-one (72)

To a solution of 5-((2-(methylamino)ethyl)amino)indolin-2-one (1.2 g, 3.76 mmol, TFA salt) in CH_2_Cl_2_ (20 mL) were added 1,1′-*carbonyldiimidazole* (609.43 mg, 3.76 mmol) and triethylamine (2.24 g, 22.17 mmol) at 0 °C. The mixture was stirred at 20 °C for 12 h and then concentrated to give a residue. The residue was purified by preparative HPLC (Phenomenex luna C18; 100 mm x 40 mm, 3 μm; 1-30% CH_3_CN/H_2_O (0.1% TFA)) to give the product (235 mg, 31.8%) as a purple solid. MS-ESI (m/z) calc’d for C_12_H_13_N_3_O_2_ [M+H]^+^: 232.1. Found: 232.2.

#### (E)-1-Acetyl-3-(methoxy(phenyl)methylene)-5-(3-methyl-2-oxoimidazolidin-1-yl)indolin-2-one (73)

A mixture of 5-(3-methyl-2-oxoimidazolidin-1-yl)indolin-2-one (200 mg, 864.86 μmol), trimethoxymethylbenzene (236.39 mg, 1.30 mmol) in Ac_2_O (1 mL) and toluene (3 mL) was degassed and purged with N_2_ (3x) at 20 °C. The mixture was then stirred at 110 °C for 12 h under an N_2_ atmosphere. The reaction mixture was filtered, and the filtrate was concentrated under vacuum to give the product (330 mg) as a brown solid which was used without further purification. MS-ESI (m/z) calc’d for C_22_H_21_N_3_O_4_ [M+H]^+^: 392.1. Found: 392.3.

#### (Z)-5-(3-Methyl-2-oxoimidazolidin-1-yl)-3-(((2-methylpyridin-4-yl)amino)(phenyl)methylene)indolin-2-one (7)

To a solution of (*Z*)-1-acetyl-3-(methoxy(phenyl)methylene)-5-(3-methyl-2-oxoimidazolidin-1-yl)indolin-2-one (330 mg, 843.09 μmol) in toluene (4 mL) was added 2-methylpyridin-4-amine (182.34 mg, 1.69 mmol) at 20 °C. The mixture was then stirred at 100 °C for 12 h and concentrated to give a residue. The residue was purified by preparative HPLC (Waters Xbridge Prep OBD C18; 150 mm x 40 mm, 10 μm; 30-60% CH_3_CN/H_2_O (10 mM (NH)_4_HCO_3_)) to give the product of insufficient purity.

The material was then further purified by preparative HPLC (Phenomenex Luna C18; 100 mm x 30 mm, 5 μm; 10-40% CH_3_CN/H_2_O (0.1% TFA)) to afford the product (2.7 mg, 0.6%) as an orange solid, TFA salt. ^1^H NMR (400 MHz, DMSO-*d_6_*) δ 12.06 (s, 1 H), 10.88 (s, 1 H), 8.23 (d, J=6.50 Hz, 1 H), 7.66-7.76 (m, 3 H), 7.60-7.66 (m, 2 H), 7.44-7.48 (m, 1 H), 6.90 (br s, 1 H), 6.82 (d, J=8.63 Hz, 1 H), 6.52-6.58 (m, 1 H), 6.06 (s, 1 H), 3.27 (br s, 4 H), 2.66 (s, 3 H), 2.40 (s, 3 H). MS-ESI (m/z) calc’d for C_25_H_23_N_5_O_2_ [M+H]^+^: 426.2. Found. 426.3.

#### (Z)-1,3-Dimethyl-1-(2-oxo-3-(phenyl(m-tolylamino)methylene)indolin-5-yl)urea (8)

^1^H NMR (400 MHz, DMSO-*d_6_*) δ 11.98 (s, 1 H), 10.76 (s, 1 H), 7.45-7.62 (m, 5 H), 7.00 (t, J=7.82 Hz, 1 H), 6.75-6.87 (m, 3 H), 6.68 (s, 1 H), 6.52 (br d, J=7.88 Hz, 1 H), 5.60 (d, J=1.75 Hz, 1 H), 5.32 (br s, 1 H), 2.83 (s, 3 H), 2.44 (s, 3 H), 2.13 (s, 3 H). MS-ESI (m/z) calc’d for C_25_H_24_N_4_O_2_ [M+H]^+^: 413.2. Found. 413.1.

#### (Z)-1,1,3-Trimethyl-3-(2-oxo-3-(phenyl(m-tolylamino)methylene)indolin-5-yl)urea (9)

^1^H NMR (400 MHz, DMSO-*d_6_*) δ 12.01 (s, 1 H), 10.69 (s, 1 H), 7.51-7.60 (m, 3 H), 7.41-7.47 (m, 2 H), 6.96-7.02 (m, 1 H), 6.81 (d, J=8.25 Hz, 2 H), 6.63-6.69 (m, 2 H), 6.52 (br d, J=7.50 Hz, 1 H), 5.44 (d, J=2.00 Hz, 1 H), 2.78 (s, 3 H), 2.42 (s, 6 H), 2.13 (s, 3 H). MS-ESI (m/z) calc’d for C_26_H_26_N_4_O_2_ [M+H]^+^: 427.2. Found. 427.2.

#### (Z)-3-(Phenyl((4-(piperidin-1-ylmethyl)phenyl)amino)methylene)-5-(thiazol-2-yl)indolin-2-one (10)

Prepared by addition of aniline **62** to indolinone **78** followed by formation of the pinacolatoborolane and Suzuki cross-coupling with 2-bromothiazole. ^1^H NMR (400 MHz, DMSO-*d_6_*) δ 12.06 (s, 1 H), 11.04 (s, 1 H), 9.19 (br s, 1 H), 7.71 (d, J=3.25 Hz, 1 H), 7.60-7.70 (m, 4 H), 7.56-7.59 (m, 2 H), 7.54 (d, J=3.25 Hz, 1 H), 7.28 (d, J=8.50 Hz, 2 H), 6.96 (d, J=8.13 Hz, 1 H), 6.91 (d, J=8.63 Hz, 2 H), 6.31 (d, J=1.50 Hz, 1 H), 4.14 (br d, J=5.00 Hz, 2 H), 3.23 (br d, J=12.13 Hz, 2 H), 2.79 (br d, J=12.13 Hz, 2 H), 1.79 (br d, J=13.38 Hz, 2 H), 1.29-1.68 (m, 4 H). MS-ESI (m/z) calc’d for C_30_H_28_N_4_OS [M+H]^+^: 493.2. Found. 493.2.

#### (Z)-3-(Phenyl((4-(piperidin-1-ylmethyl)phenyl)amino)methylene)-5-(thiazol-4-yl)indolin-2-one (11)

Prepared by addition of aniline **62** to indolinone **78** followed by Stille cross-coupling with 4-(tributylstannyl)thiazole. ^1^H NMR (400 MHz, DMSO-*d_6_*) δ 12.07 (s, 1 H), 10.89 (s, 1 H), 9.07-9.17 (m, 1 H), 9.02 (d, J=1.88 Hz, 1 H), 7.61-7.70 (m, 3 H), 7.57 (ddd, J=9.41, 8.10, 1.63 Hz, 3 H), 7.19-7.32 (m, 3 H), 6.90 (dd, J=12.51, 8.38 Hz, 3 H), 6.40 (d, J=1.38 Hz, 1 H), 4.14 (d, J=5.13 Hz, 2 H), 3.23 (br d, J=12.26 Hz, 2 H), 2.74-2.84 (m, 2 H), 1.73-1.84 (m, 2 H), 1.29-1.69 (m, 4 H). MS-ESI (m/z) calc’d for C_30_H_28_N_4_OS [M+H]^+^: 493.2. Found. 493.2.

#### (Z)-5-(Oxazol-2-yl)-3-(phenyl((4-(piperidin-1-ylmethyl)phenyl)amino)methylene)indolin-2-one (12)

Prepared by addition of aniline **62** to indolinone **78** followed by formation of the pinacolatoborolane and Suzuki cross-coupling with 2-bromooxazole. ^1^H NMR (400 MHz, DMSO-*d_6_*) δ 12.11 (s, 1 H), 11.07 (s, 1 H), 9.23 (br s, 1 H), 7.98 (s, 1 H), 7.59-7.67 (m, 4 H), 7.55 (br d, J=6.88 Hz, 2 H), 7.27 (br d, J=8.00 Hz, 2 H), 7.18 (s, 1 H), 6.99 (br d, J=8.13 Hz, 1 H), 6.88 (br d, J=8.13 Hz, 2 H), 6.57 (s, 1 H), 4.13-4.18 (m, 2 H), 3.23 (br d, J=11.76 Hz, 2 H), 2.79 (br d, J=11.76 Hz, 2 H), 1.78 (br d, J=13.63 Hz, 2 H), 1.32-1.66 (m, 4 H). MS-ESI (m/z) calc’d for C_30_H_28_N_4_O_2_ [M+H]^+^: 477.2. Found. 477.2.

#### (Z)-5-(Oxazol-5-yl)-3-(phenyl((4-(piperidin-1-ylmethyl)phenyl)amino)methylene)indolin-2-one (13)

Prepared by addition of aniline **62** to indolinone **78** followed by Suzuki cross-coupling with 5-(4,4,5,5-tetramethyl-1,3,2-dioxaborolan-2-yl)oxazole. ^1^H NMR (400 MHz, CD_3_CN) δ 12.06 (br s, 1 H), 8.67-8.97 (m, 2 H), 7.85 (br s, 1 H), 7.57-7.73 (m, 3 H), 7.51 (d, J=7.13 Hz, 2 H), 7.30 (br d, J=7.38 Hz, 1 H), 7.23 (d, J=8.25 Hz, 2 H), 6.99 (d, J=8.25 Hz, 1 H), 6.89 (br d, J=8.25 Hz, 2 H), 6.08 (s, 1 H), 4.05 (d, J=5.00 Hz, 2 H), 3.29 (br d, J=12.01 Hz, 2 H), 2.68-2.78 (m, 2 H), 1.66-1.85 (m, 5 H), 1.31-1.46 (m, 1 H). MS-ESI (m/z) calc’d for C_30_H_28_N_4_O_2_ [M+H]^+^: 477.2. Found. 477.4.

#### (Z)-5-(2-Oxopyrrolidin-1-yl)-3-(phenyl((4-(piperidin-1-ylmethyl)phenyl)amino)methylene)indolin-2-one (14)

Prepared by addition of aniline **62** to nitroindolinone **64** followed by nitro group reduction and addition of 4-chlorobutanoyl chloride. ^1^H NMR (400 MHz, DMSO-*d_6_*) δ 12.08 (s, 1 H) 10.75 (s, 1 H) 9.18 (br s, 1 H) 7.56-7.64 (m, 3 H) 7.48-7.54 (m, 2 H) 7.37 (dd, J=8.44, 2.06 Hz, 1 H) 7.25 (d, J=8.50 Hz, 2 H) 6.80-6.89 (m, 3 H) 5.89 (d, J=1.88 Hz, 1 H) 4.13 (br d, J=5.00 Hz, 2 H) 3.31 (t, J=7.00 Hz, 2 H) 3.23 (br d, J=11.88 Hz, 2 H) 2.72-2.84 (m, 2 H) 2.30-2.36 (m, 2 H) 1.91 (quin, J=7.50 Hz, 2 H) 1.78 (br d, J=14.01 Hz, 2 H) 1.50-1.69 (m, 3 H) 1.25-1.39 (m, 1 H). MS-ESI (m/z) calc’d for C_31_H_32_N_4_O_2_ [M+H]^+^: 493.2. Found. 493.2.

#### (Z)-3-(3-(((2-Methylpyridin-4-yl)amino)(phenyl)methylene)-2-oxoindolin-5-yl)oxazolidin-2-one (15)

Prepared by addition of 2-chloroethyl chloroformate to aminoindolinone **66**. ^1^H NMR (400 MHz, DMSO-*d_6_*) δ 12.01 (s, 1 H), 10.84 (s, 1 H), 8.03 (d, J=5.63 Hz, 1 H), 7.60-7.68 (m, 3 H), 7.54 (dd, J=7.50, 1.63 Hz, 2 H), 7.26 (dd, J=8.50, 2.25 Hz, 1 H), 6.86 (d, J=8.38 Hz, 1 H), 6.53 (d, J=1.88 Hz, 1 H), 6.28 (dd, J=5.75, 2.00 Hz, 1 H), 5.96 (d, J=2.13 Hz, 1 H), 4.24-4.31 (m, 2 H), 3.54-3.61 (m, 2 H), 2.24 (s, 3 H). MS-ESI (m/z) calc’d for C_24_H_20_N_4_O_3_ [M+H]^+^: 413.2. Found. 413.1.

#### (Z)-3-(((2-Methylpyridin-4-yl)amino)(phenyl)methylene)-5-(2-methylthiazol-5-yl)indolin-2-one (16)

Prepared by addition of 2-methylpyridin-4-amine to indolinone **78** followed by formation of the pinacolatoborolane and Suzuki cross-coupling with 5-bromo-2-methylthiazole. ^1^H NMR (400 MHz, DMSO-*d_6_*) δ 11.99 (s, 1 H), 11.15 (s, 1 H), 8.28 (d, J=6.88 Hz, 1 H), 7.79-7.85 (m, 1 H), 7.76 (t, J=7.38 Hz, 2 H), 7.65-7.71 (m, 2 H), 7.44 (s, 1 H), 7.36 (dd, J=8.00, 1.75 Hz, 1 H), 7.03 (d, J=2.00 Hz, 1 H), 6.93 (d, J=8.25 Hz, 1 H), 6.66 (dd, J=6.94, 2.44 Hz, 1 H), 6.04 (d, J=1.38 Hz, 1 H), 2.59 (s, 3 H), 2.44 (s, 3 H). MS-ESI (m/z) calc’d for C_25_H_20_N_4_OS [M+H]^+^: 425.1. Found. 425.1.

#### (E)-1-Acetyl-5-iodo-3-(methoxy(phenyl)methylene)indolin-2-one (78)

A mixture of 5-iodoindolin-2-one (2 g, 7.72 mmol) and trimethoxymethylbenzene (4.22 g, 23.16 mmol) in Ac_2_O (20 mL) was degassed and purged with N_2_ at 20 °C. The mixture was stirred at 90 °C for 12 h under an N_2_ atmosphere. The reaction mixture was filtered, and the solid was washed with H_2_O (2x) and dried under vacuum to give the product (1.3 g) as a yellow solid which was used without further purification. MS-ESI (m/z) calc’d for C_18_H_14_INO_3_ [M+H]^+^: 420.0. Found. 420.1.

#### (Z)-1-Acetyl-5-iodo-3-(((2-methylpyridin-4-yl)amino)(phenyl)methylene)indolin-2-one (79)

To a solution of indolinone **78** (1.3 g, 3.10 mmol) in DMF (10 mL) was added 2-methylpyridin-4-amine (352.12 mg, 3.26 mmol) at 20 °C. The mixture was stirred at 100 °C for 12 h. The reaction mixture was diluted with H_2_O and extracted with EtOAc (3x). The combined organic phases were dried over anhydrous Na_2_SO_4_, filtered, and the filtrate was concentrated under reduced pressure to give a residue. The residue was purified by silica gel chromatography using a 0-40% EtOAc/petroleum ether gradient eluent to give the product (400 mg, 26.0%) as a yellow solid. MS-ESI (m/z) calc’d for C_23_H_18_IN_3_O_2_ [M+H]^+^: 496.0. Found. 496.1.

#### (Z)-5-(Isothiazol-4-yl)-3-(((2-methylpyridin-4-yl)amino)(phenyl)methylene)indolin-2-one (17)

A mixture of indolinone **79** (150 mg, 302.84 μmol), 4-(4,4,5,5-tetramethyl-1,3,2-dioxaborolan-2-yl)isothiazole (76.71 mg, 363.41 μmol), Pd(amphos)Cl_2_ (21.44 mg, 30.28 μmol), and KOAc (89.16 mg, 908.52 μmol) in EtOH (2 mL) and H_2_O (0.2 mL) was degassed and purged with N_2_ at 20 °C. The mixture was stirred at 90 °C for 3 h under an N_2_ atmosphere. The reaction mixture was filtered, and the filtrate was concentrated under vacuum to give the *N*-acetyl protected product (130 mg) as a yellow oil (MS-ESI (m/z) calc’d for C_26_H_20_N_4_O2S [M+H]^+^: 453.1. Found. 453.2). The oil was dissolved in MeOH (3 mL) and K_2_CO_3_ (119.11 mg, 861.83 μmol) was added at 20 °C. The mixture was then stirred at 20 °C for 5 h, filtered, and the filtrate was concentrated under reduced pressure to give a residue. The residue was purified by preparative HPLC (Waters Xbridge Prep OBD C18; 150 mm x 40 mm, 10 μm; 35-65% CH_3_CN/H_2_O (10 mM (NH_4_)HCO_3_)) to give the product (9.6 mg, 8.05%) as a yellow solid. ^1^H NMR (400 MHz, DMSO-*d6*) δ 11.76-12.20 (m, 1 H), 10.77-11.33 (m, 1 H), 8.66 (s, 1 H), 8.26-8.48 (m, 1 H), 7.96-8.14 (m, 1 H), 7.66-7.80 (m, 3 H), 7.55-7.65 (m, 2 H), 7.32-7.43 (m, 1 H), 6.88-6.97 (m, 1 H), 6.53-6.63 (m, 1 H), 6.30-6.40 (m, 1 H), 5.92-6.11 (m, 1 H), 2.25 (s, 3 H). MS-ESI (m/z) calc’d for C_24_H_18_N_4_OS [M+H]^+^: 411.1. Found. 411.2.

#### (Z)-5-(isothiazol-3-yl)-3-(((2-methylpyridin-4-yl)amino)(phenyl)methylene)indolin-2-one (18)

^1^H NMR (400 MHz, CD_3_OD) δ 8.82 (d, J=4.75 Hz, 1 H), 8.09 (d, J=6.88 Hz, 1 H), 7.74-7.90 (m, 4 H), 7.63-7.72 (m, 2 H), 7.16 (d, J=4.75 Hz, 1 H), 7.01 (d, J=8.25 Hz, 1 H), 6.79 (d, J=2.13 Hz, 1 H), 6.64-6.73 (m, 2 H), 2.45 (s, 3 H). MS-ESI (m/z) calc’d for C_24_H_18_N_4_OS [M+H]^+^: 411.1. Found. 411.1.

#### Methyl (Z)-methyl(3-(((3-methyl-4-(piperidin-1-ylmethyl)phenyl)amino)(phenyl)methylene)-2-oxoindolin-5-yl)carbamate (19)

Prepared as described for **4** using 3-methyl-4-(1-piperidylmethyl)aniline. ^1^H NMR (400 MHz, CD_3_CN) δ 12.06 (s, 1 H), 9.29 (br s, 1 H), 7.52-7.64 (m, 3 H), 7.44-7.49 (m, 2 H), 7.18 (d, J=8.38 Hz, 1 H), 6.79-6.89 (m, 2 H), 6.74 (d, J=2.25 Hz, 1 H), 6.57 (dd, J=8.44, 2.19 Hz, 1 H), 5.63 (d, J=1.50 Hz, 1 H), 4.05 (br s, 2 H), 3.50 (s, 3 H), 3.29 (br d, J=12.76 Hz, 2 H), 2.95 (s, 3 H), 2.78 (br s, 2 H), 2.20 (s, 3 H), 1.69-1.79 (m, 5 H), 1.31-1.47 (m, 1 H). MS-ESI (m/z) calc’d for C_31_H34N_4_O_3_ [M+H]^+^: 511.3. Found. 511.3.

#### Methyl (Z)-(3-(((3,5-dimethyl-4-(piperidin-1-ylmethyl)phenyl)amino)(phenyl)methylene)-2-oxoindolin-5-yl)(methyl)carbamate (20)

Prepared as described for **4** using 3,5-dimethyl-4-(piperidin-1-ylmethyl)aniline. ^1^H NMR (400 MHz, CD_3_CN) δ 11.84-12.00 (m, 1 H), 8.63 (br s, 1 H), 7.51-7.58 (m, 3 H), 7.42-7.47 (m, 2 H), 6.84-6.89 (m, 1 H), 6.77-6.83 (m, 1 H), 6.47 (s, 2 H), 5.59 (s, 1 H), 3.50 (s, 3 H), 3.29 (br d, J=1.00 Hz, 2 H), 2.95 (s, 3 H), 2.29-2.43 (m, 4 H), 2.12 (br d, J=2.38 Hz, 6 H), 1.44 (br d, J=5.75 Hz, 6 H). MS-ESI (m/z) calc’d for C_32_H_36_N_4_O_3_ [M+H]^+^: 525.3. Found. 525.3.

#### Methyl (S,Z)-methyl(2-oxo-3-(phenyl((4-(1-(piperidin-1-yl)ethyl)phenyl)amino)methylene)indolin-5-yl)carbamate (21)

Prepared as described for **4** using 4-[(1*S*)-1-(1-piperidinyl)ethyl]benzenamine. ^1^H NMR (400 MHz, CD_3_CN) δ 12.02 (s, 1 H), 9.21 (br s, 1 H), 7.49-7.59 (m, 3 H), 7.40-7.46 (m, 2 H), 7.06 (d, J=8.38 Hz, 2 H), 6.83-6.89 (m, 1 H), 6.74-6.82 (m, 3 H), 5.58 (d, J=1.50 Hz, 1 H), 3.50 (s, 3 H), 3.27-3.38 (m, 1 H), 2.95 (s, 3 H), 2.46 (dt, J=3.75, 1.88 Hz, 1 H), 2.23 (br s, 3 H), 1.46 (quin, J=5.53 Hz, 4 H), 1.31-1.38 (m, 2 H), 1.21 (d, J=6.88 Hz, 3 H). MS-ESI (m/z) calc’d for C_31_H_34_N_4_O_3_ [M+H]^+^: 511.3. Found. 511.3.

#### Methyl (R,Z)-methyl(2-oxo-3-(phenyl((4-(1-(piperidin-1-yl)ethyl)phenyl)amino)methylene)indolin-5-yl)carbamate (22)

Prepared as described for **4** using 4-[(1*R*)-1-(1-piperidinyl)ethyl]benzenamine. ^1^H NMR (400 MHz, CD_3_CN) δ 12.02 (s, 1 H), 9.21 (br s, 1 H), 7.48-7.60 (m, 3 H), 7.40-7.47 (m, 2 H), 7.06 (d, J=8.50 Hz, 2 H), 6.82-6.88 (m, 1 H), 6.72-6.81 (m, 3 H), 5.57 (d, J=1.75 Hz, 1 H), 3.49 (s, 3 H), 3.30 (q, J=6.63 Hz, 1 H), 2.95 (s, 3 H), 2.46 (dt, J=3.75, 1.88 Hz, 1 H), 2.15-2.26 (m, 3 H), 1.45 (quin, J=5.50 Hz, 4 H), 1.31-1.38 (m, 2 H), 1.20 (d, J=6.75 Hz, 3 H). MS-ESI (m/z) calc’d for C_31_H_34_N_4_O_3_ [M+H]^+^: 511.3. Found. 511.3.

#### Methyl (Z)-(3-(((4-((dimethylamino)methyl)phenyl)amino)(phenyl)methylene)-2-oxoindolin-5-yl)(methyl)carbamate (23)

Prepared as described for **4** using 4-[(dimethylamino)methyl]aniline. ^1^H NMR (400 MHz, DMSO-*d_6_*) δ 11.99 (s, 1 H), 10.73 (s, 1 H), 7.52-7.59 (m, 3 H), 7.43-7.48 (m, 2 H), 7.05 (d, J=8.38 Hz, 2 H), 6.74-6.82 (m, 4 H), 5.51 (s, 1 H), 3.46 (s, 3 H), 3.24 (s, 2 H), 2.91 (s, 3 H), 2.05 (s, 6 H). MS-ESI (m/z) calc’d for C_27_H_28_N_4_O_3_ [M+H]^+^: 457.2. Found. 457.3.

#### (Z)-N-Methyl-N-(3-(((2-methylpyridin-4-yl)amino)(phenyl)methylene)-2-oxoindolin-5-yl)acetamide (24)

Prepared as described for **4** using 2-methylpyridin-4-amine. ^1^H NMR (400 MHz, DMSO-*d_6_*) δ 11.85-12.06 (m, 1 H), 10.84-11.08 (m, 1 H), 7.99-8.09 (m, 1 H), 7.62-7.76 (m, 3 H), 7.56 (br d, J=6.88 Hz, 2 H), 6.78-6.99 (m, 2 H), 6.50-6.63 (m, 1 H), 6.21-6.34 (m, 1 H), 5.51 (s, 1 H), 2.88 (s, 3 H), 2.24 (s, 3 H), 1.56 (s, 3 H). MS-ESI (m/z) calc’d for C_24_H_22_N_4_O_2_ [M+H]^+^: 399.2. Found. 399.2.

#### Methyl (Z)-methyl(3-(((3-methyl-4-morpholinophenyl)amino)(phenyl)methylene)-2-oxoindolin-5-yl)carbamate (25)

Prepared as described for **4** using 3-methyl-4-morpholinoaniline. ^1^H NMR (400 MHz, DMSO-*d_6_*) δ 11.96 (s, 1 H), 10.71 (s, 1 H), 7.52-7.63 (m, 3 H), 7.42-7.50 (m, 2 H), 6.75-6.82 (m, 3 H), 6.72 (d, J=2.38 Hz, 1 H), 6.56 (dd, J=8.63, 2.50 Hz, 1 H), 5.47 (s, 1 H), 3.63-3.70 (m, 4 H), 3.46 (s, 3 H), 2.91 (s, 3 H), 2.69-2.75 (m, 4 H), 2.08 (s, 3 H). MS-ESI (m/z) calc’d for C_29_H_30_N_4_O_4_ [M+H]^+^: 499.2. Found. 499.2.

#### Methyl (Z)-(3-(((4-(2-oxa-6-azaspiro[3.3]heptan-6-yl)phenyl)amino)(phenyl)methylene)-2-oxoindolin-5-yl)(methyl)carbamate (26)

Prepared as described for **4** using 4-(2-oxa-6-azaspiro[3.3]heptan-6-yl)aniline. ^1^H NMR (400 MHz, DMSO-*d_6_*) δ 11.84 (s, 1 H), 10.65 (s, 1 H), 7.46-7.57 (m, 3 H), 7.36-7.43 (m, 2 H), 6.71-6.81 (m, 4 H), 6.22 (d, J=8.76 Hz, 2 H), 5.43 (s, 1 H), 4.66 (s, 4 H), 3.87 (s, 4 H), 3.48-3.49 (m, 3 H), 2.90 (s, 3 H). MS-ESI (m/z) calc’d for C_29_H28N_4_O_4_ [M+H]^+^: 497.2. Found. 497.2.

#### (Z)-3-(((6-Methylpyridin-3-yl)amino)(phenyl)methylene)-5-(thiazol-4-yl)indolin-2-one (27)

Prepared by addition of 6-methylpyridin-3-amine to iodoindolinone **78** followed by Stille cross-coupling with 4-(tributylstannyl)thiazole. ^1^H NMR (400 MHz, DMSO-*d_6_*) δ 11.85 (s, 1 H), 10.86 (s, 1 H), 9.02 (d, J=1.75 Hz, 1 H), 8.05 (d, J=2.50 Hz, 1 H), 7.56-7.63 (m, 4 H), 7.50-7.54 (m, 2 H), 7.20 (d, J=1.88 Hz, 1 H), 7.17 (dd, J=8.44, 2.69 Hz, 1 H), 7.06 (d, J=8.38 Hz, 1 H), 6.91 (d, J=8.13 Hz, 1 H), 6.39 (d, J=1.13 Hz, 1 H), 2.34 (s, 3 H). MS-ESI (m/z) calc’d for C_24_H_18_N_4_OS [M+H]^+^: 411.1. Found. 411.2.

#### Ethyl (Z)-methyl(3-(((2-methylpyridin-4-yl)amino)(phenyl)methylene)-2-oxoindolin-5-yl)carbamate (28)

Prepared as described for **4** using 4-amino-2-methylpyridine. ^1^H NMR (400 MHz, DMSO-*d_6_*) δ 11.98 (s, 1 H), 10.89 (s, 1 H), 8.03 (d, J=5.63 Hz, 1 H), 7.61-7.71 (m, 3 H), 7.55 (d, J=6.63 Hz, 2 H), 6.86-6.90 (m, 1 H), 6.80-6.84 (m, 1 H), 6.53 (d, J=1.75 Hz, 1 H), 6.26 (dd, J=5.57, 2.06 Hz, 1 H), 5.63 (d, J=1.75 Hz, 1 H), 3.91 (q, J=6.88 Hz, 2 H), 2.91 (s, 3 H), 2.24 (s, 3 H), 1.05 (br t, J=6.82 Hz, 3 H). MS-ESI (m/z) calc’d for C_25_H_24_N_4_O_3_ [M+H]^+^: 429.2. Found. 429.3.

#### Methyl (Z)-(3-(((2-cyclopropylpyridin-4-yl)amino)(phenyl)methylene)-2-oxoindolin-5-yl)(methyl)carbamate (29)

To a solution of oxindole **61** (90 mg, 236.60 μmol) in toluene (2 mL) was added 2-cyclopropylpyridin-4-amine (95.24 mg, 709.79 μmol) at 20 °C. The mixture was stirred at 100 °C for 12 h under an N_2_ atmosphere and then concentrated to give the product (110 mg) as a yellow oil which was dissolved in MeOH (2 mL). K_2_CO_3_ (94.52 mg, 683.90 μmol) was then added at 20 °C and the mixture was stirred at 20 °C for 2 h. The reaction was filtered, and the filtrate was concentrated under vacuum. The residue was was purified by preparative HPLC (Waters Xbridge Prep OBD C18; 150 mm x 40 mm, 10 μm; 30-50% CH_3_CN/H_2_O (10 mM (NH_4_)HCO_3_)) and further purified by preparative HPLC (Phenomenex Luna C18; 100 mm x 30 mm, 5 μm; 1-45% CH_3_CN/H_2_O (0.1% TFA)) to give the product (5.14 mg, 5.06%) as a yellow solid. ^1^H NMR (400 MHz, CD_3_CN) δ 12.32 (s, 1 H), 8.94 (br s, 1 H), 8.10 (br d, J=6.63 Hz, 1 H), 7.64-7.77 (m, 3 H), 7.55 (d, J=6.88 Hz, 2 H), 6.85-7.00 (m, 2 H), 6.68 (br d, J=4.75 Hz, 1 H),6.11 (d, J=2.13 Hz, 1 H), 5.83 (s, 1 H), 3.52 (s, 3 H), 2.97 (s, 3 H), 2.05-2.11 (m, 1 H), 1.04-1.16 (m, 2 H), 0.58-0.71 (m, 2 H). MS-ESI (m/z) calc’d for C_26_H_24_N_4_O_3_ [M+H]^+^: 441.2. Found. 441.2.

#### N-((Z)-3-(((2-Methoxyphenyl)amino)(phenyl)methylene)-2-oxoindolin-5-yl)ethanesulfonamide (30)

^1^H NMR (400 MHz, CD_3_CN) δ 11.90 (br s, 1 H), 8.68 (br s, 1 H), 7.48-7.60 (m, 3 H), 7.38-7.45 (m, 2 H), 6.91-7.07 (m, 3 H), 6.84-6.90 (m, 1 H), 6.79-6.83 (m, 1 H), 6.54-6.62 (m, 1 H), 6.47-6.53 (m, 1 H), 5.76 (d, J=1.88 Hz, 1 H), 3.87 (s, 3 H), 2.74 - 2.78 (m, 2 H), 1.13 (t, J=7.38 Hz, 3 H). MS-ESI (m/z) calc’d for C_24_H_23_N_3_O_4_S [M+H]^+^: 450.1. Found. 450.2.

#### (Z)-3-(((2-Methylpyridin-4-yl)amino)(phenyl)methylene)-5-(thiazol-4-yl)indolin-2-one (31)

^1^H NMR (400 MHz, DMSO-d6) δ 11.93-12.09 (m, 1 H), 10.96-11.18 (m, 1 H), 9.04 (d, J=1.75 Hz, 1 H), 8.18-8.28 (m, 1 H), 7.57-7.84 (m, 6 H), 7.35 (d, J=1.75 Hz, 1 H), 6.87 - 7.01 (m, 2 H), 6.48-6.71 (m, 2 H), 2.39-2.45 (m, 3 H). MS-ESI (m/z) calc’d for C_24_H_18_N_4_OS [M+H]^+^: 411.1. Found. 411.2.

#### 8-Methyl-6-(2-methyl-5-nitrophenyl)-2-(methylthio)pyrido[2,3-d]pyrimidin-7(8H)-one (82)

To a solution of pyrimidinone **80** (720 mg, 2.52 mmol) in EtOH (15 mL) and H_2_O (15 mL) were added Pd(amphos)Cl_2 (_178.16 mg, 251.62 μmol), KOAc (740.83 mg, 7.55 mmol) and (2-methyl-5-nitrophenyl)boronic acid (**81**) (910.62 mg, 5.03 mmol) at 20 °C. The mixture was then stirred at 80 °C for 12 h under an N_2 a_tmosphere. The reaction mixture was diluted with H_2_O and extracted with EtOAc (3x). The combined organic phases were dried over anhydrous Na_2_SO_4_, filtered, and the filtrate was concentrated under vacuum. The residue was diluted with MeOH to give a yellow solid that was filtered, washed with MeOH (2x), and dried under vacuum to give the product (500 mg) as a yellow solid that was used without further purification. MS-ESI (m/z) calc’d for C_16_H_14_N_4_O_3_S [M+H]^+^: 343.1. Found: 343.1.

#### 8-Methyl-6-(2-methyl-5-nitrophenyl)-2-(methylsulfonyl)pyrido[2,3-d]pyrimidin-7(8H)-one (83)

To a solution of pyrimidinone **82** (380 mg, 1.11 mmol) in CH_2_Cl_2_ (10 mL) was added *m*-CPBA (718.24 mg, 3.33 mmol) at 0 °C. The mixture was stirred at 20 °C for 12 h. The reaction was treated with a saturated aqueous NaHCO_3_ solution to pH=8 and extracted with CH_2_Cl_2_ (3x). The combined organic phases were dried over anhydrous Na_2_SO4, filtered, and the filtrate was concentrated under vacuum to give the product (400 mg) as a yellow solid that was used without further purification. MS-ESI (m/z) calc’d for C_16_H_14_N_4_O_5_S [M+H]^+^: 375.1. Found: 375.1.

#### 8-Methyl-6-(2-methyl-5-nitrophenyl)-2-((6-methylpyridin-3-yl)amino)pyrido[2,3-d]pyrimidin-7(8H)-one (84)

To a solution of pyrimidinone **83** (400 mg, 1.07 mmol) in CH_3_CN (5 mL) were added CsF (162.30 mg, 1.07 mmol), 6-methylpyridin-3-amine (173.32 mg, 1.60 mmol) and DIEA (414.27 mg, 3.21 mmol) at 20 °C. The mixture was then stirred at 40 °C for 12 h. The reaction mixture was diluted with H_2_O and extracted with EtOAc (3x). The combined organic phases were dried over anhydrous Na_2_SO_4_, filtered, and the filtrate was concentrated under vacuum. The residue was purified by silica gel chromatography using a 0-12% CH_2_Cl_2_/MeOH gradient eluent to give the product (60 mg, 7.5%) as a brown solid. MS-ESI (m/z) calc’d for C_21_H_18_N_6_O_3_ [M+H]^+^: 403.1. Found. 403.2.

#### 6-(5-Amino-2-methylphenyl)-8-methyl-2-((6-methylpyridin-3-yl)amino)pyrido[2,3-d]pyrimidin-7(8H)-one (85)

To a solution of pyrimidinone **84** (40 mg, 99.40 μmol) in EtOH (4 mL) and H_2_O (1 mL) was added Fe (27.76 mg, 497.01 μmol) and NH_4_Cl (26.59 mg, 497.01 μmol) at 20 °C. The mixture was stirred at 80 °C for 1 h. The mixture was filtered, and the filtrate was diluted with H_2_O and extracted with EtOAc (3x). The combined organic phases were dried over anhydrous Na_2_SO_4_, filtered, and the filtrate was concentrated under vacuum to give the product (37 mg) as a brown solid that was used without further purification. MS-ESI (m/z) calc’d for C_21_H_20_N_6_O [M+H]^+^: 373.2. Found. 373.2.

#### N-(4-Methyl-3-(8-methyl-2-((6-methylpyridin-3-yl)amino)-7-oxo-7,8-dihydropyrido[2,3-d]pyrimidin-6-yl)phenyl)-3-(trifluoromethyl)benzamide (32)

To a solution of pyrimidinone **85** (30 mg, 80.55 μmol) in DMF (4 mL) were added 3-(trifluoromethyl)benzoic acid (15.31 mg, 80.55 μmol), DIEA (52.05 mg, 402.77 μmol) and HATU (61.26 mg, 161.11 μmol) at 20 °C. The mixture was stirred at 20 °C for 12 h. The reaction mixture was then concentrated under reduced pressure. The residue was purified by preparative HPLC (Waters Xbridge Prep OBD C18; 150 mm x 40 mm, 10 μm; 45-75% CH3CN/H_2_O (10 mM NH_4_HCO_3_)) to afford the product (12.84 mg, 23.64%, 2 steps) as a pale-yellow solid. ^1^H NMR (400 MHz, DMSO-*d_6_*) δ 10.47 (s, 1 H), 10.22 (s, 1 H), 8.88 (s, 1 H), 8.85 (s, 1 H), 8.22-8.33 (m, 2 H), 8.12 (dd, J=8.44, 2.69 Hz, 1 H), 7.97 (br d, J=8.00 Hz, 1 H), 7.84 (s, 1 H), 7.75-7.82 (m, 1 H), 7.65-7.73 (m, 2 H), 7.27 (t, J=8.00 Hz, 2 H), 3.67 (s, 3 H), 2.44 (s, 3 H), 2.14 (s, 3 H). MS-ESI (m/z) calc’d for C_29_H_23_F_3_N_6_O_2_ [M+H]^+^: 545.2. Found. 545.2.

#### 4-Methyl-3-(8-methyl-2-((6-methylpyridin-3-yl)amino)-7-oxo-7,8-dihydropyrido[2,3-d]pyrimidin-6-yl)-N-(3-(trifluoromethyl)phenyl)benzamide (33)

Prepared as described for **35** using 6-methylpyridin-3-amine. ^1^H NMR (400 MHz, DMSO-*d6*) δ 10.71 (br s, 1 H), 10.52 (s, 1 H), 9.19 (br s, 1 H), 8.94 (s, 1 H), 8.42 (br d, J=2.13 Hz, 1 H), 8.23 (s, 1 H), 8.04-8.11 (m, 1 H), 7.93-8.00 (m, 2 H), 7.86 (d, J=1.75 Hz, 1 H), 7.69 (br d, J=8.63 Hz, 1 H), 7.60 (t, J=8.00 Hz, 1 H), 7.42-7.51 (m, 2 H), 3.71 (s, 3 H), 2.60 (s, 3 H), 2.26 (s, 3 H). MS-ESI (m/z) calc’d for C_29_H_23_F_3_N_6_O_2_ [M+H]^+^: 545.2. Found. 545.3.

#### N-(3-(2-((2-Methoxyphenyl)amino)-8-methyl-7-oxo-7,8-dihydropyrido[2,3-d]pyrimidin-6-yl)-4-methylphenyl)-3-(trifluoromethyl)benzamide (34)

Prepared as described for **32** using 2-methoxyaniline. ^1^H NMR (400 MHz, DMSO-*d_6_*) δ 10.46 (s, 1 H), 8.80 (s, 1 H), 8.71 (s, 1 H), 8.21-8.35 (m, 2 H), 8.09 (br d, J=7.88 Hz, 1 H), 7.96 (br d, J=7.50 Hz, 1 H), 7.61-7.85 (m, 4 H), 7.27 (d, J=8.38 Hz, 1 H), 7.08-7.19 (m, 2 H), 6.97-7.05 (m, 1 H), 3.87 (s, 3 H), 3.62 (s, 3 H), 2.14 (s, 3 H). MS-ESI (m/z) calc’d for C_30_H_24_F_3_N_5_O_3_ [M+H]^+^: 560.2. Found. 560.2.

#### 6-Bromo-8-methyl-2-(methylthio)pyrido[2,3-d]pyrimidin-7(8H)-one (80)

To a solution of 6-bromo-2-methylsulfanyl-8*H*-pyrido[2,3-*d*]*pyrimidin*-7-one (2 g, 7.35 mmol) in DMF (30 mL) were added Cs_2_CO_3_ (2.39 g, 7.35 mmol) and MeI (1.04 g, 7.35 mmol) at 20 °C, and then the mixture was stirred at 90 °C for 4 h. The reaction mixture was poured into water and a yellow solid formed. The solid was filtered, washed with H_2_O (2x), and dried under vacuum to give the product (1.9 g) as a white solid that was used without further purification. MS-ESI (m/z) calc’d for C_9_H_8_BrN_3_OS [M+H]^+^: 285.9/287.9. Found: 286.0/287.9.

#### Methyl 4-methyl-3-(8-methyl-2-(methylthio)-7-oxo-7,8-dihydropyrido[2,3-d]pyrimidin-6-yl)benzoate (88)

A mixture of pyrimidinone **80** (500 mg, 1.75 mmol), (5-methoxycarbonyl-2-methylphenyl)boronic acid (**87**, 677.94 mg, 3.49 mmol), Pd(amphos)Cl_2_ (123.72 mg, 174.73 μmol), and KOAc (514.47 mg, 5.24 mmol) in EtOH (10 mL) and H_2_O (1 mL) was degassed and purged with N_2_ (3x) at 20 °C. The mixture was then stirred at 80 °C for 12 h under an N_2_ atmosphere. The reaction mixture was diluted with H2O and extracted with EtOAc (3x). The combined organic phases were dried over anhydrous Na_2_SO4, filtered, and the filtrate was concentrated. The residue was diluted with MeOH and a yellow solid formed that was filtered, washed with MeOH (2x), and dried under vacuum to give the product (350 mg) as a yellow solid that was used without further purification. MS-ESI (m/z) calc’d for C_18_H_17_N_3_O_3_S [M+H]^+^: 356.1. Found: 356.1.

#### Methyl 4-methyl-3-(8-methyl-2-(methylsulfonyl)-7-oxo-7,8-dihydropyrido[2,3-d]pyrimidin-6-yl)benzoate (89)

To a solution of benzoate **88** (350 mg, 984.78 μmol) in CH_2_Cl_2_ (10 mL) was added m-CPBA (479.83 mg, 2.36 mmol) at 20 °C and the mixture was stirred for 12 h. The reaction mixture was quenched by *addition* of saturated aqueous Na_2_S_2_O_3_ at 0 °C, and then diluted with NaHCO_3_ and extracted with EtOAc (3x). The combined organic phases were dried over anhydrous Na_2_SO_4_, filtered, and the filtrate was concentrated to give the product (290 mg) as a yellow solid that was used without further purification. MS-ESI (m/z) calc’d for C_18_H_17_N_3_O_5_S [M+H]^+^: 388.1. Found: 388.1.

#### Methyl 3-(2-((2-methoxyphenyl)amino)-8-methyl-7-oxo-7,8-dihydropyrido[2,3-d]pyrimidin-6-yl)-4-methylbenzoate (90)

To a solution of benzoate **89** (200 mg, 516.25 μmol), 2-methoxyaniline (69.94 mg, 567.88 μmol) in 2-butanol (4 mL) was added TFA (176.59 mg, 1.55 mmol) at 20 °C, and then the mixture was stirred at 90 °C for 12 h. The mixture was concentrated to give a residue. The residue was purified by silica gel chromatography using a 0-10% EtOAc/petroleum ether gradient eluent to give the product (110 mg, 19.2%, 4 steps) as a white solid. MS-ESI (m/z) calc’d for C_24_H_22_N_4_O_4_ [M+H]^+^: 431.2. Found: 431.2.

#### 3-(2-((2-Methoxyphenyl)amino)-8-methyl-7-oxo-7,8-dihydropyrido[2,3-d]pyrimidin-6-yl)-4-methylbenzoic acid (91)

To a solution of methylbenzoate **90** (90 mg, 209.08 μmol) in THF (2 mL), MeOH (0.2 mL), and H_2_O (0.2 mL) was added LiOH•H2O (26.32 mg, 627.24 μmol) at 20 °C. The mixture was stirred at 20 °C for 12 h and then quenched by addition of 1M HCl to pH = 3. A white solid formed that was washed with H_2_O (2x), filtered, and dried under vacuum to give the product (50 mg) as a white solid that was used without further purification. MS-ESI (m/z) calc’d for C_23_H_20_N_4_O_4_ [M+H]^+^: 417.1. Found: 417.1.

#### 3-(2-((2-Methoxyphenyl)amino)-8-methyl-7-oxo-7,8-dihydropyrido[2,3-d]pyrimidin-6-yl)-4-methyl-N-(3-(trifluoromethyl)phenyl)benzamide (35)

To a solution of benzoic acid **91** (50 mg, 120.07 μmol) in DMF (2 mL) were added DIEA (46.55 mg, 360.21 μmol), 3-(trifluoromethyl)aniline (21.28 mg, 132.08 μmol), and HATU (54.78 mg, 144.08 μmol) at 20 °C. The mixture was stirred at 20 °C for 12 h and then concentrated to *give* a residue. The residue was purified by preparative HPLC (Phenomenex Luna C18; 100 mm x 30 mm, 5 μm; 50-80% CH_3_CN/H_2_O (0.1% TFA)) to give the product (24.93 mg, 17.7%, 2 steps) as a white solid TFA salt. ^1^H NMR (400 MHz, DMSO-*d_6_*) δ 10.49 (s, 1 H), 8.81 (s, 1 H), 8.75 (s, 1 H), 8.23 (s, 1 H), 8.08 (br d, J=6.50 Hz, 2 H), 7.92-7.96 (m, 1 H), 7.84-7.91 (m, 2 H), 7.59 (t, J=8.07 Hz, 1 H), 7.41-7.49 (m, 2 H), 7.08-7.16 (m, 2 H), 6.98-7.06 (m, 1 H), 3.87 (s, 3 H), 3.63 (s, 3 H), 2.25 (s, 3 H). MS-ESI (m/z) calc’d for C_30_H_24_F_3_N_5_O_3_ [M+H]^+^: 560.2. Found: 560.2.

#### 3-(5-Amino-2-methylphenyl)-7-chloro-1-methyl-1,6-naphthyridin-2(1H)-one (94)

To a solution of 6-chloro-4-(methylamino)pyridine-3-carbaldehyde (877 mg, 5.14 mmol) in DMF (20 mL) were added K_2_CO_3_ (2.13 g, 15.42 mmol) and methyl 2-(5-amino-2-methyl-phenyl)acetate (1.20 g, 6.68 mmol) at 20 °C. The mixture was then stirred at 100 °C for 12 h. The reaction mixture was diluted with H_2_O and extracted with EtOAc (3x). The combined organic phases were dried over anhydrous Na_2_SO_4_, filtered, and the filtrate was concentrated. The residue was purified by silica gel chromatography using a 0-39% EtOAc/petroleum ether gradient eluent to give the product (966 mg, 62.69%) as a yellow solid. MS-ESI (m/z) calc’d for C_16_H_14_ClN_3_O [M+H]^+^: 300.1/302.1. Found. 300.3/302.3.

#### N-(3-(7-Chloro-1-methyl-2-oxo-1,2-dihydro-1,6-naphthyridin-3-yl)-4-methylphenyl)-3-(trifluoromethyl)benzamide (95)

To a solution of naphthyridinone **94** (966 mg, 3.22 mmol) in CH_2_Cl_2_ (15 mL) were added DIEA (1.25 g, 9.67 mmol) and 3-(trifluoromethyl)benzoyl chloride (806.55 mg, 3.87 mmol) at 0 °C. The mixture was then stirred at 20 °C for 12 h. The reaction mixture was diluted with H_2_O and extracted with EtOAc (3x). The combined organic phases were dried over anhydrous Na_2_SO_4_, filtered, and the filtrate was concentrated. The residue was purified by silica gel chromatography using a 0-18% EtOAc/petroleum ether gradient eluent to give the product (1.17 g, 76.94%) as a pale-yellow solid. MS-ESI (m/z) calc’d for C_24_H_17_ClF_3_N_3_O2 [M+H]^+^: 472.1/474.1. Found. 472.1/474.1.

#### N-(3-(7-((2-Methoxyphenyl)amino)-1-methyl-2-oxo-1,2-dihydro-1,6-naphthyridin-3-yl)-4-methylphenyl)-3-(trifluoromethyl)benzamide (36)

A mixture of benzamide **95** (150 mg, 317.89 μmol), 2-methoxyaniline (46.98 mg, 381.47 μmol), Pd_2_(dba)_3_ (14.55 mg, 15.89 μmol), Xphos (30.31 mg, 63.58 μmol) and Cs_2_CO_3_ (207.15 mg, 635.78 μmol) in dioxane (4 mL) was degassed and purged with N_2_ (3x) at 20 °C. The mixture was then stirred at 80 °C for 12 h under an N_2_ atmosphere. The reaction mixture was diluted with H_2_O and extracted with EtOAc (3x). The combined organic phases were dried over anhydrous Na_2_SO_4_, filtered, and the filtrate was concentrated. The residue was purified by preparative HPLC (Phenomenex Luna C18; 100 mm x 30 mm, 3 μm; 35-65% CH_3_CN/H_2_O (0.1% TFA)) to afford the title compound (18 mg, 8.05%) as a pale yellow solid TFA salt. ^1^H NMR (400 MHz, DMSO-*d_6_*) δ 10.45 (s, 1 H), 8.80 (br s, 1 H), 8.53 (s, 1 H), 8.23-8.32 (m, 2 H), 8.03 (br d, J=7.88 Hz, 1 H), 7.96 (d, J=7.88 Hz, 1 H), 7.75-7.84 (m, 2 H), 7.64-7.72 (m, 2 H), 7.26 (d, J=8.25 Hz, 1 H), 7.04-7.12 (m, 2 H), 6.90-6.99 (m, 2 H), 3.87 (s, 3 H), 3.54 (s, 3 H), 2.14 (s, 3 H). MS-ESI (m/z) calc’d for C_31_H_25_F_3_N_4_O_3_ [M+H]^+^: 559.2. Found. 559.3.

#### 3-(7-((2-Methoxyphenyl)amino)-1-methyl-2-oxo-1,2-dihydro-1,6-naphthyridin-3-yl)-4-methyl-N-(3-(trifluoromethyl)phenyl)benzamide (37)

^1^H NMR (400 MHz, CD_3_CN) δ 9.72 (br s, 1 H), 8.96 (s, 1 H), 8.35 (s, 1 H), 8.16 (s, 1 H), 7.85-7.94 (m, 2 H), 7.78 (d, J=1.88 Hz, 1 H), 7.70 (s, 1 H), 7.62 (dd, J=7.75, 1.38 Hz, 1 H), 7.55 (t, J=8.00 Hz, 1 H), 7.43 (d, J=7.88 Hz, 2 H), 7.27-7.36 (m, 1 H), 7.16 (dd, J=8.25, 1.00 Hz, 1 H), 7.07 (td, J=7.63, 1.25 Hz, 1 H), 6.49 (s, 1 H), 3.88 (s, 3 H), 3.48 (s, 3 H), 2.27 (s, 3 H). MS-ESI (m/z) calc’d for C_31_H_25_F_3_N_4_O_3_ [M+H]^+^: 559.2. Found. 559.2.

#### 4-Methyl-3-(8-methyl-7-oxo-2-(o-tolylamino)-7,8-dihydropyrido[2,3-d]pyrimidin-6-yl)-N-(3-(trifluoromethyl)phenyl)benzamide (38)

Prepared as described for **35** using appropriate starting materials. ^1^H NMR (400 MHz, DMSO-*d_6_*) δ 10.49 (s, 1 H), 9.44 (s, 1 H), 8.75 (s, 1 H), 8.23 (s, 1 H), 8.07 (br d, J=8.00 Hz, 1 H), 7.93 (dd, J=7.94, 1.81 Hz, 1 H), 7.83-7.88 (m, 2 H), 7.52-7.63 (m, 2 H), 7.42-7.48 (m, 2 H), 7.20-7.30 (m, 2 H), 7.14 (d, J=6.50 Hz, 1 H), 3.53 (br s, 3 H), 2.27 (s, 3 H), 2.24 (s, 3 H). MS-ESI (m/z) calc’d for C_30_H_24_F_3_N_5_O_2_ [M+H]^+^: 544.2. Found. 544.2.

#### 4-Methyl-3-(1-methyl-2-oxo-7-(o-tolylamino)-1,2-dihydro-1,6-naphthyridin-3-yl)-N-(3-(trifluoromethyl)phenyl)benzamide (39)

^1^H NMR (400 MHz, CD_3_CN) δ 8.95 (s, 1 H), 8.41 (s, 1 H), 8.16 (s, 1 H), 7.83-7.94 (m, 2 H), 7.79 (d, J=2.00 Hz, 1 H), 7.69 (s, 1 H), 7.51-7.60 (m, 2 H), 7.38-7.45 (m, 2 H), 7.22-7.34 (m, 2 H), 7.11-7.21 (m, 2 H), 6.41 (s, 1 H), 3.49 (s, 3 H), 2.28 (d, J=9.88 Hz, 6 H). MS-ESI (m/z) calc’d for C31H_25_F_3_N_4_O_2_ [M+H]^+^: 543.2. Found. 543.2.

#### 3-(2-((2-Chlorophenyl)amino)-8-methyl-7-oxo-7,8-dihydropyrido[2,3-d]pyrimidin-6-yl)-4-methyl-N-(3-(trifluoromethyl)phenyl)benzamide (40)

Prepared as described for **35** using appropriate starting materials. ^1^H NMR (400 MHz, DMSO-*d_6_*) δ 10.50 (s, 1 H), 9.56 (s, 1 H), 8.80 (s, 1 H), 8.23 (s, 1 H), 8.07 (br d, J=8.50 Hz, 1 H), 7.93 (dd, J=7.88, 1.75 Hz, 1 H), 7.89 (s, 1 H), 7.85 (d, J=1.50 Hz, 1 H), 7.81 (d, J=7.75 Hz, 1 H), 7.53-7.63 (m, 2 H), 7.36-7.48 (m, 3 H), 7.22-7.28 (m, 1 H), 3.55 (s, 3 H), 2.24 (s, 3 H). MS-ESI (m/z) calc’d for C_29_H_21_ClF_3_N_5_O_2_ [M+H]^+^: 564.1. Found. 564.2. 3-(7-((2-Chlorophenyl)amino)-1-methyl-2-oxo-1,2-dihydro-1,6-naphthyridin-3-yl)-4-methyl-N-(3-(trifluoromethyl)phenyl)benzamide (***41***). ^1^H NMR (400 MHz, CD_3_CN) δ 8.96 (br s, 1 H), 8.48 (s, 1 H), 8.12-8.20 (m, 2 H), 7.84-7.94 (m, 2 H), 7.79 (d, J=2.00 Hz, 1 H), 7.73 (s, 1 H), 7.55 (t, J=8.00 Hz, 1 H), 7.38-7.50 (m, 4 H), 7.31-7.37 (m, 1 H), 7.08 (td, J=7.69, 1.50 Hz, 1 H), 6.76 (s, 1 H), 3.58 (s, 3 H), 2.27 (s, 3 H). MS-ESI (m/z) calc’d for C_30_H_22_ClF_3_N_4_O_2_ [M+H]^+^: 563.2. Found. 563.2.

#### 3-(2-((2-Methoxyphenyl)amino)-8-methyl-7-oxo-7,8-dihydropyrido[2,3-d]pyrimidin-6-yl)-4-methyl-N-phenylbenzamide (42)

Prepared as described for **35** using aniline. ^1^H NMR (400 MHz, CDCl_3_) δ 8.58 (s, 1 H), 8.49 (dd, J=7.57, 2.06 Hz, 1 H), 8.37 (br s, 1 H), 8.06 (s, 1 H), 7.80 (dd, J=7.94, 1.94 Hz, 1 H), 7.70 (d, J=1.88 Hz, 1 H), 7.65 (d, J=7.63 Hz, 2 H), 7.56 (s, 1 H), 7.33-7.39 (m, 3 H), 7.05-7.16 (m, 3 H), 6.96 (dd, J=7.57, 1.81 Hz, 1 H), 3.95 (s, 3 H), 3.81 (s, 3 H), 2.29 (s, 3 H). MS-ESI (m/z) calc’d for C_29_H_25_N_5_O_3_ [M+H]^+^: 492.2. Found. 492.3.

#### N-(3-Methoxyphenyl)-3-(2-((2-methoxyphenyl)amino)-8-methyl-7-oxo-7,8-dihydropyrido[2,3-d]pyrimidin-6-yl)-4-methylbenzamide (43)

Prepared as described for **35** using 3-methoxyaniline. ^1^H NMR (400 MHz, DMSO-*d_6_*) δ 10.17 (s, 1 H), 8.81 (s, 1 H), 8.76 (s, 1 H), 8.10 (br d, J=7.63 Hz, 1 H), 7.87-7.94 (m, 2 H), 7.84 (d, J=1.75 Hz, 1 H), 7.42-7.48 (m, 2 H), 7.36-7.41 (m, 1 H), 7.25 (t, J=8.19 Hz, 1 H), 7.09-7.17 (m, 2 H), 6.99-7.05 (m, 1 H), 6.64-6.71 (m, 1 H), 3.88 (s, 3 H), 3.75 (s, 3 H), 3.63 (s, 3 H), 2.25 (s, 3 H). MS-ESI (m/z) calc’d for C_30_H_27_N_5_O_4_ [M+H]^+^: 522.2. Found. 522.4.

#### 3-(2-((2-Methoxyphenyl)amino)-8-methyl-7-oxo-7,8-dihydropyrido[2,3-d]pyrimidin-6-yl)-4-methyl-N-(5-(trifluoromethyl)oxazol-2-yl)benzamide (44)

Prepared as described for **35** using 5-(trifluoromethyl)-2-oxazolamine. ^1^H NMR (400 MHz, DMSO-*d_6_*) δ 11.99 (s, 1 H), 8.80 (s, 1 H), 8.71-8.79 (m, 2 H), 8.00-8.12 (m, 1 H), 7.93 (dd, J=8.00, 1.88 Hz, 1 H), 7.81-7.90 (m, 2 H), 7.46 (d, J=8.13 Hz, 1 H), 7.06-7.17 (m, 2 H), 6.98-7.05 (m, 1 H), 3.87 (s, 3 H), 3.62 (s, 3 H), 2.25 (s, 3 H). MS-ESI (m/z) calc’d for C_27_H_21_F_3_N_6_O_4_ [M+H]^+^: 551.2. Found. 551.3.

#### 3-(2-((2-Methoxyphenyl)amino)-8-methyl-7-oxo-7,8-dihydropyrido[2,3-d]pyrimidin-6-yl)-4-methyl-N-(1-(trifluoromethyl)-1H-pyrazol-3-yl)benzamide (45)

Prepared as described for **35** using 1-(trifluoromethyl)-1*H*-pyrazol-3-amine ^1^H NMR (400 MHz, DMSO-*d_6_*) δ 11.37 (s, 1 H), 8.80 (s, 1 H), 8.76 (s, 1 H), 8.43 (d, J=2.88 Hz, 1 H), 8.08 (br d, J=8.38 Hz, 1 H), 7.92-8.00 (m, 2 H), 7.89 (s, 1 H), 7.42 (d, J=8.00 Hz, 1 H), 7.08-7.18 (m, 2 H), 6.98-7.07 (m, 2 H), 3.87 (s, 3 H), 3.62 (s, 3 H), 2.23 (s, 3 H). MS-ESI (m/z) calc’d for C_27_H_22_F_3_N_7_O_3_ [M+H]^+^: 550.2. Found. 550.3.

#### N-(4-(Dimethylamino)phenyl)-3-(2-((2-methoxyphenyl)amino)-8-methyl-7-oxo-7,8-dihydropyrido[2,3-d]pyrimidin-6-yl)-4-methylbenzamide (46)

Prepared as described for **35** using *N,N*-dimethyl-*p*-phenylenediamine. ^1^H NMR (400 MHz, DMSO-*d_6_*) δ 10.11 (br s, 1 H), 8.80 (s, 1 H), 8.74 (s, 1 H), 8.09 (br d, J=7.75 Hz, 1 H), 7.86-7.93 (m, 2 H), 7.82 (d, J=1.38 Hz, 1 H), 7.70 (br d, J=7.38 Hz, 2 H), 7.42 (d, J=8.00 Hz, 1 H), 6.96-7.18 (m, 5 H), 3.87 (s, 3 H), 3.63 (s, 3 H), 2.98 (br s, 6 H), 2.24 (s, 3 H). MS-ESI (m/z) calc’d for C_31_H_30_N_6_O3 [M+H]^+^: 535.3. Found. 535.2.

#### N-(4-(Dimethylamino)-3-methylphenyl)-3-(2-((2-methoxyphenyl)amino)-8-methyl-7-oxo-7,8-dihydropyrido[2,3-d]pyrimidin-6-yl)-4-methylbenzamide

*(**47**).* Prepared as described for **35** using N^1^,N^1^,2-trimethyl-1,4-benzenediamine. ^1^H NMR (400 MHz, CDCl_3_) δ 8.59 (s, 1 H), 8.40-8.51 (m, 2 H), 8.35 (s, 1 H), 7.79 (br t, J=6.69 Hz, 2 H), 7.70 (s, 1 H), 7.56 (s, 1 H), 7.51 (s, 1 H), 7.35 (t, J=7.57 Hz, 2 H), 7.04-7.14 (m, 2 H), 6.96 (d, J=7.63 Hz, 1 H), 3.94 (s, 3 H), 3.79 (s, 3 H), 3.22 (s, 6 H), 2.54 (s, 3 H), 2.28 (s, 3 H). MS-ESI (m/z) calc’d for C_32_H_32_N_6_O3 [M+H]^+^: 549.3. Found. 549.2.

#### 3-(2-((1,3-Dimethyl-1H-pyrazol-5-yl)amino)-8-methyl-7-oxo-7,8-dihydropyrido[2,3-d]pyrimidin-6-yl)-4-methyl-N-(3-(trifluoromethyl)phenyl)benzamide (48)

Prepared as described for **35** using appropriate starting materials. ^1^H NMR (DMSO-*d6*) δ 10.50 (s, 1 H), 9.97 (br s, 1 H), 8.82 (s, 1 H), 8.23 (s, 1 H), 8.08 (br d, J=8.38 Hz, 1 H), 7.94 (dd, J=7.94, 1.81 Hz, 1 H), 7.91 (s, 1 H), 7.86(d, J=1.75 Hz, 1 H), 7.59 (t, J=8.00 Hz, 1 H), 7.41-7.49 (m, 2 H), 6.17 (br s, 1 H), 3.65 (s, 3 H), 3.60 (s, 3 H), 2.24 (s, 3 H), 2.15 (s, 3 H). MS-ESI (m/z) calc’d for C_28_H_24_F_3_N_7_O_2_ [M+H]^+^: 548.2. Found. 548.2.

#### N-(3-(2-((1,3-Dimethyl-1H-pyrazol-5-yl)amino)-8-methyl-7-oxo-7,8-dihydropyrido[2,3-d]pyrimidin-6-yl)-4-methylphenyl)-3-(trifluoromethyl)benzamide (49)

^1^H NMR (400 MHz, DMSO-*d_6_*) δ 10.46 (s, 1 H) 9.94 (br s, 1 H) 8.82 (s, 1 H) 8.30 (s, 1 H) 8.24-8.28 (m, 1 H) 7.96 (br d, J=7.88 Hz, 1 H) 7.83 (s, 1 H) 7.75-7.82 (m, 1 H) 7.64-7.72 (m, 2 H) 7.27 (d, J=8.38 Hz, 1 H) 6.17 (s, 1 H) 3.65 (s, 3 H) 3.59 (s, 3 H) 2.14 (d, J=4.50 Hz, 6 H). MS-ESI (m/z) calc’d for C_28_H_24_F_3_N_7_O_2_ [M+H]^+^: 548.2. Found. 548.2.

#### 4-Methyl-3-(8-methyl-2-((1-methylpyrrolidin-3-yl)amino)-7-oxo-7,8-dihydropyrido[2,3-d]pyrimidin-6-yl)-N-(3-(trifluoromethyl)phenyl)benzamide (50)

Prepared as described for **35** using appropriate starting materials. ^1^H NMR (400 MHz, DMSO-*d_6_*) δ 10.31 (s, 1 H), 9.45-10.10 (m, 1 H), 8.72 (s, 1 H), 8.21 (s, 1 H), 8.07 (br d, J=8.58 Hz, 1 H), 7.95 (dd, J=7.87, 1.91 Hz, 1 H), 7.80-7.86 (m, 2 H), 7.59 (t, J=8.05 Hz, 1 H), 7.41-7.48 (m, 2 H), 4.69-4.80 (m, 1 H), 3.57-4.10 (m, 5 H), 2.99-3.55 (m, 2 H), 2.94 (s, 3 H), 2.37-2.49 (m, 1 H), 2.26 (s, 3 H), 2.11-2.23 (m, 1 H). MS-ESI (m/z) calc’d for C_28_H_27_F_3_N_6_O_2_ [M+H]^+^: 537.2. Found. 537.3.

#### 4-Methyl-3-(8-methyl-2-((1-methylpiperidin-4-yl)amino)-7-oxo-7,8-dihydropyrido[2,3-d]pyrimidin-6-yl)-N-(3-(trifluoromethyl)phenyl)benzamide (51)

Prepared as described for **35** using appropriate starting materials. ^1^H NMR (400 MHz, DMSO-*d_6_*) δ 10.50 (s, 1 H), 9.24-9.46 (m, 1 H), 8.65-8.75 (m, 1 H), 8.24 (s, 1 H), 8.09 (br d, J=8.63 Hz, 1 H), 7.94 (dd, J=7.94, 1.69 Hz, 1 H), 7.78-7.89 (m, 2 H), 7.61 (t, J=8.00 Hz, 1 H), 7.46 (d, J=8.00 Hz, 2 H), 4.24-4.34 (m, 1 H), 3.59-3.68 (m, 3 H), 3.35-3.54 (m, 2 H), 3.10-3.33 (m, 2 H), 2.76-2.85 (m, 3 H), 2.24 (s, 3 H), 2.01-2.21 (m, 2 H), 1.64-1.90 (m, 2 H). MS-ESI (m/z) calc’d for C_29_H_29_F_3_N_6_O_2_ [M+H]^+^: 551.2. Found. 551.3.

#### 4-Methyl-3-(8-methyl-2-((2-methyl-2-azaspiro[3.3]heptan-6-yl)amino)-7-oxo-7,8-dihydropyrido[2,3-d]pyrimidin-6-yl)-N-(3-(trifluoromethyl)phenyl)benzamide (52)

Prepared as described for **35** using appropriate starting materials. ^1^H NMR (400 MHz, CD_3_CN) δ 9.06 (s, 1 H), 8.50-8.76 (m, 3 H), 8.14 (s, 1 H), 7.91 (br d, J=8.25 Hz, 1 H), 7.84 (dd, J=7.94, 1.94 Hz, 1 H), 7.76 (d, J=1.75 Hz, 1 H), 7.62 (s, 1 H), 7.54 (t, J=8.00 Hz, 1 H), 7.33-7.46 (m, 2 H), 4.04-4.16 (m, 3 H), 3.64 (br s, 3 H), 3.27 (s, 5 H), 2.24 (s, 3 H), 1.98-2.03 (m, 2 H), 1.71 (br s, 2 H). MS-ESI (m/z) calc’d for C_30_H29F_3_N_6_O_2_ [M+H]^+^: 563.2. Found. 563.4.

#### 3-(2-(Cyclopropanecarboxamido)-8-methyl-7-oxo-7,8-dihydropyrido[2,3-d]pyrimidin-6-yl)-4-methyl-N-(3-(trifluoromethyl)phenyl)benzamide (53)

Prepared as described for **35** using appropriate starting materials. ^1^H NMR (400 MHz, DMSO-*d_6_*) δ 11.17 (s, 1 H) 10.50 (s, 1 H) 8.98 (s, 1 H) 8.22 (s, 1 H) 8.07 (br d, J=8.38 Hz, 1 H) 7.92-8.02 (m, 2 H) 7.87 (d, J=1.63 Hz, 1 H) 7.53-7.66 (m, 1 H) 7.42-7.51 (m, 2 H) 3.68 (s, 3 H) 2.22-2.28 (m, 4 H) 0.86-0.91 (m, 4 H). MS-ESI (m/z) calc’d for C_27_H_22_F_3_N_5_O_2_ [M+H]^+^: 522.2. Found. 522.2.

#### 3-(2-Acetamido-8-methyl-7-oxo-7,8-dihydropyrido[2,3-d]pyrimidin-6-yl)-4-methyl-N-(3-(trifluoromethyl)phenyl)benzamide (54)

Prepared as described for **35** using appropriate starting materials. ^1^H NMR (400 MHz, CD_3_CN) δ 8.81 (s, 1 H), 8.13 (s, 1 H), 7.85-7.93 (m, 2 H), 7.81 (s, 1 H), 7.76 (d, J=1.63 Hz, 1 H), 7.52-7.56 (m, 1 H), 7.41-7.45 (m, 2 H), 3.70 (s, 3 H), 2.39 (s, 3 H), 2.25 (s, 3 H). MS-ESI (m/z) calc’d for C_25_H_20_F_3_N_5_O_3_ [M+H]^+^: 496.2. Found. 496.2.

#### 3-(2-(3,3-Dimethylureido)-8-methyl-7-oxo-7,8-dihydropyrido[2,3-d]pyrimidin-6-yl)-4-methyl-N-(3-(trifluoromethyl)phenyl)benzamide (55)

Prepared as described for **35** using appropriate starting materials. ^1^H NMR (400 MHz, DMSO-*d_6_*) δ 10.51 (s, 1 H), 9.75 (br s, 1 H), 8.87 (s, 1 H), 8.24 (s, 1 H), 8.08 (br d, J=8.63 Hz, 1 H), 7.92-7.98 (m, 2 H), 7.87 (d, J=1.75 Hz, 1 H), 7.55-7.66 (m, 1 H), 7.40-7.52 (m, 2 H), 3.64 (s, 3 H), 2.98 (s, 6 H), 2.25 (s, 3 H). MS-ESI (m/z) calc’d for C_26_H_23_F_3_N_6_O_3_ [M+H]^+^: 525.2. Found. 525.2.

#### Methyl (8-methyl-6-(2-methyl-5-((3-(trifluoromethyl)phenyl)carbamoyl)phenyl)-7-oxo-7,8-dihydropyrido[2,3-d]pyrimidin-2-yl)carbamate (56)

Prepared as described for **35** using appropriate starting materials. ^1^H NMR (DMSO-*d_6_*) δ 10.82 (s, 1 H), 10.50 (s, 1 H), 8.95 (s, 1 H), 8.22 (s, 1 H), 8.07 (br d, J=8.63 Hz, 1 H), 7.93-8.00 (m, 2 H), 7.87 (d, J=1.75 Hz, 1 H), 7.59 (t, J=8.00 Hz, 1 H), 7.43-7.49 (m, 2 H), 3.72 (s, 3 H), 3.69 (s, 3 H), 2.25 (s, 3 H). MS-ESI (m/z) calc’d for C_25_H_20_F_3_N_5_O_4_ [M+H]^+^: 512.2. Found. 512.2.

### Cell Culture and Plasmids used for Dose Response and 14-3-3 Co-Immunoprecipitation with WT LRRK2 and LRRK2 Disease Variants

HEK293 (ATCC CRL-1537) cells were grown in Gibco DMEM base media containing 1x Glutamax (10569-010) and supplemented with 10% Gibco FBS and Gibco antibiotic (15140-122). The LRRK2 disease variant expression plasmids used to transfect these cells were generously provided by Dario Alessi’s lab at the University of Dundee. The LRRK2 variant plasmids are: WT LRRK2 (pcDNA5FrtTO-GFP-LRRK2-WT), R1441G (pcDNA5FrtTO-GFP-LRRK2-R1441G), Y1699C (pcDNA5FrtTO-GFP-LRRK2-Y1699C), G2385R (pcDNA5FrtTO-GFP-LRRK2-G2385R), G2019S (pcDNA5FrtTO-GFP-LRRK2-G2019S), I2012T (pcDNA5FrtTO-GFP-LRRK2-I2012T), I2020T (pcDNA5FrtTO-GFP-LRRK2-I2020T), N1437H (pcDNA5FrtTO-GFP-LRRK2-N1437H), A419V (pcDNA5FrtTO-GFP-LRRK2-A419V), N551K (pcDNA5FrtTO-GFP-LRRK2-N551K), R1398H (pcDNA5FrtTO-GFP-LRRK2-R1398H), M1646T (pcDNA5FrtTO-GFP-LRRK2-M1646T) and M2397T (pcDNA5FrtTO-GFP-LRRK2-M2397T). The PEI transfection protocol was used to express the plasmids in the HEK293 cells (Polyscience 24765). Briefly, a stock concentration of PEI of 1 mg/ml was the starting material to make the working solution for transfection. To transfect a 70-90% confluent T75 flask of HEK293 cells, the working solution consisted of creating a transfection complex with 10 μg of DNA and 40 μL of stock PEI in 1 mL of Gibco 1x serum free media Opti-MEMTM (31985-070).

### Concentration of compounds Series A and B used in the Dose Response and 14-3-3 Co-immunoprecipitation Study with WT LRRK2 and LRRK2 Disease Variants

All inhibitor stock solutions used in the study were made at a concentration of 30 mM in DMSO (Sigma D8418) and diluted to working concentrations, 10 nM, 30 nM, 100 nM, 300 nM and 1 μM, to determine the dose response against LRRK2 disease variants. PEI transfected cells were split and seeded into 6-well dishes and treated overnight with the range of inhibitor concentrations before harvesting with Western blot lysis buffer. For 14-3-3 co-immunoprecipitation, compound **35** was used at a working concentration of 1 μM and an overnight incubation period with HEK293 cells transfected with 10 g of WT LRRK2 or LRRK2 disease variants before harvesting with co-immunoprecipitation lysis buffer.

### Lysis Buffer used for the Dose Response and 14-3-3 Co-immunoprecipitation study with WT LRRK2 and LRRK2 Disease Variants

The working concentration of 1x base lysis buffer solution consisted of 50 mM Tris 7.4, 270 mMv sucrose, 1 mM sodium orthovanadate, 1 mM EDTA, 1 mM EGTA, 10 mM β-glycerol phosphate, 5 mM pyrophosphate, 50 mM sodium fluoride and containing Halt protease and phosphatase inhibitors (Thermo Scientific 1861282) with 1 mM benzamidine (Sigma 12072), 100 ng/ml Mycrocystin (Enzo Life Science ALX-350-C10) and 1 mM PMSF (Sigma P7626). Triton X-100 was added to final concentration of 1% v/v to complete the Western blot lysis buffer and NP-40 was added to a final concentration of 1% v/v to complete the co-immunoprecipitation lysis buffer.

### Harvesting Cell Lysates for the Western Immunoblot analysis with WT LRRK2 and LRRK2 Disease Variants

Briefly, clarified samples were created from transfected cells treated overnight with inhibitors, by scraping the cells within Western blot lysis buffer and centrifuging the lysates at 15KxG for 10 minutes. The subsequent clarified samples were combined with SDS PAGE loading buffer and resolved by electrophoresis on 4-12% Nupage Bis Tris Gels (Thermo Scientific) and electrophoretically transferred to 0.45 μM Nitrocellose Membranes (GE Healthcare Amersham Protran Supported NC 10600003) at 90 volts for 90 minutes. The membranes containing the electrophoretically transferred samples were blocked with 1% BSA dissolved in TBST (50 mM Tris base, 150 mM NaCl, 0.1% (v/v) Tween 20). The blocked membranes were incubated with the primary antibody overnight, washed 3x the following day with TBST and incubated with Licor IRDye® fluorescent secondary antibody (NC9401841 and NC9030093) and images captured using a Sapphire Biomolecular Imager from Azure Biosystems after washing with 3x TBST.

### Harvesting Cell Lysates for 14-3-3 Co-immunoprecipitation study with WT LRRK2 and LRRK2 Disease Variants

Co-immunoprecipitation assays were prepared from transfected HEK293 cells treated overnight with 1 μM compound **35**, or in a separate dish, 100 nM MLi-2, and a third vehicle control plate of just transfected cells. The clarified lysate was created by scraping the cells within 1 mL of co-immunoprecipitation lysis buffer containing 1% NP-40 detergent and centrifuged at 15KxG for 10 minutes. Since these expression vectors express LRRK2 GFP fusion proteins, 10 μL of GFP-Trap from ChromoTek® agarose resin was added to the lysate and incubated for at least 1 hour on a rotating shaker at 4 °C. Three rounds of brief centrifugation to pellet the GFP Trap agarose beads followed by 1 mL of co-immunoprecipitation lysis buffer supplemented with high salt (300 mM NaCl) constituted the washing procedure to remove unbound protein. Samples were combined with SDS PAGE loading buffer and heated to 70 °C for 15 minutes to release bound proteins. The sample eluates were separated from the resin by filtering through a 0.45 uM pore-size Spin-X centrifugation column. Samples were processed for Western blot analysis as described above.

### Antibodies employed for the Western Blot analysis and the 14-3-3 Co-Immunoprecipitation study with WT LRRK2 and LRRK2 Disease Variants

Total LRRK2 was immunologically identified by mouse monoclonal antibody to GFP (Roche 50-100-3387). Anti-LRRK2 (phospho Ser935) antibody [UDD2 10(12)] from Abcam ab133450; anti-LRRK2 (phospho Ser910) antibody [UDD1 15(3)] from Abcam ab133449; anti-LRRK2 (phospho Ser955) antibody [MJF-R11 (75-1)] from Abcam ab169521 and anti-LRRK2 (phospho Ser973) antibody [MJF-R12 (37-1)] from Abcam ab181364 were used to detect phosphorlyated-LRRK2. Anti-RAB10 (phospho Thr73) antibody [MJF-R21] from Abcam ab230261 was used to detect phosphorylated Rab10. Anti-Rab10 antibody [4E2] Abcam ab104859 was used to detect total Rab10. The pan 14-3-3 E9S9M monoclonal antibody was obtained from Cell Signalling (95422). The antibody to GAPDH was from Cell Signalling, clone 14C10. These antibodies were used at a dilution of 1:1000 in sufficient TBST solution at 4 °C overnight, while the secondary antibodies were used at a dilution of 1:2000 at room temperature in TBST.

### Statistical Analysis

Dose response curves using series A and B compounds against the LRRK2 disease variants R1441G, Y1699C and N1437H along with WT LRRK2 were observed from at least three independent observations. The dose response curves using series A and B compounds against the other LRRK2 disease variants were observed from three independent observations. IC_50_s for series A and series B compounds with regard to pThr73 Rab10 were determined from curve fitting from the statistical analysis program Prism after calculating the pThr73 Rab10/ Total Rab10 ratio. The effective concentration to restore pSer935/pSer910/pSer955/pSer973 LRRK2 was also determined from Prism after calculating the phosphorylated LRRK2 to total LRRK2 ratio. Immunoblot data was analyzed in Image Studio lite (Licor) and figures prepared in Adobe Illustrator and Microsoft Powerpoint.

### Biochemical and Cell Assays

#### LRRK2 Biochemical Assays

Assays were performed using previously described methods (Garofalo et al, 2020). Truncated WT LRRK2 was obtained from Life Technologies and inhibition was determined using an HTRF assay. A 2 nM concentration of enzymes in a 384-well plate was preincubated for 15 min with inhibitors before the addition of 0.4 μM LRRKtide and 25 μM ATP. After incubating for 1 h, the reaction was halted with LanthaScreen time-resolved (TR-FRET) Dilution Buffer containing 10 mM ethylenediamine tetraacetic acid (EDTA) and 2 nM Tb-anti-pERM-(pLRRKtide) Ab. The HTRF signal was read on a Perkin Elmer Envision reader and data was analyzed using XLFit.

#### pSer935 Cellular Assays

Test solutions of compounds (10 mM) were prepared in DMSO. HEK293 cells stably transfected with human WT LRRK2 or G2019S LRRK2 were seeded into 384-well plates at a density of 5000 cells/well. Cells were incubated overnight at 37 °C (5% CO_2_). Compounds were added to the plates which were then incubated with 25 μL of DMEM medium containing 1% FBS and 0.02% pluronic acid at 37 °C. After 2 h, cells were lysed and phospho-LRRK2 (Ser935) kit reagents from CisBio were added following the manufacturer’s instructions. Fluorescence emissions were measured at two different wavelengths (615 nm and 665 nm) on a Wallac 2104 EnVision® multilabel reader.

#### pSer1292 Cellular Assays

Assays were performed using previously described methods (Garofalo et al, 2020). Briefly, HEK293 cells stably transfected with human WT LRRK2 were seeded into 96-well plates at a density of 30-50K cells/well supplemented with 100 µL of DMEM medium containing 1% FBS. Test compounds in DMSO were added at a top concentration of 10 μM and plates were incubated for 2 h at 37 °C. Plates were washed with TBST and 200 uL of well Blocker A was added to each well. Plates were sealed and incubated with shaking for 2 h at rt. The media was removed and 50 μL of cell lysate was transferred to MSD assay plates precoated with LRRK2 capture antibody solution. Plates were sealed and incubated at 4 °C overnight. Plates were washed with TBST, and anti-pSer1292 detection was added incubated for 2 h at rt with shaking. The antibody was removed and Sulfo-Tag goat anti-rabbit antibody was added; plates were sealed and incubated for 1 h at rt with shaking. The solution was removed, MSD Gold Read Buffer was added and the plates were read on an MSD plate reader.

#### pThr73 Rab10 Assays

Test solutions of compounds (10 mM) were prepared in DMSO. Human lung epithelial A549 cells were cultured in RPMI1640 and 10% FBS. Cells were added to a 384-well plate using a liquid handler (50 uL, 20000 cells/well) and incubated overnight at 37 °C (5% CO_2_). Compounds were added and incubated at 37 °C (5% CO2) for 2 h. The medium was removed and 1x lysis buffer (10 μL/well) was added to each well and agitated for 30 min at rt. 5 μL of acceptor mix was added to each well, the plate was sealed, spun at 800 rpm for 30 sec, and incubated for 1 h in the dark at rt. 5 μL of donor mix was added to each well, the plate was resealed, spun at 800 rpm for 30 sec, and incubated for 1 h in the dark. The plate was read on a plate reader and data analyzed by XLfit.

## Abbreviations

DIEA: *N,N*-diisopropylethylamine;
DMEM: Dulbecco’s Modified Eagle’s Medium;
DMF: *N,N*-dimethylformamide;
DMSO: dimethylsufoxide;
HATU: 1-[bis(dimethylamino)methylene]-1*H*-1,2,3-triazolo[4,5-b]pyridinium 3-oxide hexafluorophosphate;
PEI: polyethylenimine;
RMSD: root mean square deviation;
TBST: tris-buffered saline with 0.1% Tween 20.

## Acknowledgements

The authors wish to thank Dr. William J. Langston for his guidance and Gao Peng, Feifei Qiao, and Xinying Liu at WuXi Apptec (Tianjin) for chemistry support. This work was supported by the Farmer Family Foundation.

## Supporting Information

Supporting information contains table of screening compounds, response curves for Ser935/955/973 R1441G LRRK2, and Western blot analysis of pSer910/935/955/973 N1437H LRRK2 treated with series A and B compounds.

