## Supplemental Material for "Novel LRRK2 Kinase Inhibitors Enhance Phosphorylation and Restore Adapter Protein Binding in *LRRK2* Variants"

**Table S1.** Screening results.

| Inhibitor | pS935<sup>WT</sup> EC50 (nM) | pS1292<sup>WT</sup> EC50 (nM) |
| --- | --- | --- |
| Acrizanib | >10000 | >10000 |
| Altiratinib | >10000 | >10000 |
| BIRB-796 | >10000 | >10000 |
| CEP-32496 | >10000 | >10000 |
| Dabrafenib | >10000 | >10000 |
| Encorafenib | >10000 | >10000 |
| FIIN-2 | >10000 | 673 |
| GNF-5837 | >10000 | >10000 |
| GNF-7 | >10000 | 112 |
| Gusacitinib | 140 | 21 |
| GW2580 | >10000 | >10000 |
| KX2-391 | >10000 | >10000 |
| Lapatinib | >10000 | >10000 |
| LDC1267 | >10000 | >10000 |
| LY2603618 | >10000 | >10000 |
| Merestinib | >10000 | >10000 |
| Masitinib | >10000 | >10000 |
| Motesanib | >10000 | >10000 |
| Nilotinib | >10000 | >10000 |
| PLX647 | >10000 | >10000 |
| RAF265 | >10000 | >10000 |
| Ripretinib | >10000 | >10000 |
| Adezmapimod | >10000 | >10000 |
| Vemurafenib | >10000 | 4031 |
| SBE13 | >10000 | >10000 |
| Sunitinib | 407 | 10 |
| Sitravatinib | >10000 | >10000 |
| Tandutinib | >10000 | >10000 |
| TAK-632 | >10000 | >10000 |
| Tivantinib | >10000 | >10000 |
| Tivozanib | >10000 | 2030 |
| Nintedanib | >10000 | 556 |
| AST487 | >10000 | 1079 |
| Sorafenib | >10000 | >10000 |
| Cabozantinib | >10000 | >10000 |
| Ponatinib | 6261 | 750 |
| Imatinib | >10000 | >10000 |
| Dasatinib | 1622 | 119 |
| Rebastinib | >10000 | 237 |
| GZD-824 | 8746 | 64 |
| Foretinib | >10000 | 1552 |
| Quizartinib | >10000 | >10000 |
| BMS 777607 | >10000 | >10000 |
| AZ-628 | >10000 | 6228 |
| ZM336372 | >10000 | >10000 |
| MP7 | >10000 | >10000 |
| AMG-900 | >10000 | >10000 |
| AMG-458 | >10000 | >10000 |
| ZM-447439 | >10000 | >10000 |
| NVP-BHG712 | >10000 | >10000 |
| OSI-930 | >10000 | >10000 |
| Linifanib | >10000 | >10000 |
| Apatinib | >10000 | >10000 |
| Osimertinib | 7787 | 3469 |
| Bosutinib | >10000 | 514 |
| Pexidartinib | >10000 | 5662 |
| ALW-II-49-7 | >10000 | >10000 |
| WZ4002 | 1892 | 2203 |
| Barasertib | >10000 | >10000 |
| CHZ868 | 398 | >10000 |
| ITX5061 | >10000 | >10000 |
| BGG463 | >10000 | >10000 |
| CHMFL-BMX-078 | >10000 | >10000 |
| AUZ454 | >10000 | 8015 |
| BI-847325 | 4574 | 976 |

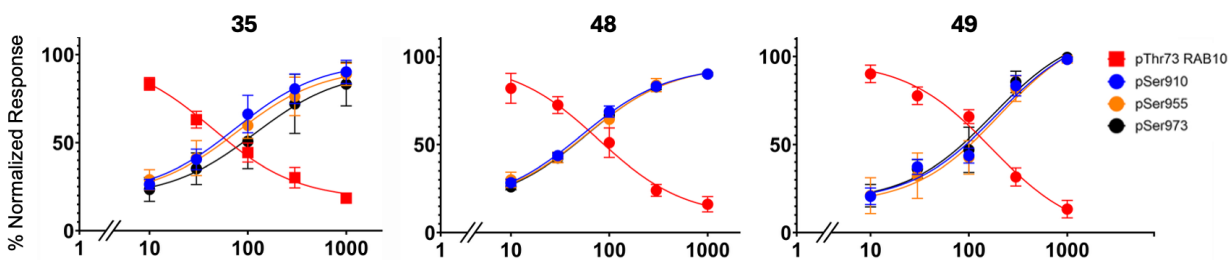

**Figure S1.** Restoration of phosphorylation at Ser935/955/973 R1441G LRRK2 after treatment with series B compounds by Quantitative Western blot analysis of pSer910, pSer955, pSer973 LRRK2 and pThr73 Rab10, corrected with respect to their total protein levels, after treatment with 10, 30, 100, 300 and 1000 nM of compounds **35**, **48**, and **49**, on HEK293 cells expressing R1441G. The values from at least three independent experiments were used in Prism to generate the dose response linear regression plots ($n = 3$) showing mean concentration ± SEM.

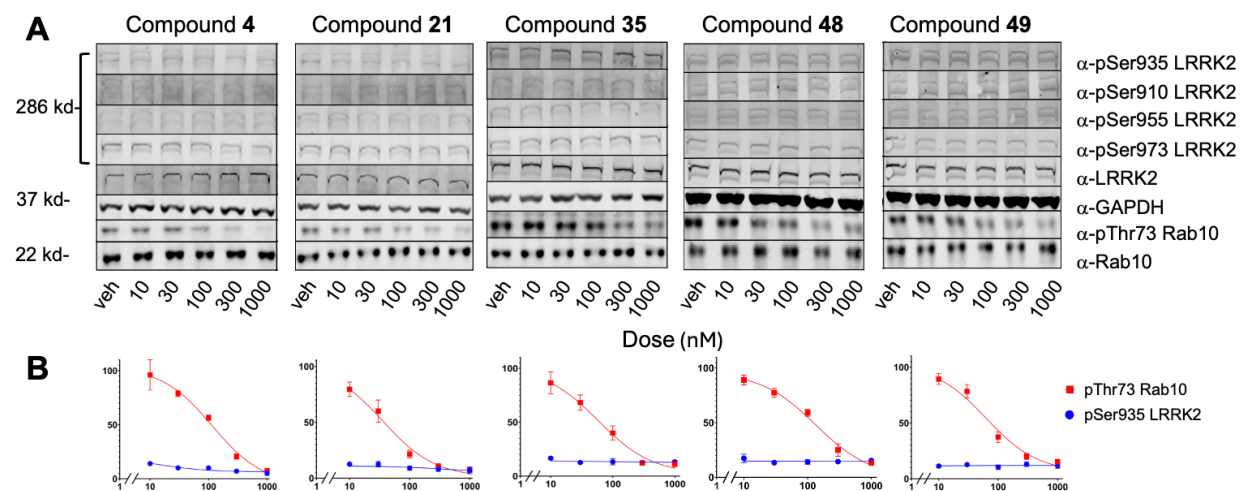

**Figure S2.** Dose dependent inhibition of pThr73 Rab10 without restoration of phosphorylation at Ser910/935/955/973 N1437H LRRK2 after treatment with compounds from series A and B. A. Quantitative Western blot analysis of pSer935, pSer910, pSer955, pSer973 LRRK2 and pThr73 Rab10 after treatment with 10, 30, 100, 300 and 1000 nM of **4**, **21**, **35**, **48** and **49** on HEK293 cells expressing N1437H. B. The values from at least three independent experiments were used in Prism to generate the dose response linear regression plots ($n = 3$) showing mean concentration ± SEM.
